# Meta-analysis of the pathogen *Leishmania donovani*’s transcriptome reveals multiple modes of regulation including two reciprocally regulated gene modules

**DOI:** 10.1101/2024.04.26.591251

**Authors:** Pradyumna Paranjape, P K Vinod, Tushar Vaidya

**Affiliations:** CSIR-Center for Cellular and Molecular Biology, Hyderabad, 500007, India; International Institute of Information Technology, Hyderabad, 500032, India

## Abstract

*Leishmania donovani* causes a neglected tropical disease called visceral leishmaniasis. Additionally, leishmaniasis also manifests opportunistically, under conditions of immune compromise. The continued non-availability of effectively curative interventions (drugs or vaccines) against this disease necessitates a deeper knowledge of *Leishmania* biology in order to evolve novel strategies against the disease. We have used a meta-analysis approach to analyse *Leishmania*’s composite genetic network rather than investigating individual candidate genes. We performed Weighted Gene Co-expression Network Analysis (WGCNA) on publicly available *Leishmania donovani* transcriptome data to identify co-regulatory genetic modules. This clustering of *Leishmania donovani* transcriptomes revealed that genes fall in 30 distinct co-regulated modules with 32 to 3012 genes. In order to analyse the distribution of genes in *Leishmania* gene modules, we queried the enrichment or depletion of various annotation-qualifiers in the modules. We observed that several modules are specifically and individually enriched or depleted for annotation qualifiers derived from GO-annotation and KEGG-pathways and are differentially associated with life-phases and experimental conditions. Additionally, modules are also enriched or depleted for sequence based genetic features such as chromosomal location, location on co-transcriptional segment, rank of transcript from initiation of transcription, skewed usage of known RNA Binding Protein motifs. Classification of uncharacterized transcripts into co-regulatory modules provides insights in their probable characteristics, aiding future empirical investigation. Strikingly, two of the modules have reciprocal features including individual associations with logarithmic or stationary growth phases of *Leishmania*, two important life-phases that simulate the vector-dwelling pro-cyclic and the pre-infective meta-cyclic forms. Collectively, our analyses of *Leishmania* co-regulated gene modules is suggestive of additional regulatory modes over the mere differential mRNA stabilization.

## 1. Introduction

*Leishmania* causes visceral, cutaneous and muco-cutaneous leishmaniasis, affects populations in tropics, sub-tropics and southern Europe. Descriptions as early as 7^th^ century^1^ BC report lesions indicative of leishmaniasis. Visceral leishmaniasis (VL), which is fatal if untreated, is caused by *Leishmania donovani* and *infantum*. World Health Organisation estimates annual occurrence of 50,000 to 90,000 new cases of VL, with poverty and malnutrition being some of the risk factors. Treatment is limited to non-specific drugs such as pentavalent antimonials and anti-fungals^2^, which come with risks of severe side-effects. Thearpy is further limited by evolution of drug resistant^3^ strains. Lack of effective vaccine poses an urgency for development of targeted drugs and improved disease intervention. Comprehensive knowledge of *Leishmania* biology, particularly the genetic framework has become essential for informed design of targeted drugs and vaccines.

Phenotypically observable changes in cells are brought through remodelling of the internal cellular environment, with activation and modulation of concentrations of biomolecules. Such biochemical states of the cell, observable as steady-stable relative concentrations of various biomolecules, are achieved and maintained by regulatory genetic circuits that together form the framework of a cellular genetic network. Gene-interactions and not mere presence of gene-products forms the backbone of cellular working. Whole genome sequence information of several *Leishmania* species have become available over the last decade. However, we are yet to have a comprehensive understanding of *Leishmania* genetics, which could be leveraged in to advances in therapy and prevention. While DNA (genome) forms the repository of genetic information, RNA (transcriptome) generally forms the first level of concentration-modulation. An integration of data accumulated from varied sources such as genome and transcriptome would help fill the completeness of our knowledge of *Leishmania* biology.

Several transcriptomes covering various life conditions, independently generated by multiple research group are available in public databases. RNA sequencing of complete transcriptome of a cellular condition presents a quantitative measure for its phenotypic description, enabling numerical analyses. Such analyses can help in identification of prospective genetic elements that may contribute to establishment and/or maintenance of biologically defined target phenotype in the organism. Differential expression analyses yield sets of transcripts, which are typically up-regulated or down-regulated in the target phenotype, compared to a control. This may be extended to identify sets of co-expressing transcripts.

Demonstration of co-localization has been the *litmus* test for interaction of gene-products. Since the gene products ought to exist together to co-localize, co-expression of their respective mRNA becomes imperative. Thus, classification of coexpressing genes becomes an initial metric to group similar genes. However, co-expressing transcripts are insufficient in describing the regulatory context and interacting partners of a given gene, since genes with expression profiles tracing characteristics reciprocal to that gene are discounted. Such antagonistic transcripts may be counter-partners of (repressors, repressed by) the given gene, important for its characterization. Co-expressing transcripts are expected to be regulated by shared upstream elements. Differential expression profiles of unknown phenotypes may also cluster with known phenotypes, for descriptive qualification.

Exploration of functions of gene-products in order to establish interactions and successive fulfillment of lacunae of the common knowledge-base has been the approach towards understanding cellular-biology. Qualification of genes by their classification using co-regulatory network analysis, on the other hand, focuses on signal processing through gene-networks rather than the particular bio-chemical mechanisms to carry out the intended functions. Individual studies describe test conditions chosen by research-groups, yielding lists of genes which are differentially expressed under those conditions, compared against a commonly accepted arbitrary control. Ensemble effects of interplay between various modules of genetic regulation are lost in such studies. Description through robust physico-chemical interactions of candidate genes, hence becomes imperative. A meta analysis of gene expression information across multiple such studies allows for establishment of correlation between the behaviour of global sets of transcripts. Such cumulative analyses of transcriptomes aids in reconstruction of potential genetic-regulatory networks. Input data about transcript concentrations across diverse biological conditions, required for statistical reconstruction of such genetic-network, are available for *Leishmania donovani*.

Regulation of gene-expression in *Leishmania* is distinct from classical eukaryotic regulation. mRNA genes form extensive poly-cistronic nascent transcripts emerging from chromosomes^4^. The intergenic regions on these nascent transcript, upon post transcriptional cleavage, form 3’UTR of the upstream gene and 5’ flank of the downstream gene. Cleaved individual mRNA transcripts are trans-spliced on the 5’ flank to a common Spliced Leader (SL) and poly-adenylated beyond the 3’ UTR to form the mature RNA. With such non-canonical mechanism and lack of known promoters and introns, transcriptional regulation is attributed to post-transcriptional mRNA stabilization or destabilization by RNA-Binding Proteins (RBP). Hence, separate investigation of *Leishmania* gene regulation becomes essential rather than an assumptive extrapolation from known eukaryotic mechanisms. Also, this limited scope of mRNA regulation either, reduces our investigative domain to attribute the cause to RNA-Binding Proteins, or persuasively indicates existence of mechanisms hitherto unknown.

WGCNA is tool that may overcome this deficiency by directly generating gene-network models through meta-analysis of gene-expression to classify genes into co-regulatory modules. We performed WGCNA with the available transcriptomes to cluster *Leishmania donovani* genes into co-regulatory modules to explore its genetic regulation.

## 2. Materials and methods

### 2.1 *Leishmania donovani* reference genome

The predicted genes of RefSeq genome for *Leishmania donovani* BPK282A1^5^ do not include information about UTRs, and only identify the CDS regions. Read-counts against such genome annotation would not correctly represent actual gene-concentrations reported in RNASeq datasets^6^. To get a more accurate readcount, LdHU3^7^ annotation was used instead. Genes are delineated in LdHU3 annotation based on evidence from RNASeq read mappings, providing more accurate extent of their lengths and UTRs, which may contain gene-regulatory information.

### 2.2 Transcriptome Datasets

Raw reads from publicly available *Leishmania donovani* transcriptome datasets (supplementary document Table 1) including classical RNASeq and SL-Seq^8^, with single and paired-end reads were retrieved from NCBI, trimmed using cut-adapt^9^, checked for quality-control using FastQC^10^ and suitably mapped to reference genome *Leishmania donovani* HU3 using HISAT2^11^ without splicing. Reads were processed through standard read quality check using FastQC and trimmed using cut-adapt. Spliced-leaders from SL-seq reads were trimmed for the leader sequence. Appropriate reads were mapped to LdHU3 genome using HISAT2 without splicing and were counted for alignment to LdHU3 transcript features corresponding to mRNA genes. Fraction of reads mapping to *Leishmania* (using HISAT2) was found to be <3% for reads with contaminant reads of host cells. To test if contaminant reads are included in specific read-mapping, we checked that *Mus musculus* reads fraction found in *Leishmania* read mappings were not significantly more for contaminant-containg samples as compared to axenic samples. Fourteen of 57 experimental runs which did not pass quality check(s) were eliminated from further analysis, leaving 43 experimental runs across 10 distinct conditions. [Supplementary table 1] This pipeline from retrieval, quality-check, mapping and batch-adjustment was coded in a python package that can be bootstrapped with accession numbers, known batches and phases to yield adjusted read counts. https://gitlab.com/pradyparanjpe/multirnaseq.git Mapping was confirmed for quality check using multiQC^**?**^ and TPM counts were obtained using TPMCalculator^12^. Supplementary Table 1 summarizes the input datasets and corresponding accession. Batch-effects were adjusted using R-packages limma::vst^13^ and ComBat^14^.

**Table 1.**
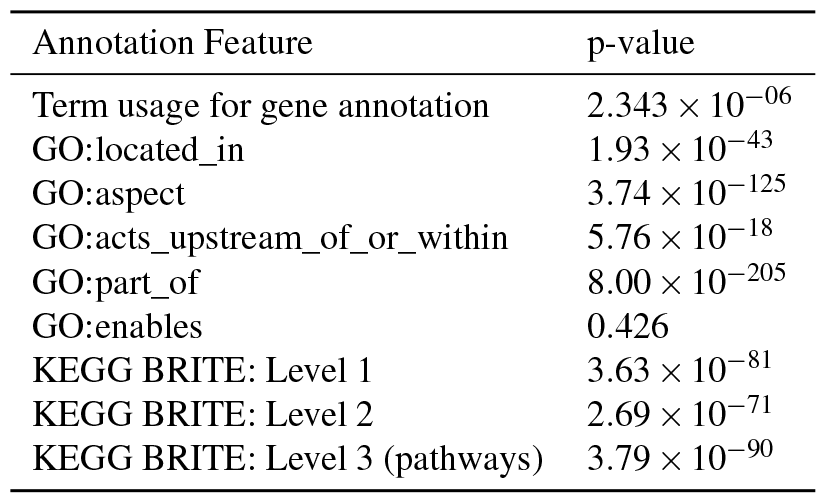
Association between WGCNA clustering and annotation features.

| Annotation Feature | p-value |
| --- | --- |
| Term usage for gene annotation | $2.343 \times 10^{-06}$ |
| GO:located_in | $1.93 \times 10^{-43}$ |
| GO:aspect | $3.74 \times 10^{-125}$ |
| GO:acts_upstream_of_or_within | $5.76 \times 10^{-18}$ |
| GO:part_of | $8.00 \times 10^{-205}$ |
| GO:enables | 0.426 |
| KEGG BRITE: Level 1 | $3.63 \times 10^{-81}$ |
| KEGG BRITE: Level 2 | $2.69 \times 10^{-71}$ |
| KEGG BRITE: Level 3 (pathways) | $3.79 \times 10^{-90}$ |

**Table 2.**
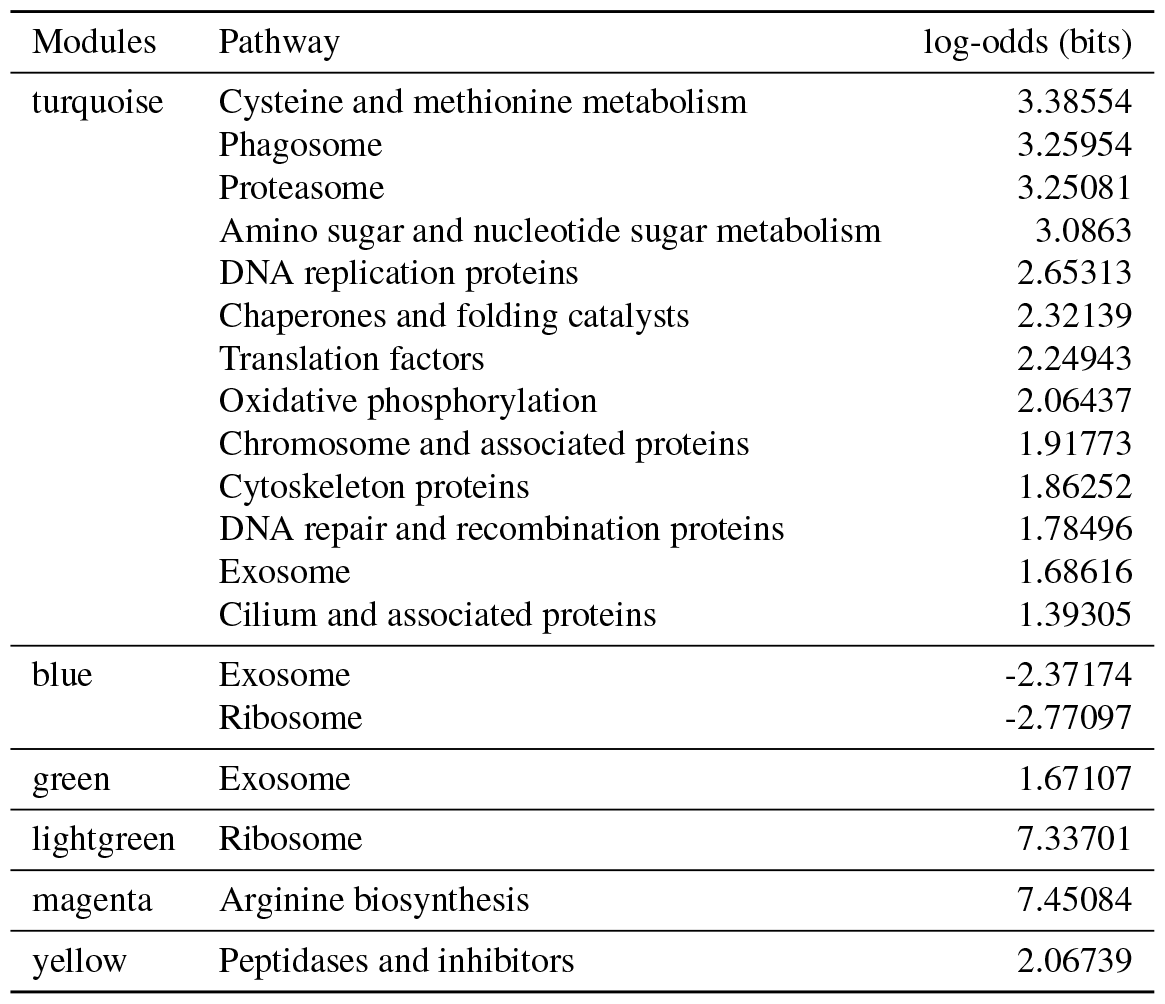
Association of KEGG Pathways with WGCNA modules. ‘Log-odds’ column indicates the extent of enrichment (negative for depletion) of genes belong to the corresponding pathway in the indicated module. (p-value < 0.001)

**Table 3.**
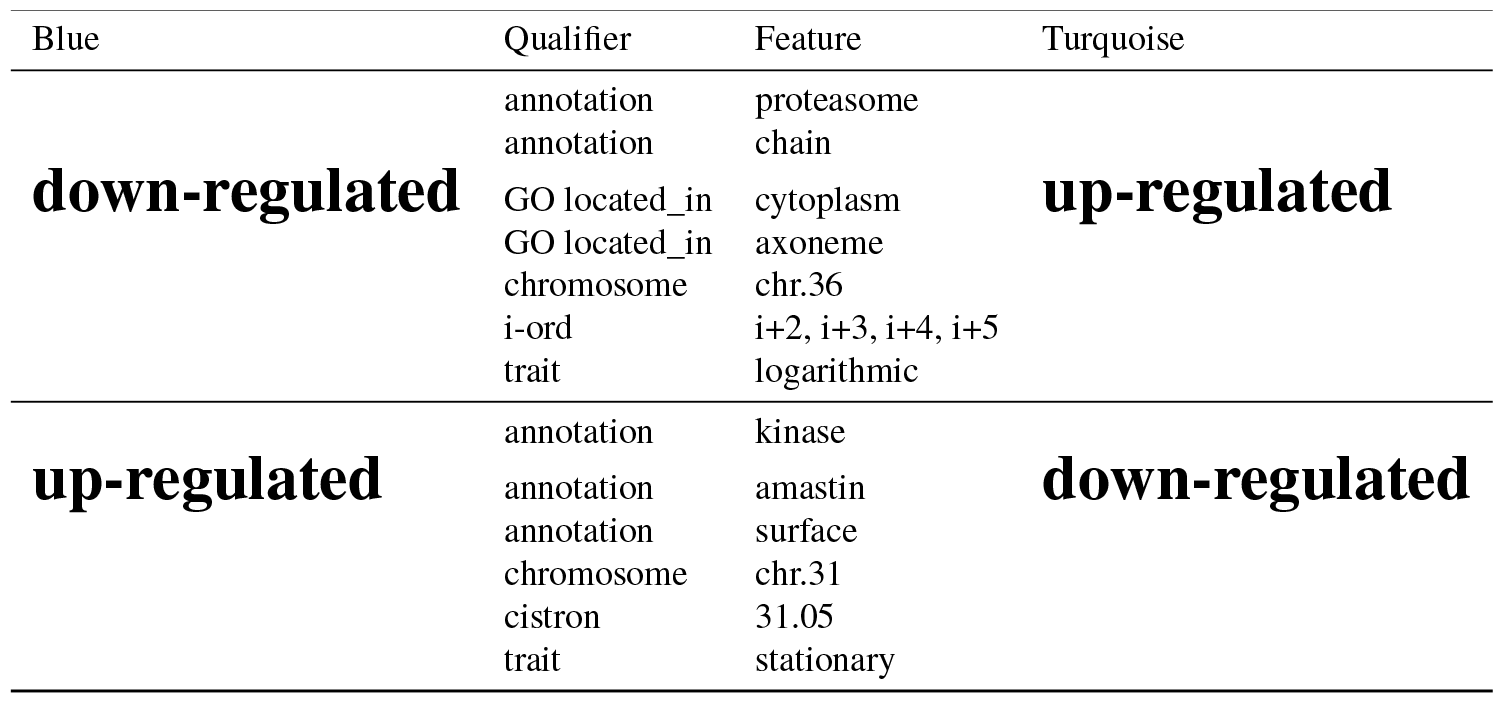
Reciprocal enrichment of turquoise/blue features.

### 2.3 WGCNA

WGCNA was performed as prescribed^15^, with parameters optimized to maximize the number of modules. Final parameters were as follows, Soft Threshold Power, 10; Minimum number of transcripts to call a Module, 30; network-type, co-regulation; Module split, 4; Modules re-merging, not permitted.

### 2.4 Poly-cistrons

Boundaries of poly-cistrons were located based on reversal of transcript-orientation. Largest contiguous chromosomal span of similarly oriented transcripts was marked as a ‘polycistron’ feature in the gtf-file. (supplementary file: mRNA_polycistrons.gtf).

### 2.5 Statistical analysis

Contingency tables for each examined genetic feature were generated to collate the occurrence of each candidate feature across the modules (supplementary data). Inter-dependence of each genetic feature with module-clusters was established using chisquared test, p-value < 0.001 considered significant. Fisher’s exact test with p-value < 0.001 was considered significant for enrichment of individual terms in modules, log-odds (bits) are plotted. Python scripts generated to extract this information about enrichment of genetic features are available at https://gitlab.com/pradyparanjpe/wgcna_extract.git.

## 3. Results

In order to identify modules of co-regulated genes, we performed clustering of publicly available *Leishmania donovani* transcriptomes.

### 3.1 Batch effects normalization

We initiated our analysis by retrieving publicly available single and paired end reads RNA-seq and SL-seq raw reads for *Leishamina donovani* from NCBI database. These transcriptome counts across experimental runs suffer from variance on account of batch effects. Any batch-effects in read-counts were adjusted using the R package limma::vst and eliminated using ComBat. Experiments SRR5272501, SRR5272504, SRR5272507, which were found to skew counts with batch effects, were dropped from further analysis. Finally, we proceeded with 40 experimental runs covering 10 distinct conditions for further analysis. In contrast to unadjusted counts, where experiments cluster by their batches, similarly described biological conditions appear nearer on a 2D PCA plot on combat-adjustment, confirming removal of batch-effects.

### 3.2 WGCNA

Combat-corrected read-counts were used to perform WGCNA using standard described protocol. WGCNA parameters were systematically tested with a goal to optimize the number of co-regulatory clusters. Thirty regulatory network clusters (and a set of unassigned genes) were identified after WGCNA, with cluster sizes ranging from a minimum of 32 to a maximum of 3013 genes. We confirmed optimal clustering by establishing no further clustering of the two largest clusters. WGCNA assigns color-names as identifiers of clusters. Color-hue similarity (if any) is completely incidental and does not indicate any feature of the module. We continue to use color-names assigned by WGCNA as identifiers of regulatory modules in all further analysis.

### 3.3 WGCNA clustering scheme and known genetic annotation features

We tested the association between WGCNA-clustering and various annotation features using the χ^2^ test (table 1). There is significant association between WGCNA clustering and annotation features such as gene-annotation terms, Gene Ontology (GO) qualifiers and KEGG-BRITE classifications. This indicates that the WGCNA clusters are consistent with biologically known schemes of classification.

#### 3.3.1 KEGG Pathways

We validated the learnt WGCNA modules by examining the non-random association of modules with common annotation terms, GO-qualifier terms (Supplementary document 1) and KEGG Pathways.

We expect genes involved in common metabolic {domains,fields} to be co-regulated for their functioning in unison. KEGG-Pathways is a database which lists common metabolic pathways for reference organisms. Since LdHU3 genome was assembled on *Leishmania major*, an organism whose databases are comparatively much better curated, we used *L major* orthologs of LdHU3 genes to analyse pathway-enrichment. We extracted orthologous *Leishmania major* gene-identifiers for LdHU3 transcript identifiers from the LdHU3 features file to identify pathways in which they are involved.

KEGG’s BRITE functional hierarchies provide hierarchical classification of gene-function. The first three levels progressively explicate the gene’s classification to pathways while the final level describes the gene. For each of the first three levels of BRITE hierarchical classification, we generated contingency table of occurrence of LdHU3 genes in WGCNA- modules and BRITE identifiers at that hierarchical level. Genes’ classification into WGCNA clusters is associated with each of the three levels of BRITE Hierarchical classification (χ^2^ tests, p-values: Level-1, 3.63e-81; Level-2, 2.69e-71; Level-3, 3.79e-90).

We further checked association between occurrence of genes in WGCNA modules and the pathway classified by the third level of BRITE hierarchies using *Fisher’s exact* test, (*p*−*value <* 10^−3^).

Table (2) lists identities of pathways enriched or depleted (if any) for each module and log-odds of their occurrence over random-distribution. *Turquoise* is associated with most (13) pathways, six which, are involved with nucleic acids. In agreement with observations from annotation terms skew, *lightgreen* module shows association with Ribosome.

### 3.4 Working theory of Transcription

Due to the lack of identified canonical cis and trans acting transcriptional elements, (emperically or through sequence homology), the current model of mRNA regulation in *Leishmania* is decoupled from chromosomal features such as classical promoters and enhancers typically observed in most eukaryotes. Transcription of *Leishmania* genes occurs as nascent poly-cistrons of adjacent genes. Differential gene-regulation in *Leishmania* is attributed to differential RNA-stabilization (or de-stabilization) through RNA-Binding Proteins (RBP) owing to lack of known transcription-factors and introns.

In the present study, we set to investigate linkage between gene-regulation and genetic features. In our analyses, for every feature considered, Fisher’s exact test was used with *p*−*value <* 10^−3^ confidence to test if the feature is enriched or depleted in each of the WGCNA modules. In each of the tests, without contribution form an additional regulatory paradigm, genetic features are not expected to be linked to regulation, and consequently, not expected to be enriched or depleted beyond random occurrence in the co-regulatory modules.

#### 3.4.1 Motifs

While RBPs and their corresponding recognition motifs on the 3’-Untranslated regions of genes (3’UTRs) are poorly defined in *Leishmania*, we asked under assumption of evolutionary relatedness and conservation whether RBP-motifs defined in other Eukaryotes are also utilized in *Leishmania*. Probability Weight Matrices (PWMs) for all currently available [297] eukaryotic RBP-Motifs were retrieved from CISBP-RNA database. Probability Weight Matrices [PWM] were converted to regular expressions such that a base was included for match if its weight in the PWM was greater than the threshold of 0.2. Motifs were located in *Leishmania* transcripts matching regular expressions for each transcript. An un-used motif would carry no information for the RBP repertoire of *Leishmania* and hence, would be randomly distributed across the transcriptome. However, a motif that carries information relevant in *Leishmania* would resist evolutionary mutation, taking its occurrence further from randomness. Expected random occurrence of irrelevant motifs was calculated from the probability of finding the regular expression accounting for propensity of nucleotides in the complete transcriptome. Of the 297 motifs with PWMs identified by CISBP-RNA, 278 (93.9%) motifs have skewed occurrence in *Leishmania* transcriptome, enhancing their potential to be used by *Leishmania*.

Further, for each RBP-motif derived from CISBP-RNA database, we tested the enrichment of genes containing that RBP-motif in each WGCNA module, with expected occurrence derived from frequency of such genes in the complete transcriptome. A gene was considered to contain a given motif if regular expression describing that motif matches at least once in the gene’s 3’UTR. Any significant skew away from expect value would imply a non-random distribution, which would imply an module-specific usage or evasion of usage of that motif. We queried UTRs of genes from co-regulatory modules for known motifs as retrieved from the CISBP database. For the query, mere occurrence of motif was considered sufficient, position of motif in the UTR, number of copies of the motifs were ignored. A motif was considered to be enriched or depleted in a WGCNA module based on the odds ratio of the number of genes containing that motif classified under the module to the number of genes expected given the propensity of occurrence of the motif and the size of the module.

Figure (3) enumerates the number of motifs skewed in each WGCNA module.

**Figure 1.**
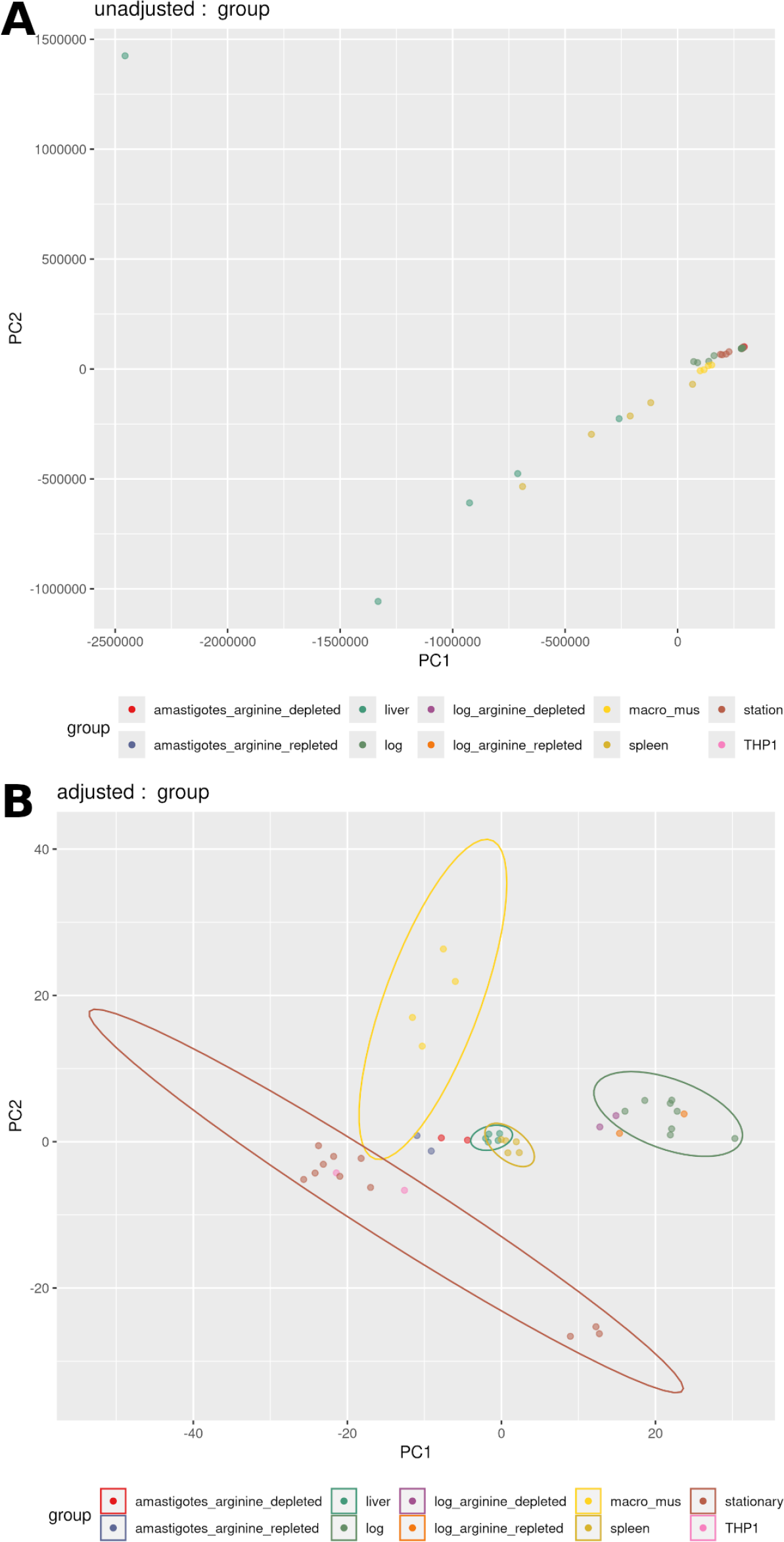
PCA of Experiments: **A**. Without batch-effects-correction. Colors indicate batches. **B**. After batch-effect-correction: Colors indicate conditions.

**Figure 2.**
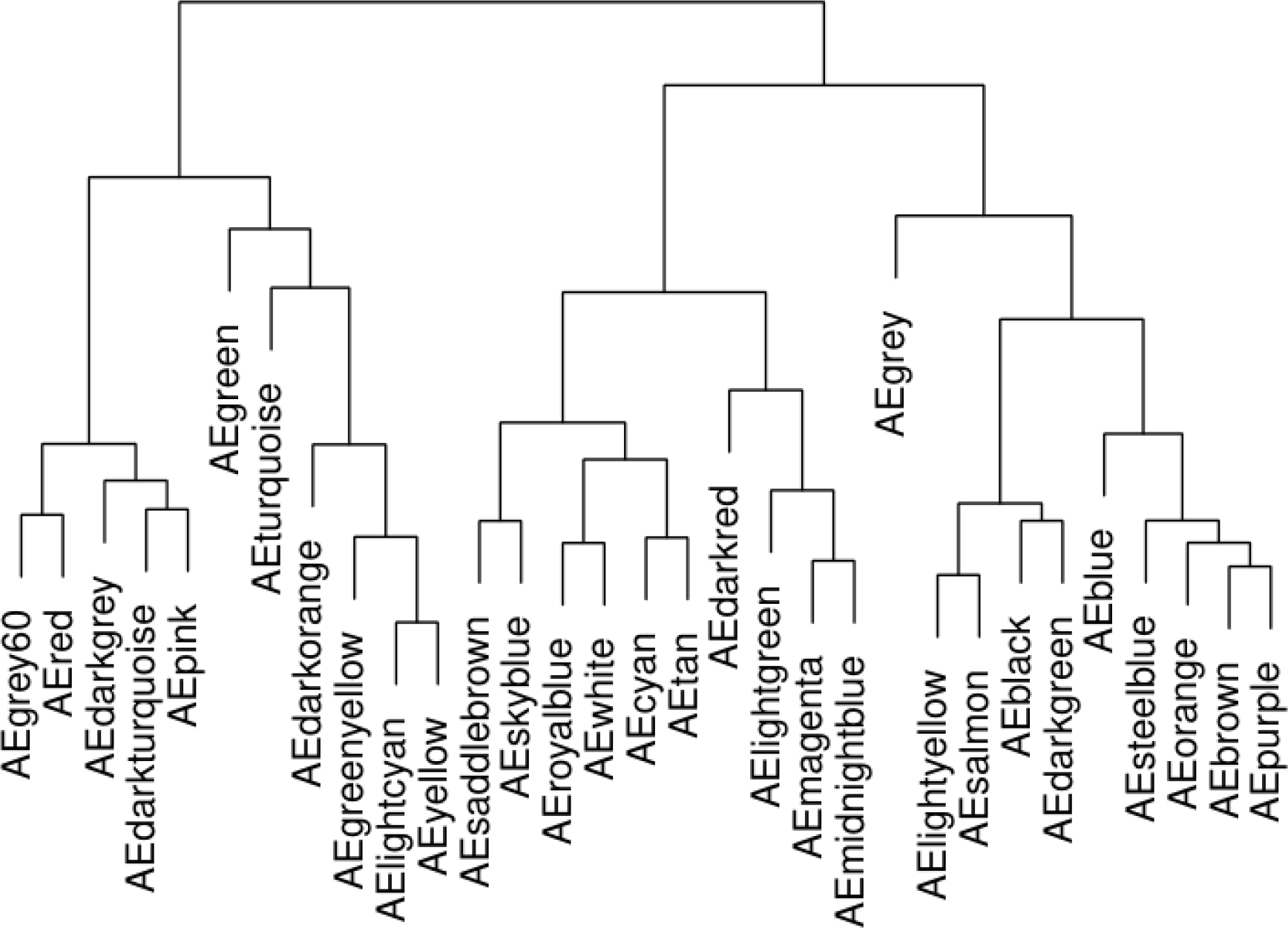
WGCNA modules: Colors indicative of distinct modules, *grey* is set of unclassified genes.

**Figure 3.**
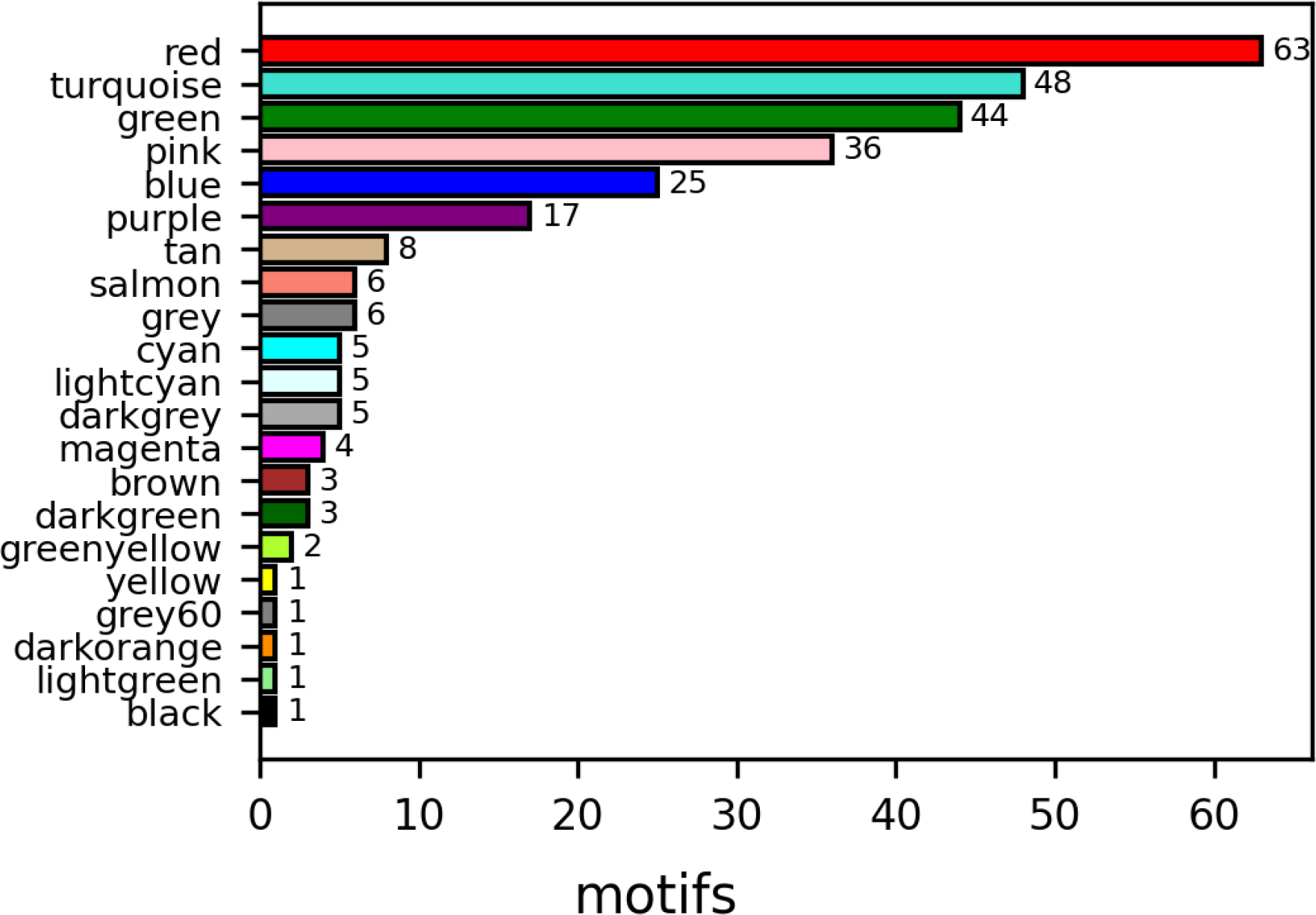
Number of motifs enriched/depleted in each module: A motif is considered *enriched* or *depleted* if genes containing that motif are enriched or depleted.

Forty-one motifs (M **N**) indicated in figure (4) are enriched while 22 are depleted in *red* (Fisher’s exact test, p-value < 10^-3^). RBP Motifs form the basis of regulation of genes under the current model of gene regulation in *Leishmania*. Thus, motifs enriched in a particular module indicate a common regulatory signal for a typical gene in that module. Supplementary table 5 enlist motifs which are enriched or depleted from modules.

**Figure 4.**
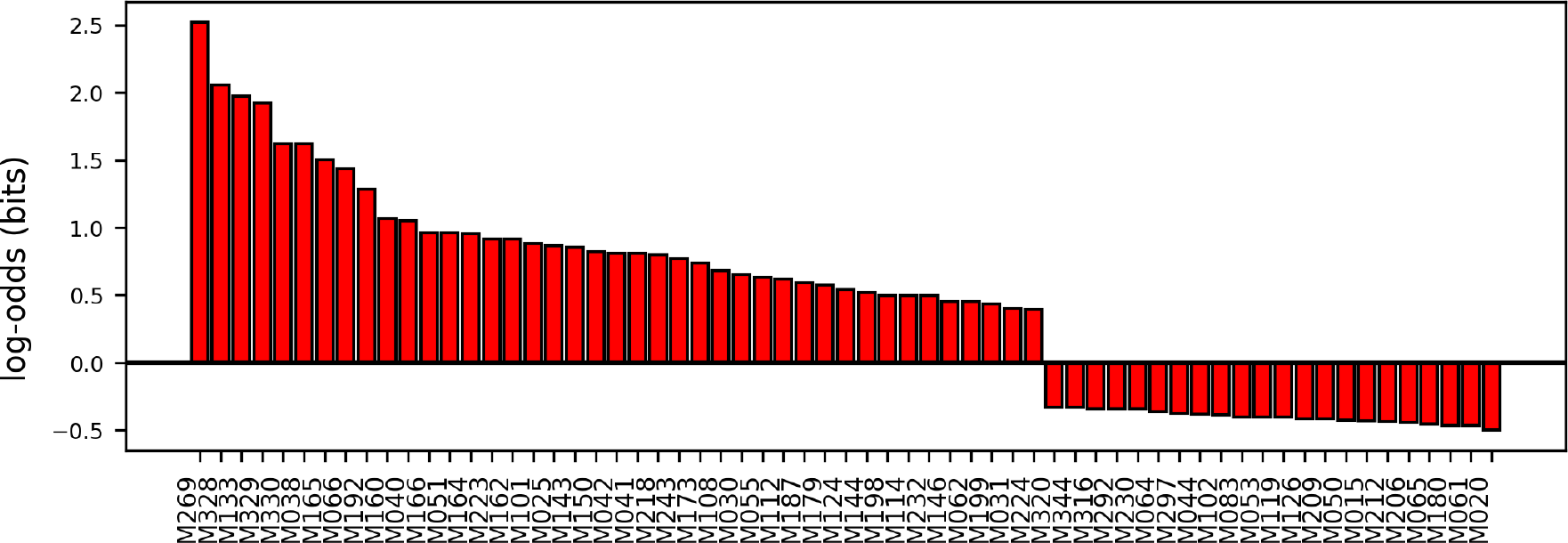
Motifs enriched / depleted in red module.

### 3.5 Association with genetic features

While informative annotation is available for only about 45% *Leishmania* genes, the *Leishmania* genome, which is a complete repository of encoded information used by *Leishmania*, is available in its entirety. Hence, we further investigated association of co-regulatory modules with genetic features for additional insights into *Leishmania*’s gene-regulation. For each feature of interest, we first checked if WGCNA-clustering scheme is associated with that feature using the χ^2^ test. Features with p-value < 0.001 were considered significantly associated with WGCNA-clustering scheme, suggesting their role in genetic regulation. These were further investigated for enrichment or depletion of each of their qualifiers in each of the WGCNA modules.

#### 3.5.1 Chromosome

The first genetic feature tested was the chromosome of origin of a given transcript. Chromosomal origin, as derived from the features file (GFF/GTF) and module-gene exports from WGCNA analysis were used to generate a contingency table, pearson’s χ^2^ test was performed on this contingency table. The observed χ 2 p-value (8 *×* 10^−40^) suggests contribution of chromosomal origin to classification of transcripts in specific WGCNA modules, and hence, involvement in genetic regulation.

Having established this, we further checked association of each chromosome-module pair using *Fisher’s exact* test (*p* − *value <* 10^−3^, same contingency table). Figure (5) represents on a logarithmic scale, odds-ratios of chromosomal enrichment (or depletion) across modules for all pairs with significant association. 11 of 36 chromosomes are associated with at least one regulatory module. In modules *cyan* (Chr. 1, 13), *pink* (Chr. 2, 12), *green* (Chr. 8, 17, 34), *magenta* (Chr. 10), *blue* (Chr. 31), *red* (Chr. 35) and *turquoise* (Chr. 36), we observe enrichment of genes encoded on particular chromosomes. *Turquoise* (Chr. 31) and *blue* (Chr. 35, 36) show depletion, indicative of regulatory *avoidance*. Not every chromosome has genes enriched or depleted in some WGCNA module. Clearly, not all genes from any given chromosome cluster into a particular module, neither does any module derive all its genes from a single chromosome, indicating an obvious lack of exclusive regulatory authority in chromosomes. *Turquoise* and *blue* modules are both enriched and depleted for genes originating from particular chromosomes. These analyses suggest contribution of chromosomal co-occurrence of genes with their co-regulation, which goes beyond the RBP’s scope of gene regulation.

**Figure 5.**
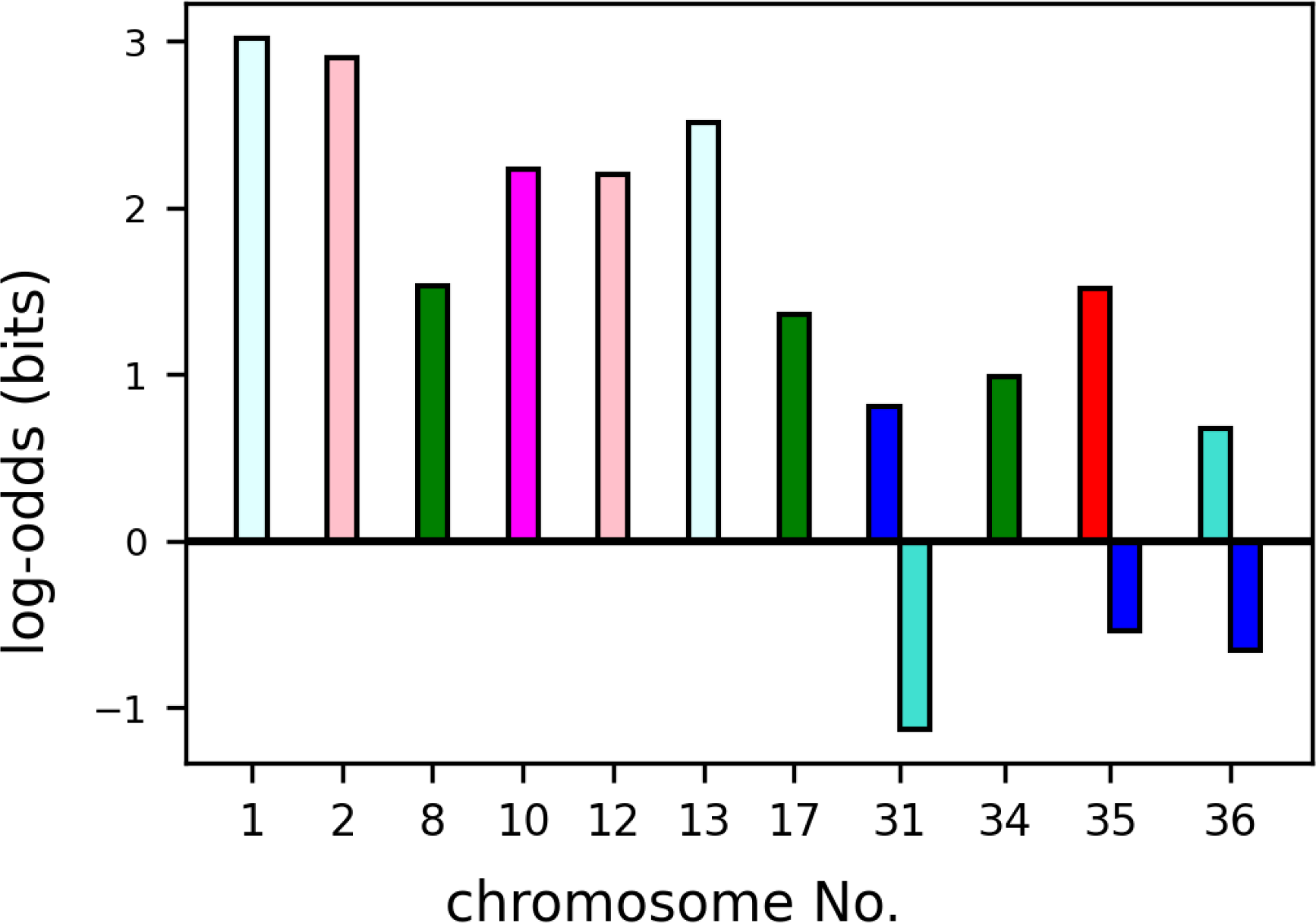
Enrichment of chromosomal origins. Color of bar indicates module. [*p*−*value <* 10^−3^]

#### 3.5.2 Poly-Cistron

Current understanding of *Leishmania* genetics suggests that genes are typically co-transcribed as poly-cistronic transcripts, which are later cleaved, poly-adenylated and capped into individual mature mRNA segments, which are then regulated by differential stabilization through RNA Binding Proteins (RBPs). Given our unexpected findings associating chromosomal origins of *Leishmania* genes with their co-regulatory modules, we examined if co-transcription would also be involved in co-regulation. Poly-cistrons were delineated as stretches of adjacent genes annotated to be transcribed in the same direction. Supplimentary file (Leishmania_donovani_HU3_mRNA_polycistrons.gtf) is an edited gtf features file and includes these polycistrons with an additional attribute ‘gene_count’ that enumerates the number of genes in that poly-cistron. A minimum of 2 (in Chr. 1, 2, 4, 17, 18, 19, 26, 31) to a maximum of 7 (Chr. 35, 36) poly-cistrons were identified for each chromosome. We tested (χ^2^ test) inter-dependence of poly-cistronic origin and WGCNA-module classification for genes from their contingency table. The poly-cistronic origin of genes is shows association with WGCNA classification (*p*−*value/*1.64 *×* 10^−19^).

We took this analysis further by exploring the association of each poly-cistron-module pair. Figure (6) represents on logarithmic scale, the odds-ratio of enrichment (or depletion) of genes from particular poly-cistrons across specific modules. These poly-cistron features were numbered sequentially in the format <CC.PP> in the direction of sequencing for each chromosome, where, ‘CC’ is the chromosome number, ‘PP’ is the poly-cistron number. For example, the first poly-cistron (in the direction of sequence assembly) on chromosome 29 is numbered *29.01. Black* (29.01), *blue* (31.01), *brown* (35.07), *green* (35.07), *greenyellow* (32.01), *pink* (12.03, 21.02), *red* (35.06), *salmon* (26.02) and *turquoise* (36.01, 36.05) modules are enriched with genes derived poly-cistrons respective poly-cistrons indicated in parentheses. At the same time, *turquoise* and *blue* are also depleted for genes derived from 31.01 and 36.02 respectively.

**Figure 6.**
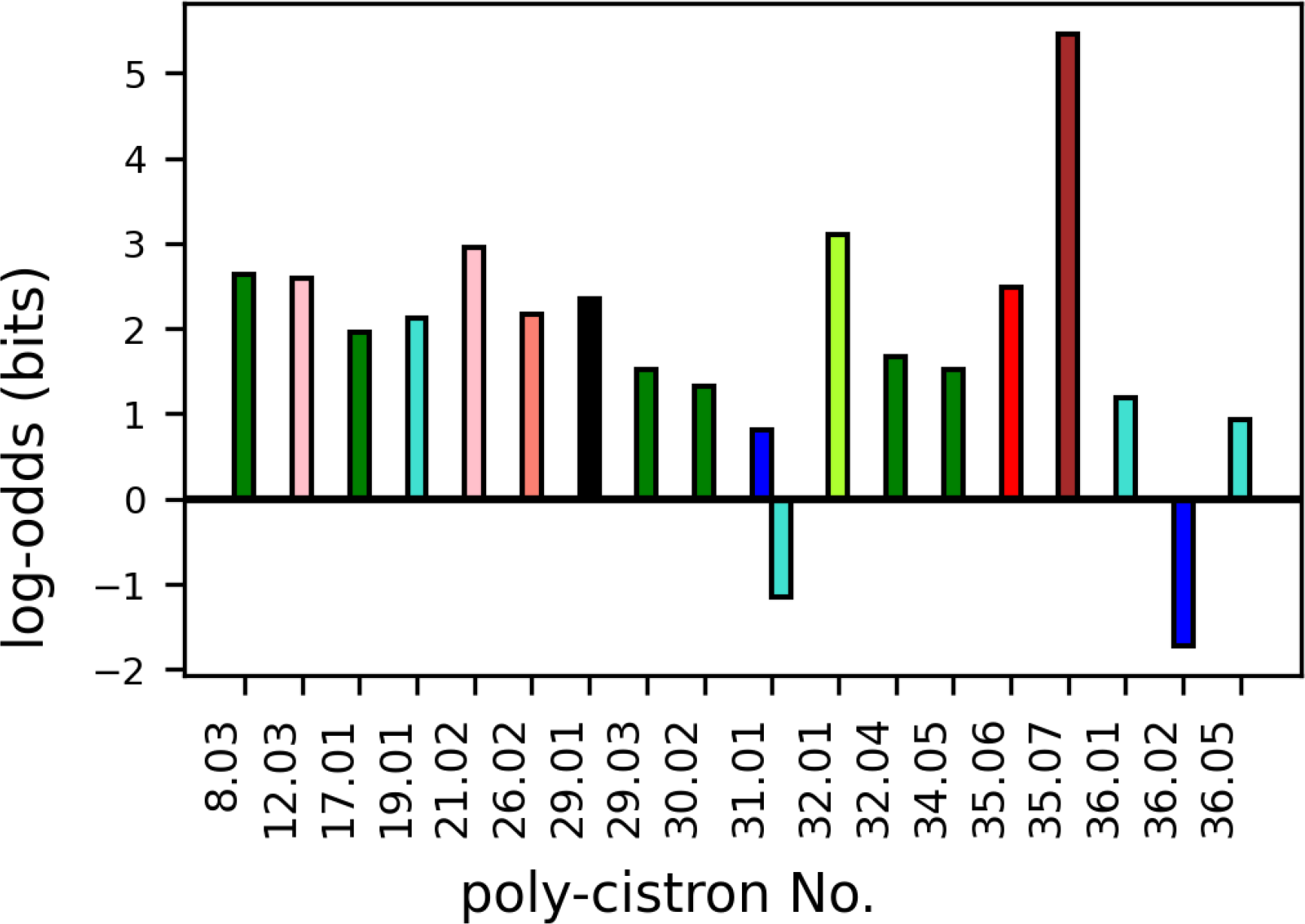
Enrichment of poly-cistronic origin. Color of bar indicates module. [*p*−*value <* 10^−3^]

*Pink* module, enriched with genes encoded on poly-cistron 12.03 is also enriched with genes originating from chromosome 12. However, whereas genes encoded on poly-cistron 21.02 is enriched in *pink*, genes from chromosome 21 aren’t significantly enriched. Conversely, while the *pink* module is enriched with genes from chromosome 2, no corresponding poly-cistronic origin from chromosome 2 appear significantly enriched in it. Similarly, genes originating from poly-cistrons from enriched chromosomes 1, 2, 10, 13 and chromosomes 19, 21, 26, 29, 30, 32 corresponding to enriched poly-cistrons are not enriched.

Thus, while poly-cistron of origin does contribute to co-regulation of some genes, this association of poly-cistronic origin can not be merely attributed to chromosomal involvement described in previous section.

#### 3.5.3 I-ord

The linear arrangement of *Leishmania* genes in co-transcribed poly-cistronic regions results in inherently unequal transcripts. Transcripts may be positioned in their poly-cistron near the beginning, at the end or generally within. Genes at the beginning (5’ end) of the poly-cistron would be less susceptible to potential events of transcriptional abortion than those located further downstream. We tested if this inequality of poly-cistronic location is reflected in association of genes with particular modules. The ordinal number of genes in their respective poly-cistron from the beginning of poly-cistron was marked as *i-ord* and the ordinal number counting from the end, the transcription-terminating *J* nucleotide^16^ was marked as *j-ord*. WGCNA-module classification is associated with *i-ord* (χ^2^ *p*−*value* = 2.344 *×* 10^−18^), however, is independent of *j-ord* (χ^2^ *p*−*value* = 1).

Hence, we tested if location of genes within the polycistron carries regulatory information by analysing association only for each *i-ord* with modules using Fisher’s exact test, *p* −*value <* 10^−3^. Figure (7 A) represents on logarithmic scale, the odds-ratio of genes with specific *i-ord* for modules with significant association. Accordingly, genes located up to 7^th^ *i-ord*, are depleted from the *blue* module while being enriched in *turquoise* or *green* module and the *grey* (non-module).

To check if enrichment of genes with downstream *i-ords* 60, 88, 114, 193 in respectively *dark-green, royal-blue, blue*, and *yellow* is informative, we tested the same positional inequality by similarly analysing for the start codon of each gene. Position of start-codon in the respective poly-cistron was discretized into 2kb bins (discretization into bins is technical phraseology in statistics), and the association of this discretized start-codon position with WGCNA modules was tested (fisher’s exact test). Supporting our earlier observations with *i-ord*, genes with start-codons appearing earlier in their poly-cistron (up to 16kb in the polycistron) are associated with their classification into *blue, turquoise, green* and *grey*. Additionally, genes with start codons located further within the poly-cistronic region are also associated with *dark-turquoise, yellow* and *dark-green*. (Figure 7 B)

**Figure 7.**
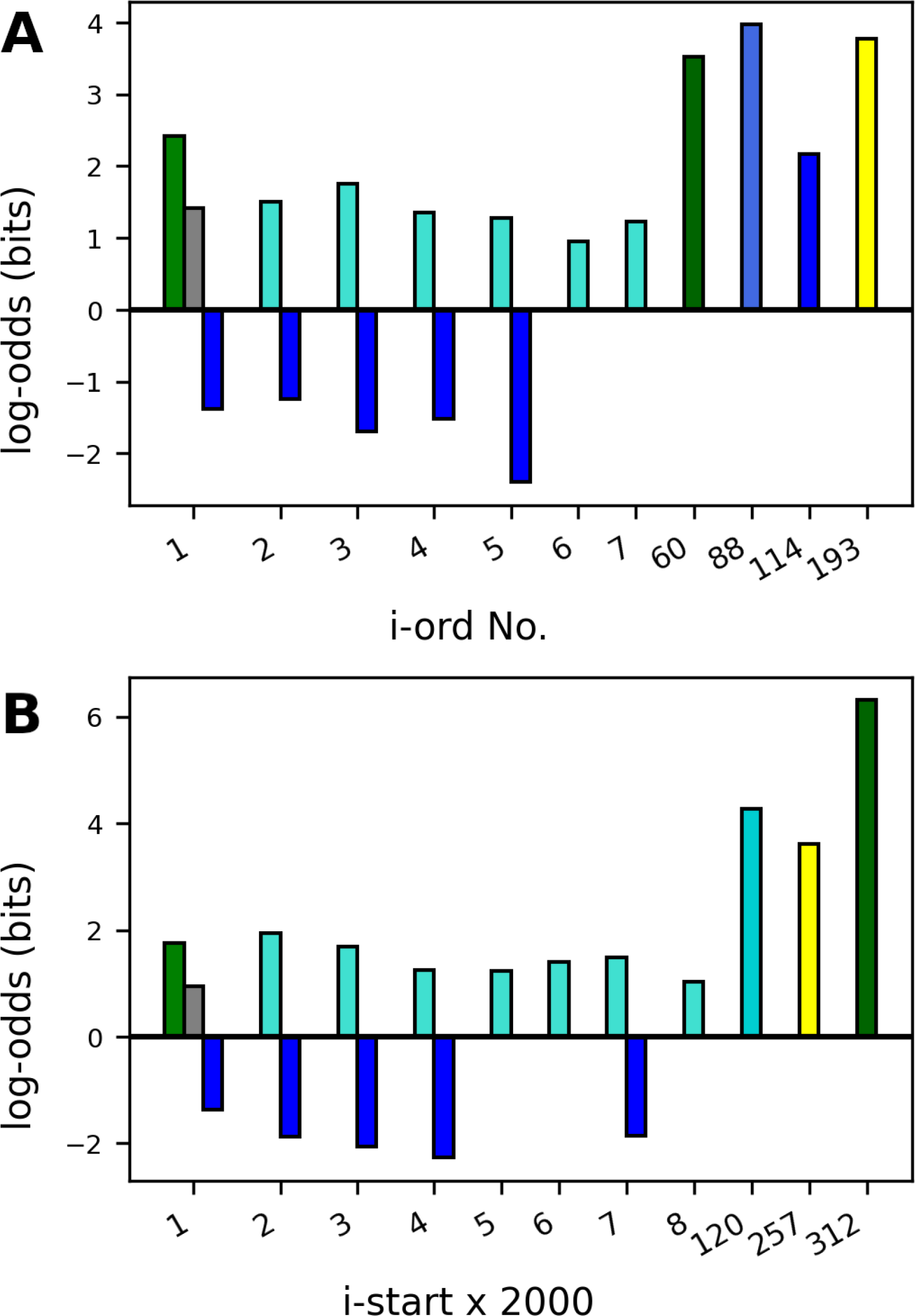
Enrichment based on location in poly-cistron. Color of bar indicates module. [*p*−*value <* 10^−3^] **A**. *i-ord*, rank of gene in its polycistron. B. i-start, 2000 × location (nucleotides) of start codon from beginning of polycistron.

### 3.6 Trait associations

Cell respond to environmental stimuli by modulating functions as necessary, primarily by regulating genes from various regulatory modules. We expect genes belonging to a module to act in unison in order to achieve common broader functions. This should reflect as an association between a common regulatory pattern of genes belonging to a module and biological conditions under consideration. Gene expressions of all genes of a given module is identified during WGCNA can be summarized as a representative ‘eigengene’ expression.

Any module’s eigengene summarizes expression of an *average* gene belonging to that module and may be used as a proxy for behavioural trends of the module. We examined the possibility of association between eigengene expression of each of our identified modules with broad *Leishmania* life-phases by hypergeometric test. Figure (8) is a heat map representing association (red: negative, blue: positive) of each module (rows) against *Leishmania* life-phases (log, stationary, amastigotes, axenic amastigotes). Modules are arranged in ascending order of their average p-value across life phases (most confident at the top). The largest modules, *turquoise* and *blue* associate strongest independently with the log and stationary phases of *Leishmania* respectively. Additionally, *skyblue, salmon, black* modules positively associate with stationary phase.

**Figure 8.**
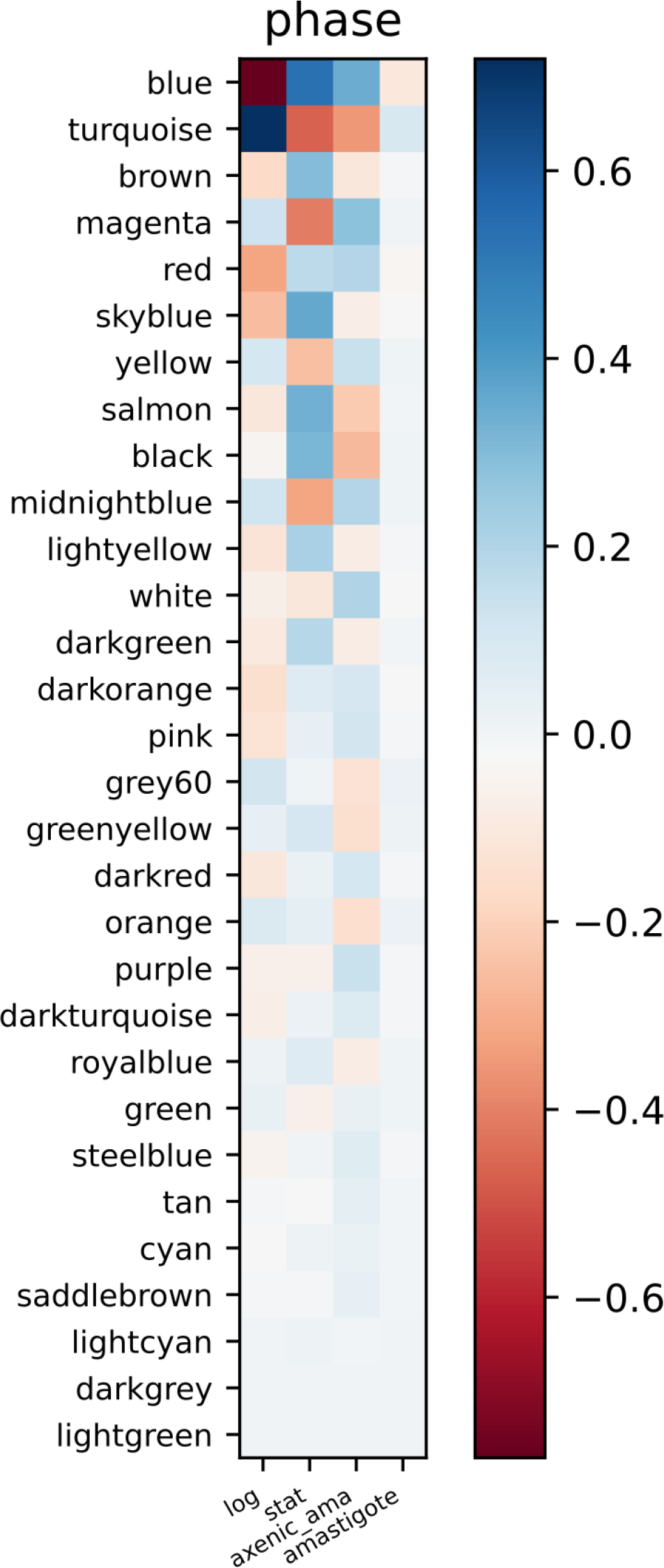
Eigengene association with conditions. ama: amastigotes. Row-order: ascending p-value

*Magenta, skyblue, yellow, salmon, black* differentiate stationary phase from amastigotes by expression in opposite direction while *blue, turquoise, red* differentiate log phase from amastigotes. The association pattern of modules with animal-derived *amastigotes*, despite *noisy*, still agrees with that with *axenic amastigotes*, the corresponding controlled laboratory equivalents.

The clearly weak association of *light-green, darkgrey, lightcyan, saddlebrown, cyan, saddlebrown* modules with any of the life-phases is indicative of lack of discernible modulation of expression of genes belonging to these modules across diverse conditions examined.

### 3.7 Reciprocity

The striking results from trait analysis wherein genes from the *blue* and *turquoise* modules show trends to be mutually exclusive prompted us to analyse properties of these two modules across each of the features we had queried thus far. We find that, *Blue* and *turquoise* reciprocate each other in multiple aspects. Table (3) summarizes enrichment results from previous sections where blue and turquoise modules show opposite trends qua annotation term skew, GO-qualifier skew, chromosomal origin, poly-cistronic origin, i-ord and trait associations. In our analyses, we do not find instances where *blue* and *turquoise* are both enriched or depleted together. Obviously, each enrichment of a given feature in module Blue is not necessarily accompanied by its depletion in *Turquoise*.

Figure 9 is the expression plot for module eigengenes for *blue* and *turquoise* modules against conditions used for analysis, broadly classified under life phases: log, stationary, amastigotes (axenic) and amastigotes. The Eigengene-expression profile indicates that genes in one module generally express in phases in which genes from the other are repressed. While *blue* Eigengene expression is strongly associated with *stationary* life phase, that for *turquoise* is strongly associated with the *log* phase.

**Figure 9.**
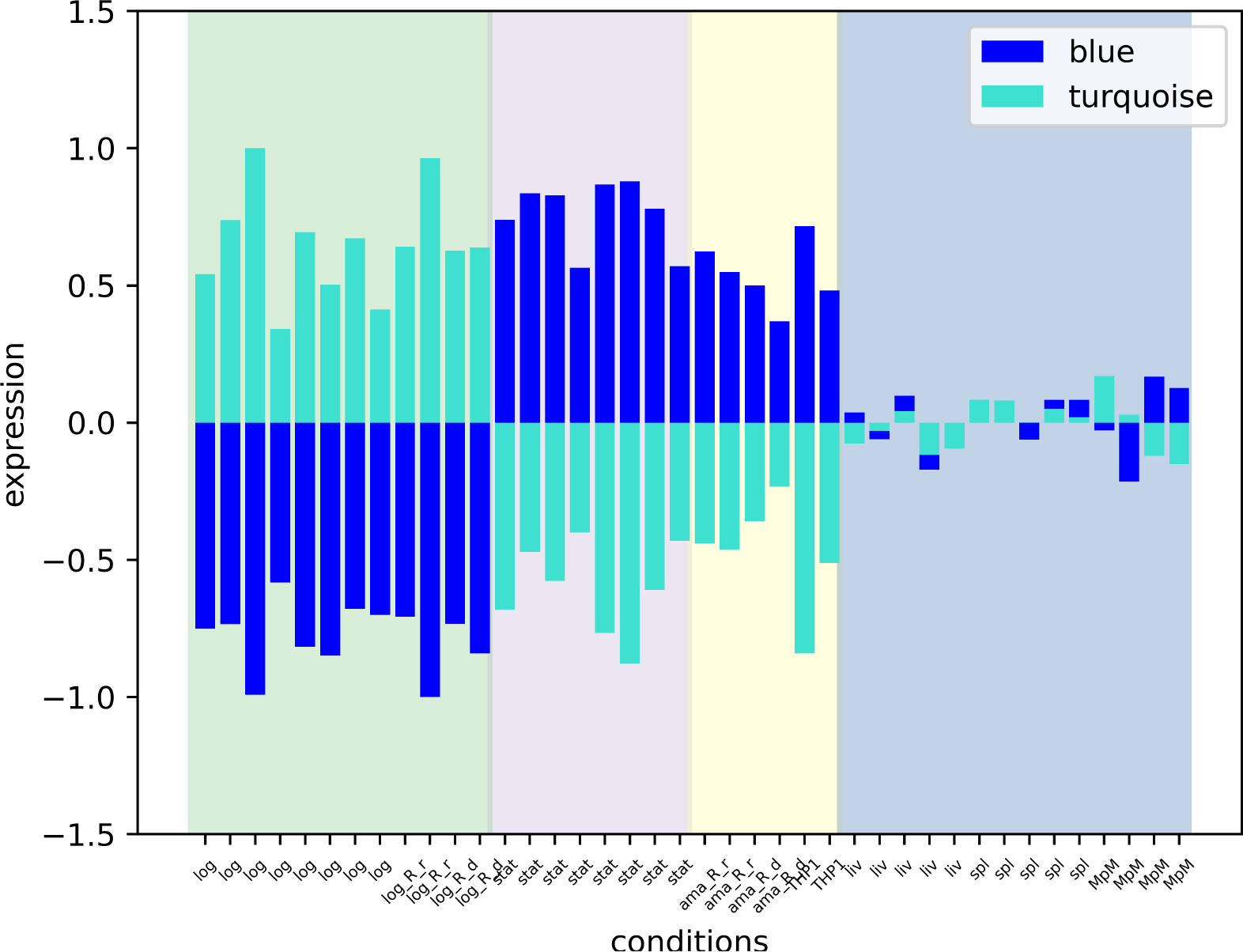
Blue/Turquoise Module Eigengenes. Background groups conditions into broad life-phases. Green: Log, Pink: Stationary, Yellow: Axenic Amastigote, Blue: Wild type Amastigotes

## 4. Discussion

We have sought to understand the global gene-regulatory architecture of *Leishmania donovani* by performing a meta-analysis of currently available transcriptomic data. This analysis of transcriptomes integrated information across diverse conditions to derive inferences going beyond the scope of individual paired studies, thereby giving a global overview of gene-regulatory networks in *Leishmania donovani* We performed WGCNA to classify the mRNA transcripts as annotated in *Leishmania donovani* HU3. The 30 resultant co-regulatory modules, conveniently referenced by various colors, span sizes from 32 (*steelblue*) to 3012 (*turquoise*) genes. The classification was performed based on co-regulation, rather than co-expression of transcripts, to consider both, enhancing as well as repressive genetic effects to form the network. We confirmed validity of the analysis by statistically demonstrating from the available annotations that transcript classification by WGCNA is consistent with currently available gene-classifiers with respect to Gene-Ontology qualifiers, KEGG pathways and choice of terms used by researchers for gene nomenclature. We observe regulatory contribution of the expected RBP-motifs as well as additionally from chromosomal and poly-cistronic origin of the transcripts and their location within the poly-cistron.

The analysis of the distribution of annotation qualifiers across the modules additionally provides following novel information about some modules. The *greenyellow* module is enriched with genes qualified by ‘GO’ as ‘located_in: axoneme’, belong to the KEGG pathway ‘Cilium and associated proteins’, and frequently have names containing the terms, ‘axonemal’, ‘paraflagellar’, ‘rod’. Similarly, *green* module is associated with membrane-terms “GP63”, “heat”, “shock”, “surface”, “ATG”, enriched for KEGG Pathway “Exosome”. The *Lightgreen* module, while strongly associating with “ribosomal” components is also enriched for KEGG Pathway “Ribosome” and qualified by GO as being ‘located_in’ “nucleolus” (Supplementary 4). These agreements across qualifiers in functional description of genes belonging to specific modules further affirm confidence in the clustering. Such observations provide insights towards functional characterization of genes belonging to these modules. The classification of informatically identified putative, conserved and uncharacterized genes into co-regulatory modules provides hints about their characteristics based on those of other annotated genes in their respective modules.

Current dogma of gene regulation in *Leishmania* involves differential gene stabilization by RBPs through identification of cognate motifs present on transcripts. *Leishmania donovani* appear to have inherited the motifs from common ancestors that it shares with the remaining metazoa, which forms the bulk of contributory organisms in the CisBP-RNA database. Alternatively, being obligate parasites within the vector insect or the host mammal, *Leishmania donovani* may have derived some of its RBP-Motif driven gene-regulation in convergence with those operative in their metazoic host. Most RBP-Motifs recognised elsewhere also appear to be meaningfully recognised by *Leishmania donovani*’s gene-regulatory machinery, evident from their overall non-random occurrence in the transcripts. Specifically, we observe contribution of specific RBP-Motifs in coregulatory modules such as the enrichment of M325 in *green*, M331 in *darkorange*, depletion of M195 from *lightgreen*. We have enlisted motifs which are enriched and depleted in each of the regulatory modules, enabling further experimental characterization of the RBP-motifs for demonstration of RBP-actuated regulation.

Our extended analyses of the WGCNA modules beyond the annotation and the current dogma of mRNA-regulation in *Leishmania donovani* through recognition of RBP-Motifs indicates additional regulatory contribution from the chromosomal and poly-cistronic origin of genes, and of their position within their respective poly-cistron. Association of chromosomal origin with regulation agrees with the various observations^17–19^ of stage-specific aneuploidy known in *Leishmania*. In some cases, this association can be further narrowed to point at a single poly-cistron within the chromosome, that is enriched in the same module. This can be observed, for instance, with *green* module enriched with transcripts form chromosomes 8, 17, 34 and respective corresponding poly-cistrons 8.03, 17.01, 34.05, *pink* module with chromodome 12 and poly-cistron 12.03, *red* with chromosome 35 and 35.06. However, it is important to note that poly-cistronic enrichment is also observed for modules such as *salmon* with transcripts from poly-cistron 26.02, *black* with 29.01, *greenyellow* with 32.01, even without a corresponding chromosomal enrichment. Such enrichment of genes from poly-cistrons is indicative of mechanism other than mere enhancement of gene dosage achieved by duplication of the whole chromosome. Poly-cistronic regulation requires existence of regulatory mechanisms that may act before dissociation of individual transcripts through cleavage, either while the immature transcript is still poly-cistronic or even earlier, during events of initiation of transcription. If experimentally demonstrated, this observation may highlight the benefit of the organisation of genes in form of poly-cistronic islands over *Leishmania* chromosomes. However, unlike with RBP motifs and in agreement with the absence of known transcription factors in *Leishmania*, we did not find non-random occurrence of currently listed transcription-factor motifs from other organisms upstream of the poly-cistrons in *Leishmania* (data not shown). Genes located at the beginning of their respective poly-cistron, upto 7^th^ *i-ord*, tend to belong to *turquoise* module and are counter-indicative of *blue* module, a trend indicative of logarithmic promastigotes. Information necessary for such regulatory trends may be encoded at the beginning of the poly-cistron or upstream to it. Such information, however is lacking at the end of poly-cistrons, i.e. at the *J*-nucleotide (*j-ord*). Enrichment of genes located much downstream within the poly-cistron, such 60^th^ i-ord enriched in *green*, 193^rd^ i-ord enriched in *yellow*, etc. was confirmed by enrichment of genes located at a corresponding number of nucleotides from the start of the poly-cistrons. Such downstream enrichment is observed despite lack of enrichment of intermediate transcript-ranks, and away from obvious transcriptional information markers such as terminal boundaries of poly-cistrons. Whether mechanisms such as presence of multiple preferencial ribosomal entry sites in long poly-cistrons or epigenetic effects from histone arrangement propensities cause these enrichment may be of experimental interest.

The two largest modules identified in WGCNA account for about 60% of all classified genes. They individually associate best with the logarithmic (*turquoise*) and stationary (*blue*) growth stages of promastigote forms of *Leishmania donovani*. These modules don’t show common enrichments, and often reciprocate with respect to many attributes considered in this analysis. Eigengenes (9) representing the two modules also show reciprocal directions of gene-regulation across promastigote and axenic amastigote conditions, a trend which reflects in the corresponding phase association (8). This observation hints towards two reciprocal regimes of transcript regulation active in two distinct phases of promastigote maturation, since logarithmic and stationary growth stages roughly simulate the early pro-cyclic promastigotes, that have recently entered the vector’s gut and late meta-cyclic promastigotes that are ready to infect the next host respectively.

After batch-correction and normalization of transcripts, a preliminary clustering of the experimental conditions had already indicated that phases which may be broadly called ‘stationary promastigotes’ cluster transcriptionally closer with ‘amastigotes’ rather than the morphologically similar ‘logarithmic promastigotes’. This is further independently supported by the similarity of associative trends of many co-regulatory modules with stationary promastigotes and axenic amastigotes (8). This is suggestive of a preparatory ‘amastigote-like’ environment developed by *Leishmania donovani* while already in the stationary promastigote phase poised for infection into a new host.

The minimal association of the modules *lightgreen, dark-grey, lightcyan, saddlebrown* with any of the analysed experimental conditions identifies sets of genes which are minimally modulated across the life-phases of *Leishmania*. Genes in these modules may be essential for *Leishmania donovani* at a certain level across various conditions. For example, the *lightgreen* module identified in this category also associates with *Ribosomal* structures, supporting the idea that cells tend to maintain constant amounts of ribosomal function. Genes in such invariant modules have a potential for being novel drug-targets as well as promising candidates for relative quantification of transcription by techniques such as qPCR. Our analysis provide associative observations that identify plausible additional repositories of gene-regulatory information. The results are arrived at through statistical association demonstrated across diverse transcriptional datasets derived by various research groups, batch-corrected to remove experimental biases, if any. These may form starting points for exploration of biochemical mechanisms that bring about the regulation.

For rapid simplified analysis of RBP (or TF) motifs, we have considered presence of the motif, at least once anywhere within the transcript as sufficient for its recognition by the corresponding regulator within *Leishmania donovani*. However, it is evident that multiplicity of occurrence, exact sequence and location(s) of the motif within the transcript may have an effect on the signalling and binding strength of the motifs. This complexity may be further compounded by harmonious or conflicting interplay between similar or different motifs, strengths of such synergies and factors influencing them. As bio-chemical mechanism of RBP-Motif interactions becomes clearer, motif-search may be further enhanced to quantify and include these factors. Similarly, *Leishmania donovani* have been shown to utilize alternative poly-adenylation sites (PAS), whose locations are available in the referred GFF file. Mere mapping of transcriptome reads withing the scope of a given gene is insufficient to attribute the read to any specific PAS among the putative alternatives. A chemical methodology that recognises the choice of PAS corresponding to each read or better knowledge about mechanism of utilization of alternate PAS in *Leishmania* would be essential for inclusion of regulatory information that they may encode in a similar meta-analysis. Isoforms of each transcript (corresponding to each alternate PAS), rather than the full-length of annotated transcripts would then serve as the unit of regulatory classification. Larger modules such as *blue* and *turquoise*, as well as to classify currently unclassified genes (*grey*) identified in this WGCNA can be resolved further as transcriptomes become available for additional conditions.

Overall, we have provided an overview of gene-regulatory network in *Leishmania donovani* as genetic modules. While confirming contribution of RBP-motifs in gene-regulation, we have provided evidence for existence of additional contributory factors such as chromosomal and poly-cistronic origin of genes and their location within their poly-cistron, as also the identification of a pair of reciprocating gene regulatory modules associating with logarithmic and stationary promastigote phases of *Leishmania*. Analyses of gene enrichment or depletion from these modules have led to indication of additional modes of gene-regulation in *Leishmania*. Such meta information relating any gene of interest to its co-regulatory cohort may provide initial aid towards its characterization. Contribution of any other putative mechanism towards formation of regulatory modules may be tested using the developed tool-set. The pipeline developed for using diverse transcriptome datasets should aid further granular resolution of modules through quick inclusion of transcriptomes which may become available in future. Many of our observations about *Leishmania donovani* should be conserved across the genus in general. Similar analyses for other *Leishmania* species will help identify conserved modular structure in *Leishmania*, as also to identify subtle differences in modular organisation of genes, both of which will aid informed design of intervention strategies, understand tropism, insect specificity and overall biology of the pathogen.

## Supporting information

Enrichments

Polycistron Annotation

## Notes

### Competing Interest Statement

The authors have declared no competing interest.

