## Supplementary material for "Meta-analysis of the pathogen *Leishmania donovani*’s transcriptome reveals multiple modes of regulation including two reciprocally regulated gene modules": Enrichments: acts_upstream_of_or_within_log_odds.pdf

### Contents

| Modules | Pathway | log-odds (bits) |
|---|---|---|
| turquoise | cell cycle DNA replication | 4.82126 |
| | ubiquitin-dependent protein catabolic process | 4.65058 |
| | long-chain fatty acid biosynthetic process | 4.65058 |
| | chromosome segregation | 4.23841 |
| | post-transcriptional regulation of gene expression | 1.39289 |
| blue | post-transcriptional regulation of gene expression | -0.975099 |
| green | post-transcriptional regulation of gene expression | -inf |
