## Supplementary material for "Meta-analysis of the pathogen *Leishmania donovani*’s transcriptome reveals multiple modes of regulation including two reciprocally regulated gene modules": Enrichments: kegg_path_level3_log_odds.org

# -*- mode: org; coding: utf-8; -*-
#+SETUPFILE: ../.automation-scripts/setup.org
#+TITLE: Pathways
#+OPTIONS: toc:nil

#+CAPTION: Enrichment of genes belonging to Pathways. Color of bar indicates module. [$p-value < 10^{-3}$]
#+LABEL: fig_path
#+ATTR_LATEX: :width \linewidth
[[file:../img/]]


#+CAPTION: Names of pathways, genes belonging to which, are enriched in at least 1 module.
|------------+---------------------------------------------+-----------------|
| Modules    | Pathway                                     | log-odds (bits) |
|------------+---------------------------------------------+-----------------|
| turquoise  | Cysteine and methionine metabolism          |         3.38554 |
|            | Phagosome                                   |         3.25954 |
|            | Proteasome                                  |         3.25081 |
|            | Amino sugar and nucleotide sugar metabolism |          3.0863 |
|            | DNA replication proteins                    |         2.65313 |
|            | Chaperones and folding catalysts            |         2.32139 |
|            | Translation factors                         |         2.24943 |
|            | Oxidative phosphorylation                   |         2.06437 |
|            | Chromosome and associated proteins          |         1.91773 |
|            | Cytoskeleton proteins                       |         1.86252 |
|            | DNA repair and recombination proteins       |         1.78496 |
|            | Exosome                                     |         1.68616 |
|            | Cilium and associated proteins              |         1.39305 |
|------------+---------------------------------------------+-----------------|
| blue       | Exosome                                     |        -2.37174 |
|            | Ribosome                                    |        -2.77097 |
|------------+---------------------------------------------+-----------------|
| green      | Exosome                                     |         1.67107 |
|------------+---------------------------------------------+-----------------|
| lightgreen | Ribosome                                    |         7.33701 |
|------------+---------------------------------------------+-----------------|
| magenta    | Arginine biosynthesis                       |         7.45084 |
|------------+---------------------------------------------+-----------------|
| yellow     | Peptidases and inhibitors                   |         2.06739 |
|------------+---------------------------------------------+-----------------|
