## Supplementary material for "Meta-analysis of the pathogen *Leishmania donovani*’s transcriptome reveals multiple modes of regulation including two reciprocally regulated gene modules": Enrichments: kegg_path_level3_log_odds.pdf

### Pathways

Pradyumna Paranjape

December 20, 2023

`../img/`

Table 1: Names of pathways, genes belonging to which, are enriched in at least 1 module.

| Modules | Pathway | log-odds (bits) |
|---|---|---:|
| turquoise | Cysteine and methionine metabolism | 3.38554 |
| | Phagosome | 3.25954 |
| | Proteasome | 3.25081 |
| | Amino sugar and nucleotide sugar metabolism | 3.0863 |
| | DNA replication proteins | 2.65313 |
| | Chaperones and folding catalysts | 2.32139 |
| | Translation factors | 2.24943 |
| | Oxidative phosphorylation | 2.06437 |
| | Chromosome and associated proteins | 1.91773 |
| | Cytoskeleton proteins | 1.86252 |
| | DNA repair and recombination proteins | 1.78496 |
| | Exosome | 1.68616 |
| | Cilium and associated proteins | 1.39305 |
| blue | Exosome | -2.37174 |
| | Ribosome | -2.77097 |
| green | Exosome | 1.67107 |
| lightgreen | Ribosome | 7.33701 |
| magenta | Arginine biosynthesis | 7.45084 |
| yellow | Peptidases and inhibitors | 2.06739 |
