## Supplementary material for "Meta-analysis of the pathogen *Leishmania donovani*’s transcriptome reveals multiple modes of regulation including two reciprocally regulated gene modules": Enrichments: kegg_path_level3_log_odds.tex

% Created 2023-12-20 Wed 16:01
% Intended LaTeX compiler: pdflatex
\documentclass[11pt]{article}
\usepackage[utf8]{inputenc}
\usepackage[T1]{fontenc}
\usepackage{graphicx}
\usepackage{longtable}
\usepackage{wrapfig}
\usepackage{rotating}
\usepackage[normalem]{ulem}
\usepackage{amsmath}
\usepackage{amssymb}
\usepackage{capt-of}
\usepackage{hyperref}
\author{Pradyumna Paranjape}
\date{\today}
\title{Pathways}
\hypersetup{
 pdfauthor={Pradyumna Paranjape},
 pdftitle={Pathways},
 pdfkeywords={},
 pdfsubject={},
 pdfcreator={Emacs 29.1.90 (Org mode 9.6.13)}, 
 pdflang={English}}
\begin{document}

\maketitle
\url{../img/}


\begin{table}[htbp]
\caption{Names of pathways, genes belonging to which, are enriched in at least 1 module.}
\centering
\begin{tabular}{llr}
\hline
Modules & Pathway & log-odds (bits)\\[0pt]
\hline
turquoise & Cysteine and methionine metabolism & 3.38554\\[0pt]
 & Phagosome & 3.25954\\[0pt]
 & Proteasome & 3.25081\\[0pt]
 & Amino sugar and nucleotide sugar metabolism & 3.0863\\[0pt]
 & DNA replication proteins & 2.65313\\[0pt]
 & Chaperones and folding catalysts & 2.32139\\[0pt]
 & Translation factors & 2.24943\\[0pt]
 & Oxidative phosphorylation & 2.06437\\[0pt]
 & Chromosome and associated proteins & 1.91773\\[0pt]
 & Cytoskeleton proteins & 1.86252\\[0pt]
 & DNA repair and recombination proteins & 1.78496\\[0pt]
 & Exosome & 1.68616\\[0pt]
 & Cilium and associated proteins & 1.39305\\[0pt]
\hline
blue & Exosome & -2.37174\\[0pt]
 & Ribosome & -2.77097\\[0pt]
\hline
green & Exosome & 1.67107\\[0pt]
\hline
lightgreen & Ribosome & 7.33701\\[0pt]
\hline
magenta & Arginine biosynthesis & 7.45084\\[0pt]
\hline
yellow & Peptidases and inhibitors & 2.06739\\[0pt]
\hline
\end{tabular}
\end{table}
\end{document}
