## Supplementary material for "Meta-analysis of the pathogen *Leishmania donovani*’s transcriptome reveals multiple modes of regulation including two reciprocally regulated gene modules": Enrichments: located_in_log_odds.pdf

### Contents

| Modules | Pathway | log-odds (bits) |
|---|---|---|
| turquoise | cortical cytoskeleton | inf |
| | mitochondrial outer membrane | 1.82101 |
| | glycosome | 1.50607 |
| | axoneme | 1.41892 |
| | ciliary basal body | 1.13597 |
| | mitochondrion | 0.997882 |
| | nucleoplasm | 0.946244 |
| | kinetoplast | 0.91153 |
| | ciliary plasm | 0.810482 |
| | cytoplasm | 0.509139 |
| blue | cytoplasm | -0.426849 |
| | cilium | -1.84241 |
| | axoneme | -3.38921 |
| green | cytoplasm | -1.49227 |
| | mitochondrion | -3.54584 |
| lightgreen | nucleolus | 2.71899 |
| | cytoplasm | 2.4179 |
| yellow | endoplasmic reticulum | 1.63519 |
| | axoneme | -inf |
| cyan | nucleolus | 2.51482 |
| darkorange | nucleolus | 3.26951 |
| greenyellow | axoneme | 3.81191 |
| grey60 | axoneme | 4.00421 |
