## Supplementary material for "Meta-analysis of the pathogen *Leishmania donovani*’s transcriptome reveals multiple modes of regulation including two reciprocally regulated gene modules": Enrichments: located_in_log_odds.tex

% Created 2023-12-20 Wed 16:01
% Intended LaTeX compiler: pdflatex
\documentclass[11pt]{article}
\usepackage[utf8]{inputenc}
\usepackage[T1]{fontenc}
\usepackage{graphicx}
\usepackage{longtable}
\usepackage{wrapfig}
\usepackage{rotating}
\usepackage[normalem]{ulem}
\usepackage{amsmath}
\usepackage{amssymb}
\usepackage{capt-of}
\usepackage{hyperref}
\author{Pradyumna Paranjape}
\date{\today}
\title{}
\hypersetup{
 pdfauthor={Pradyumna Paranjape},
 pdftitle={},
 pdfkeywords={},
 pdfsubject={},
 pdfcreator={Emacs 29.1.90 (Org mode 9.6.13)}, 
 pdflang={English}}
\begin{document}

\tableofcontents

\begin{center}
\begin{tabular}{llr}
\hline
Modules & Pathway & log-odds (bits)\\[0pt]
\hline
turquoise & cortical cytoskeleton & inf\\[0pt]
 & mitochondrial outer membrane & 1.82101\\[0pt]
 & glycosome & 1.50607\\[0pt]
 & axoneme & 1.41892\\[0pt]
 & ciliary basal body & 1.13597\\[0pt]
 & mitochondrion & 0.997882\\[0pt]
 & nucleoplasm & 0.946244\\[0pt]
 & kinetoplast & 0.91153\\[0pt]
 & ciliary plasm & 0.810482\\[0pt]
 & cytoplasm & 0.509139\\[0pt]
\hline
blue & cytoplasm & -0.426849\\[0pt]
 & cilium & -1.84241\\[0pt]
 & axoneme & -3.38921\\[0pt]
\hline
green & cytoplasm & -1.49227\\[0pt]
 & mitochondrion & -3.54584\\[0pt]
\hline
lightgreen & nucleolus & 2.71899\\[0pt]
 & cytoplasm & 2.4179\\[0pt]
\hline
yellow & endoplasmic reticulum & 1.63519\\[0pt]
 & axoneme & -inf\\[0pt]
\hline
cyan & nucleolus & 2.51482\\[0pt]
\hline
darkorange & nucleolus & 3.26951\\[0pt]
\hline
greenyellow & axoneme & 3.81191\\[0pt]
\hline
grey60 & axoneme & 4.00421\\[0pt]
\hline
\end{tabular}
\end{center}
\end{document}
