## Supplementary material for "Meta-analysis of the pathogen *Leishmania donovani*’s transcriptome reveals multiple modes of regulation including two reciprocally regulated gene modules": Enrichments: part_of_log_odds.org

|-----------+-------------------------------------------------------------------------------+-----------------|
| Module    | Pathway                                                                       | log-odds (bits) |
|-----------+-------------------------------------------------------------------------------+-----------------|
| turquoise | mitochondrial outer membrane translocase complex                              |             inf |
|           | mitochondrial proton-transporting ATP synthase complex                        |             inf |
|           | eukaryotic translation initiation factor 3 complex                            |             inf |
|           | mitochondrial proton-transporting ATP synthase complex, catalytic sector F(1) |         4.32088 |
|           | proteasome regulatory particle                                                |         4.06106 |
|           | CMG complex                                                                   |         3.73745 |
|           | intraciliary transport particle                                               |          2.5728 |
|           | mitochondrial small ribosomal subunit                                         |         1.74715 |
|           | mitochondrial large ribosomal subunit                                         |         1.55286 |
|-----------+-------------------------------------------------------------------------------+-----------------|
