## Supplementary material for "Meta-analysis of the pathogen *Leishmania donovani*’s transcriptome reveals multiple modes of regulation including two reciprocally regulated gene modules": Enrichments: part_of_log_odds.pdf

### Contents

| Module | Pathway | log-odd |
|---|---|---|
| turquoise | mitochondrial outer membrane translocase complex | |
| | mitochondrial proton-transporting ATP synthase complex | |
| | eukaryotic translation initiation factor 3 complex | |
| | mitochondrial proton-transporting ATP synthase complex, catalytic sector F(1) | |
| | proteasome regulatory particle | |
| | CMG complex | |
| | intraciliary transport particle | |
| | mitochondrial small ribosomal subunit | |
| | mitochondrial large ribosomal subunit | |
