## Supplementary material for "Meta-analysis of the pathogen *Leishmania donovani*’s transcriptome reveals multiple modes of regulation including two reciprocally regulated gene modules": Enrichments: part_of_log_odds.tex

% Created 2023-12-20 Wed 16:01
% Intended LaTeX compiler: pdflatex
\documentclass[11pt]{article}
\usepackage[utf8]{inputenc}
\usepackage[T1]{fontenc}
\usepackage{graphicx}
\usepackage{longtable}
\usepackage{wrapfig}
\usepackage{rotating}
\usepackage[normalem]{ulem}
\usepackage{amsmath}
\usepackage{amssymb}
\usepackage{capt-of}
\usepackage{hyperref}
\author{Pradyumna Paranjape}
\date{\today}
\title{}
\hypersetup{
 pdfauthor={Pradyumna Paranjape},
 pdftitle={},
 pdfkeywords={},
 pdfsubject={},
 pdfcreator={Emacs 29.1.90 (Org mode 9.6.13)}, 
 pdflang={English}}
\begin{document}

\tableofcontents

\begin{center}
\begin{tabular}{llr}
\hline
Module & Pathway & log-odds (bits)\\[0pt]
\hline
turquoise & mitochondrial outer membrane translocase complex & inf\\[0pt]
 & mitochondrial proton-transporting ATP synthase complex & inf\\[0pt]
 & eukaryotic translation initiation factor 3 complex & inf\\[0pt]
 & mitochondrial proton-transporting ATP synthase complex, catalytic sector F(1) & 4.32088\\[0pt]
 & proteasome regulatory particle & 4.06106\\[0pt]
 & CMG complex & 3.73745\\[0pt]
 & intraciliary transport particle & 2.5728\\[0pt]
 & mitochondrial small ribosomal subunit & 1.74715\\[0pt]
 & mitochondrial large ribosomal subunit & 1.55286\\[0pt]
\hline
\end{tabular}
\end{center}
\end{document}
