## Supplementary figures and images for "Meta-analysis of the pathogen *Leishmania donovani*’s transcriptome reveals multiple modes of regulation including two reciprocally regulated gene modules"

### acts_upstream_of_or_within_log_odds.png

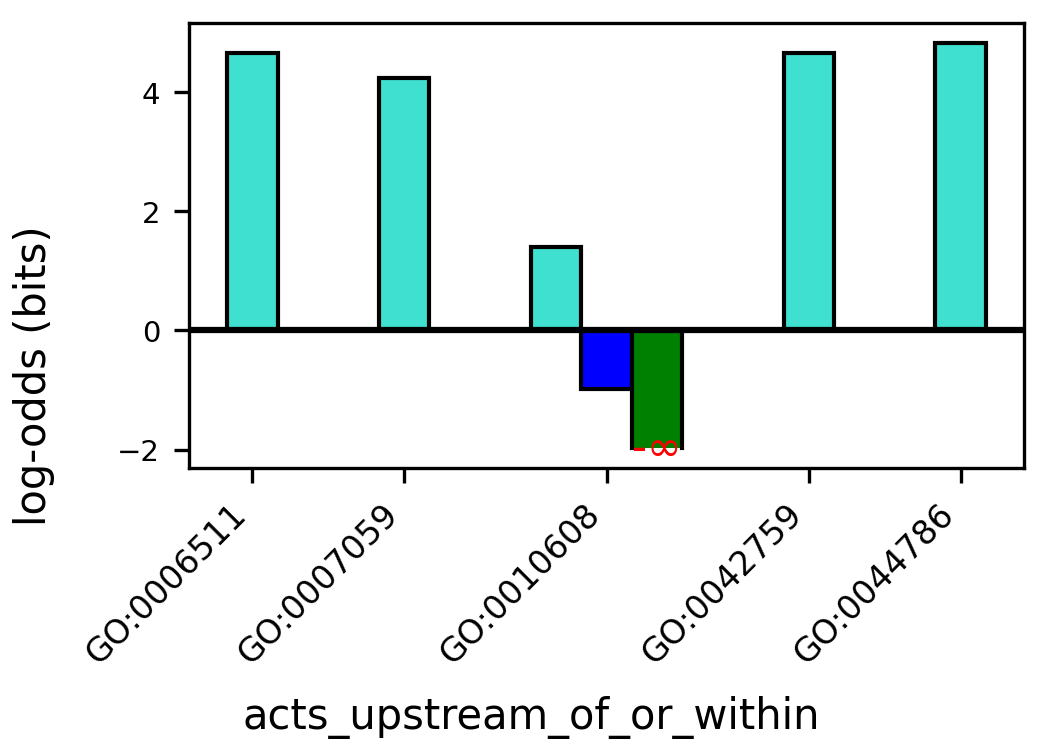

### aspect_log_odds.png

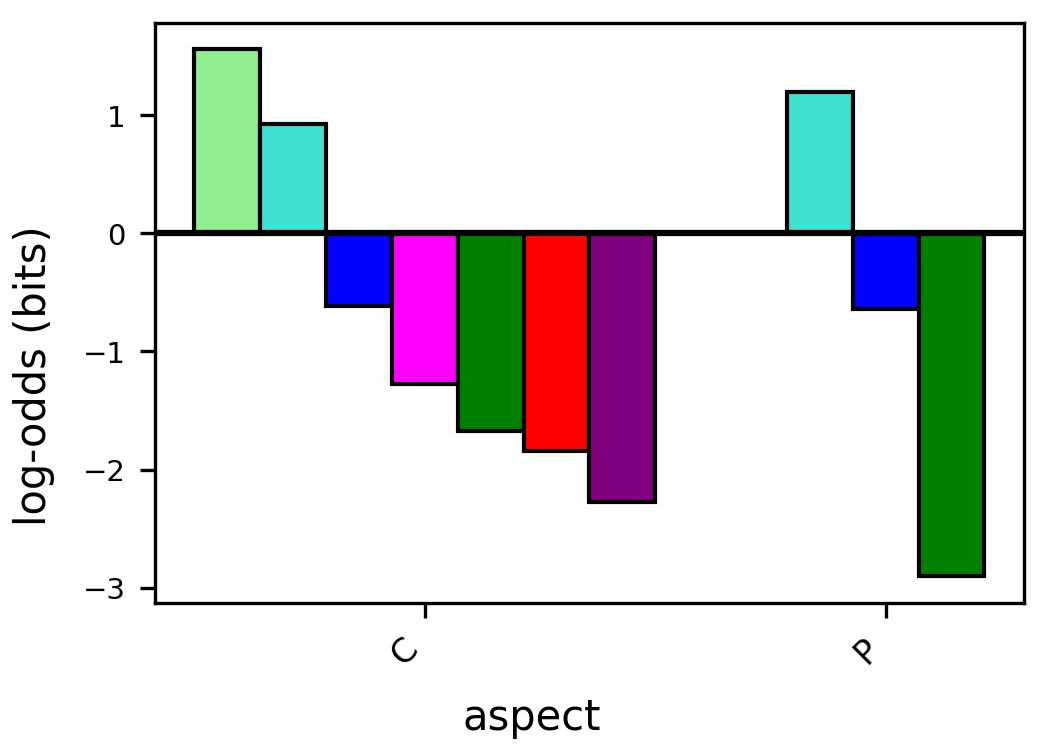

### chromosome_log_odds.png

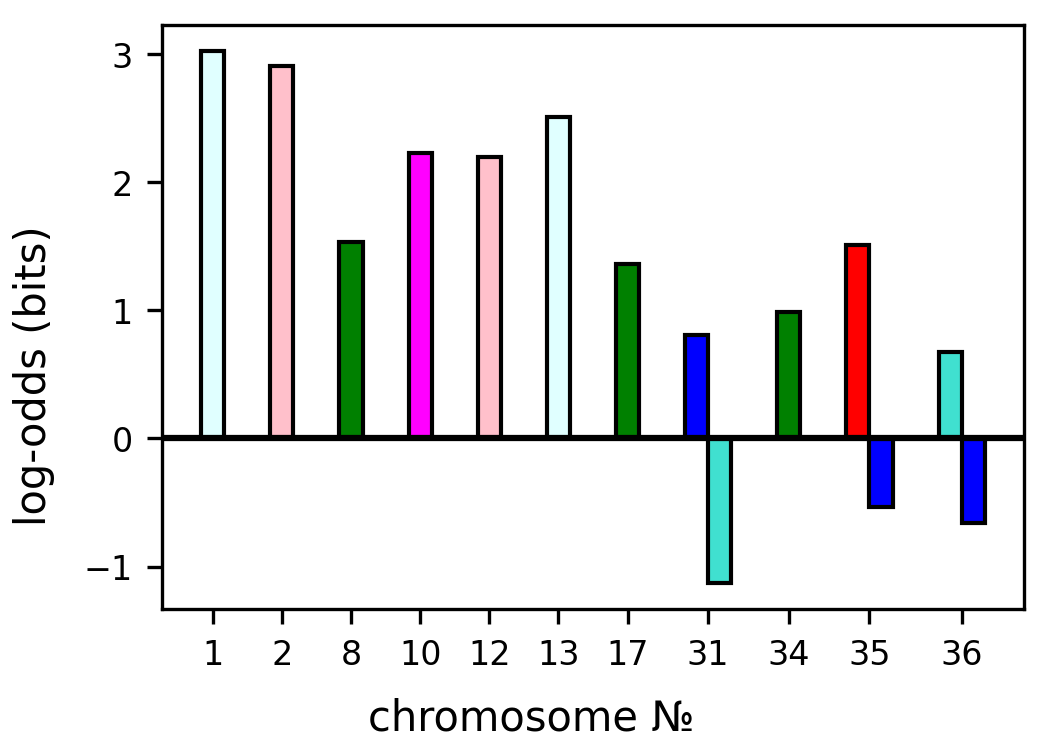

### GO-terms_blue_log_odds.png

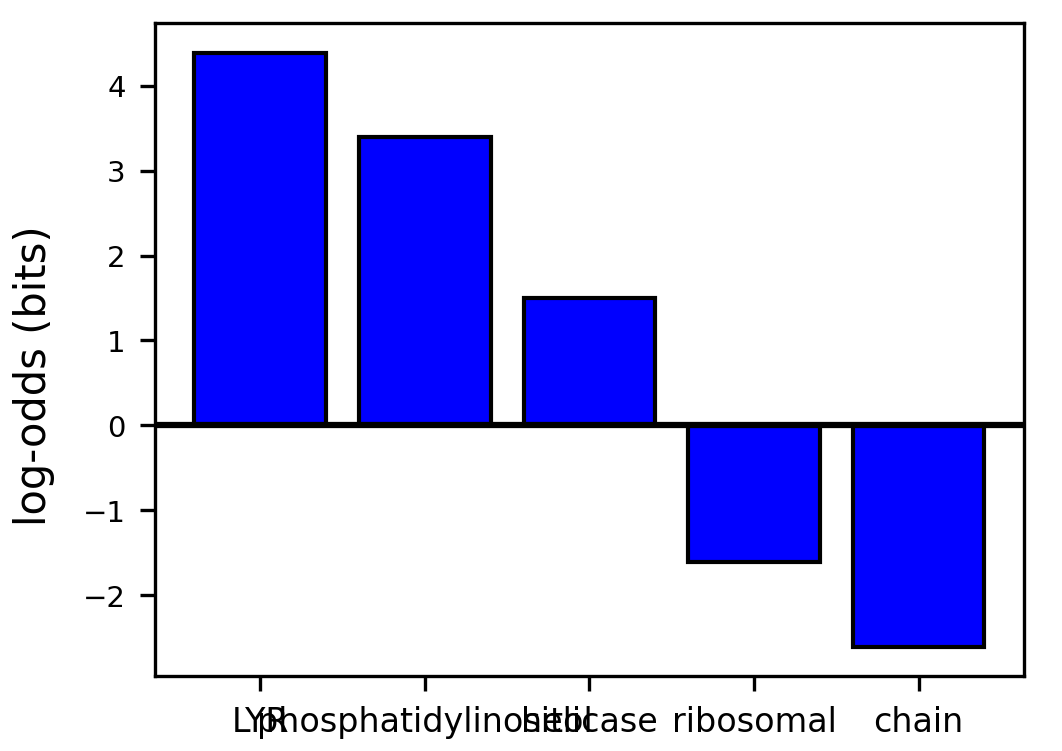

### GO-terms_brown_log_odds.png

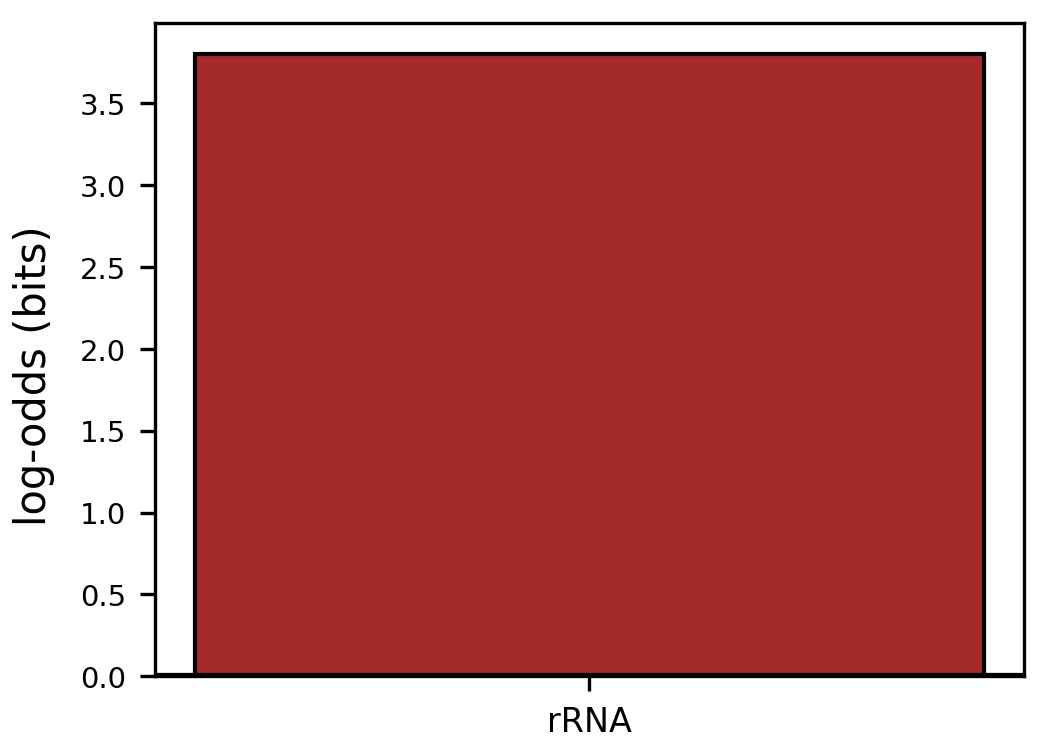

### GO-terms_green_log_odds.png

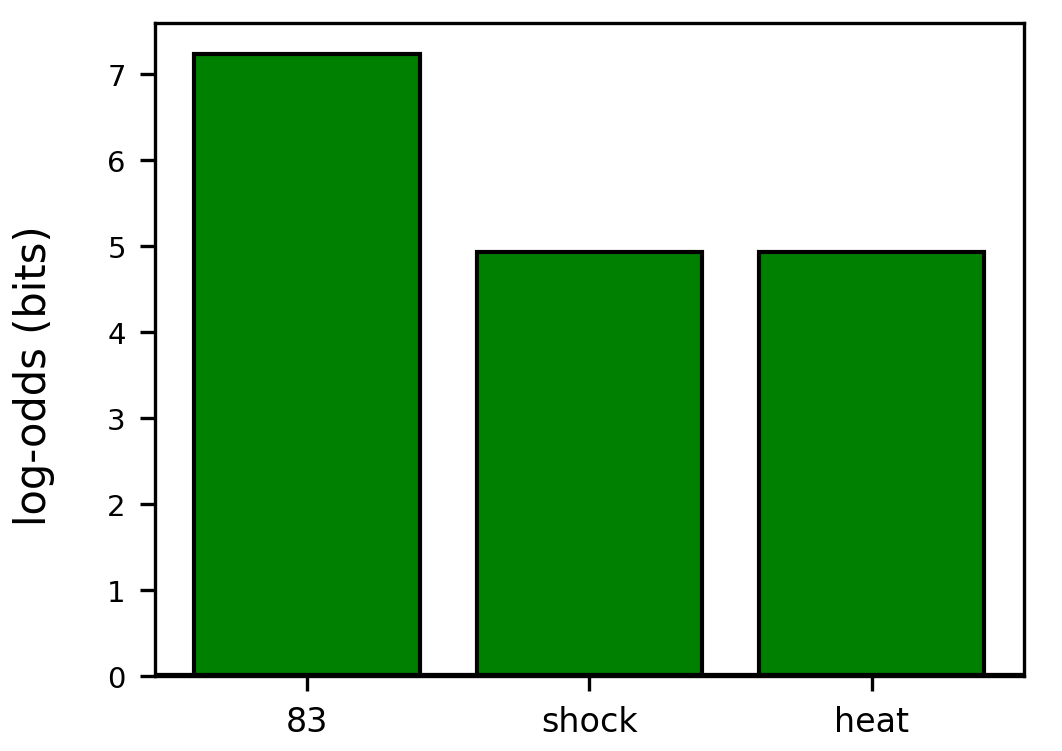

### GO-terms_greenyellow_log_odds.png

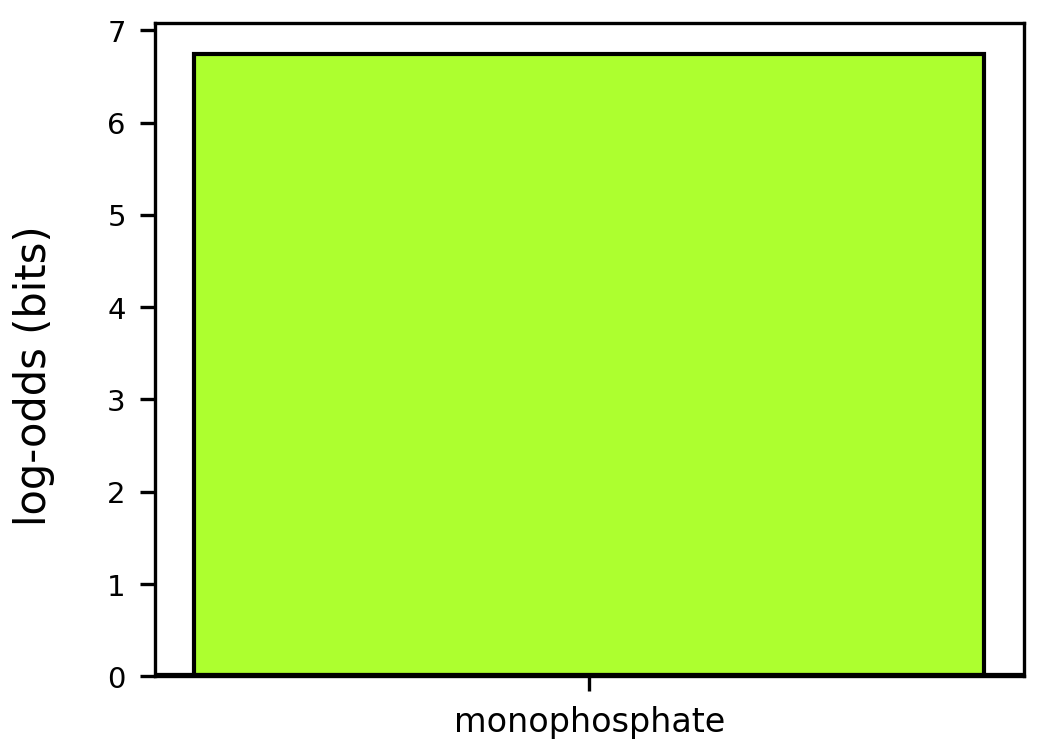

### GO-terms_lightgreen_log_odds.png

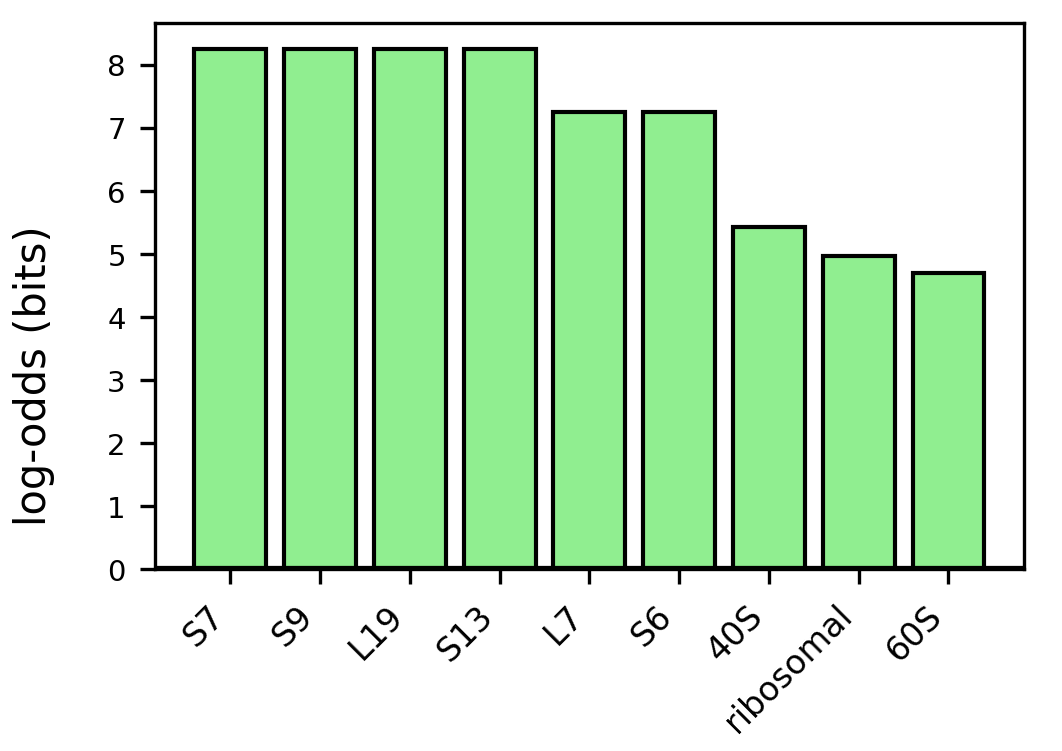

### GO-terms_lightyellow_log_odds.png

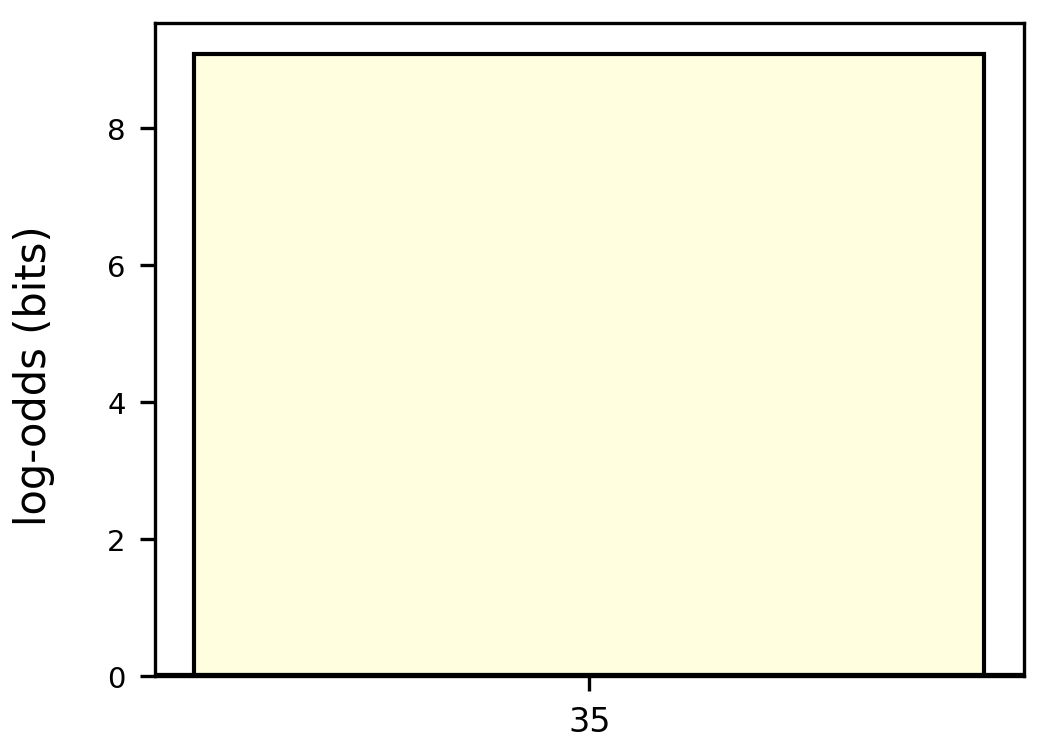

### GO-terms_log_odds.png

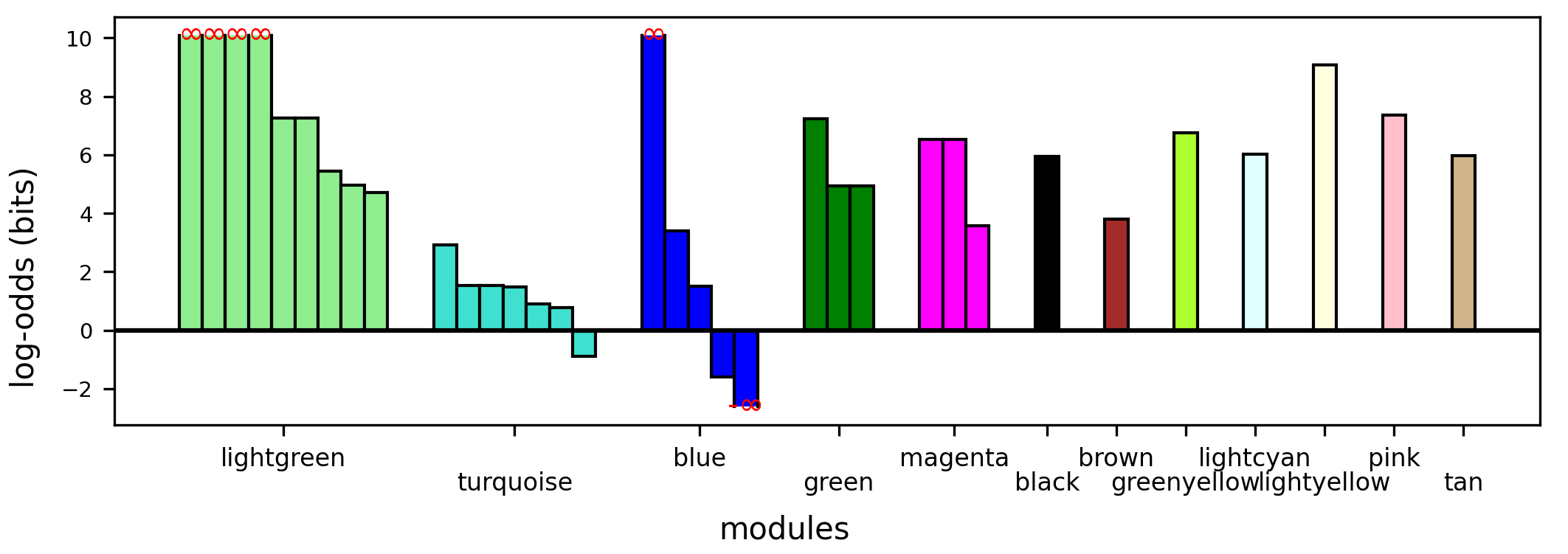

### GO-terms_magenta_log_odds.png

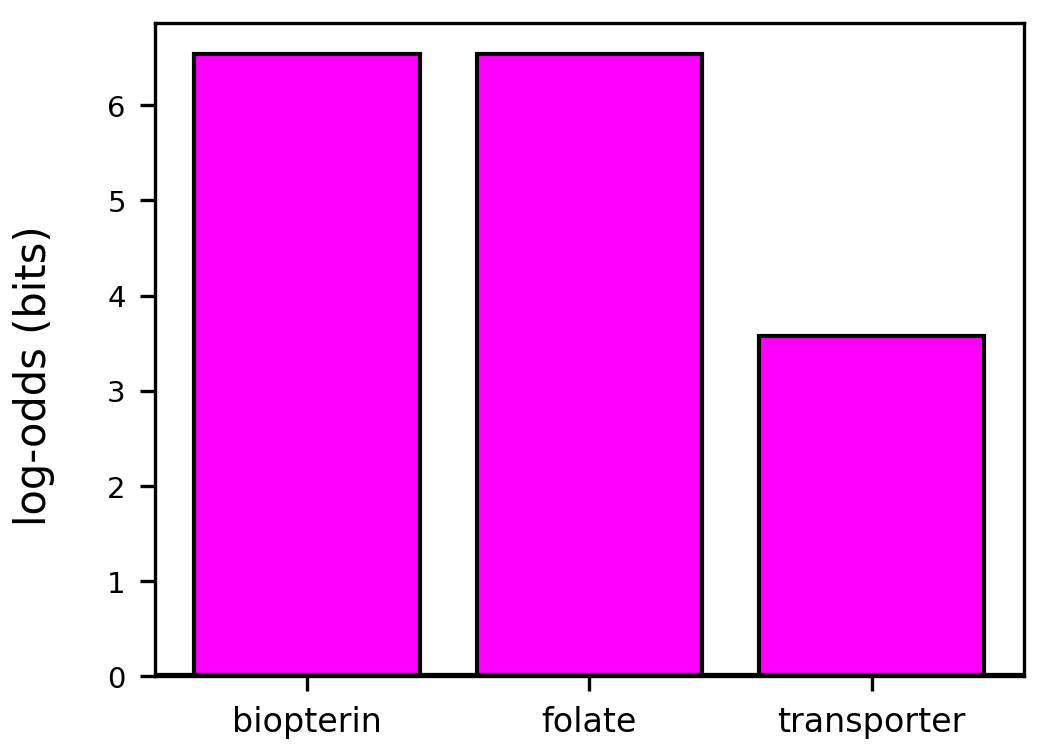

### GO-terms_num.png

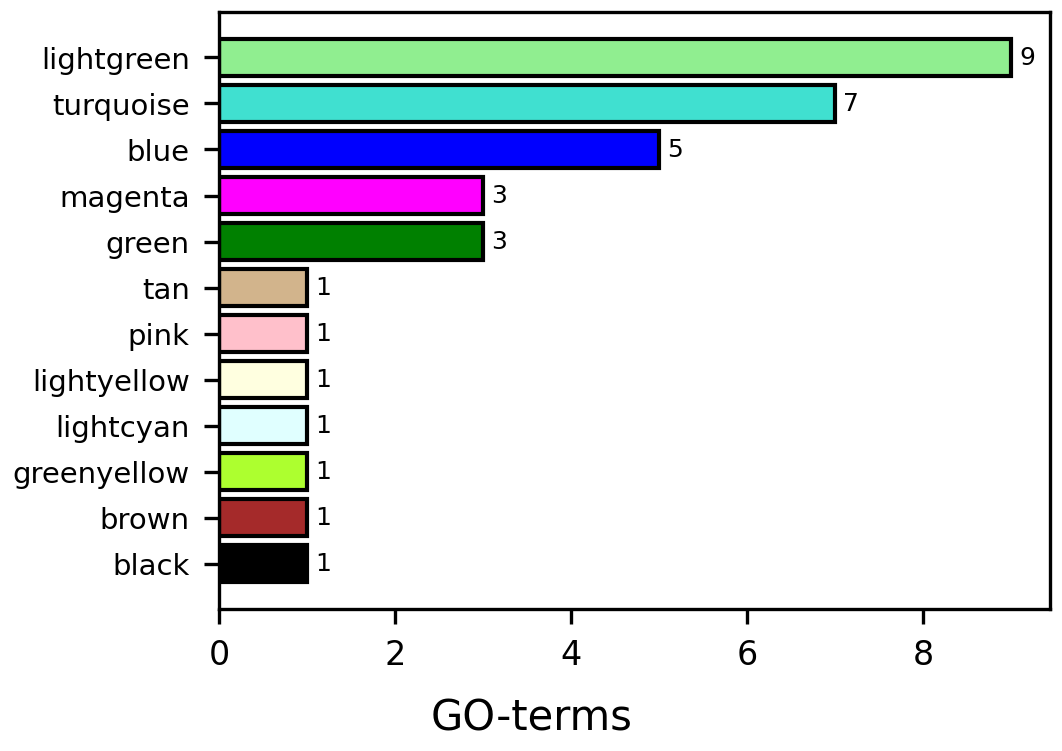

### GO-terms_pink_log_odds.png

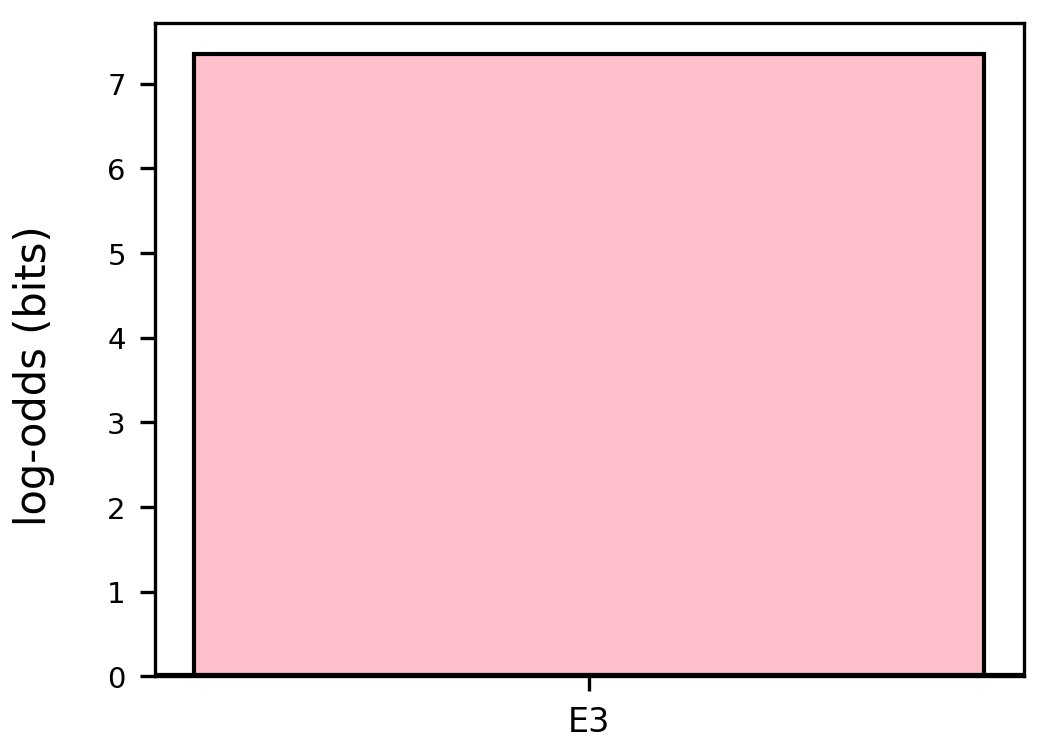

### GO-terms_tan_log_odds.png

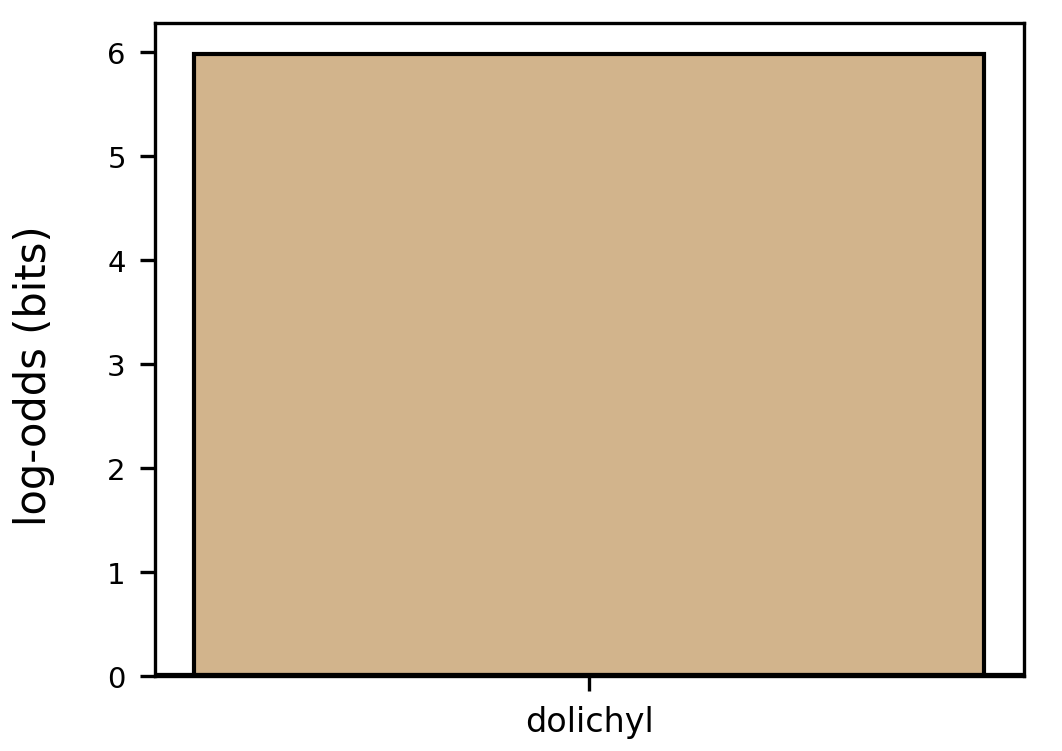

### GO-terms_turquoise_log_odds.png

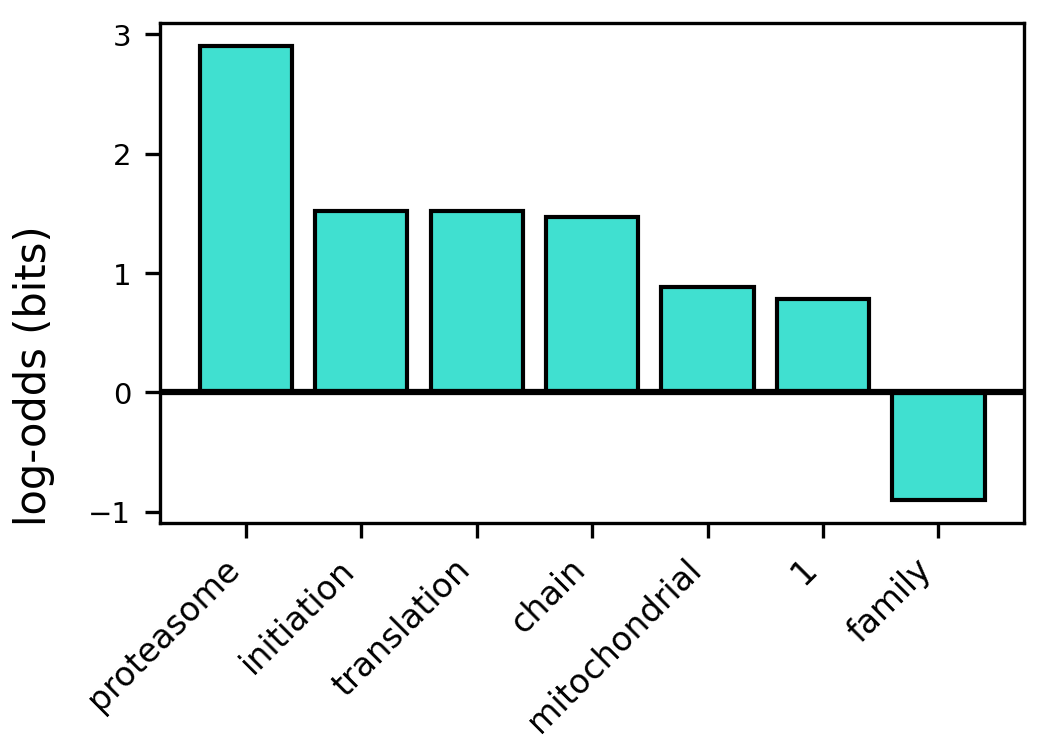

### greenyellow_kegg_path_level2_log_odds.png

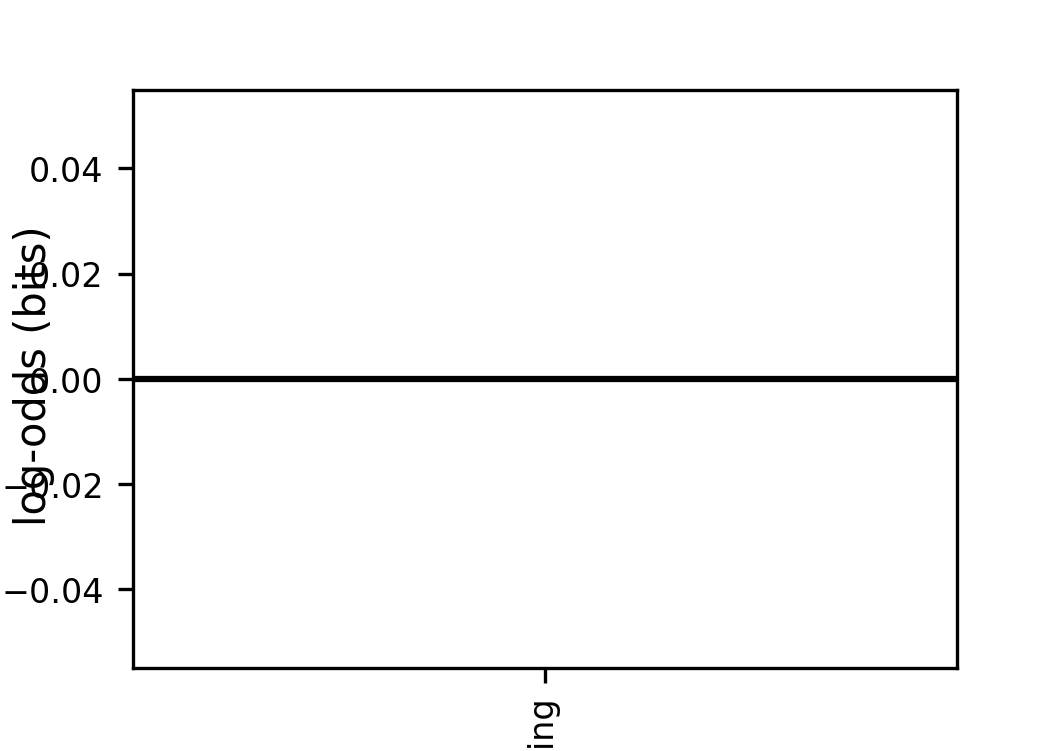

### i-ord_log_odds.png

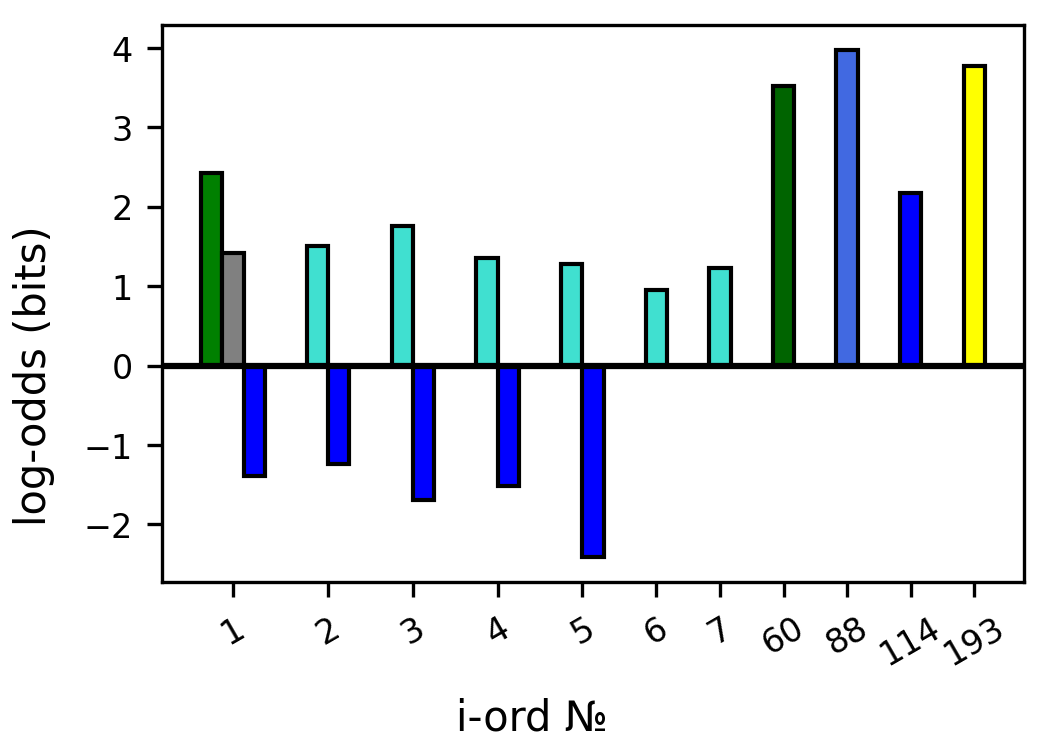

### i-start_log_odds.png

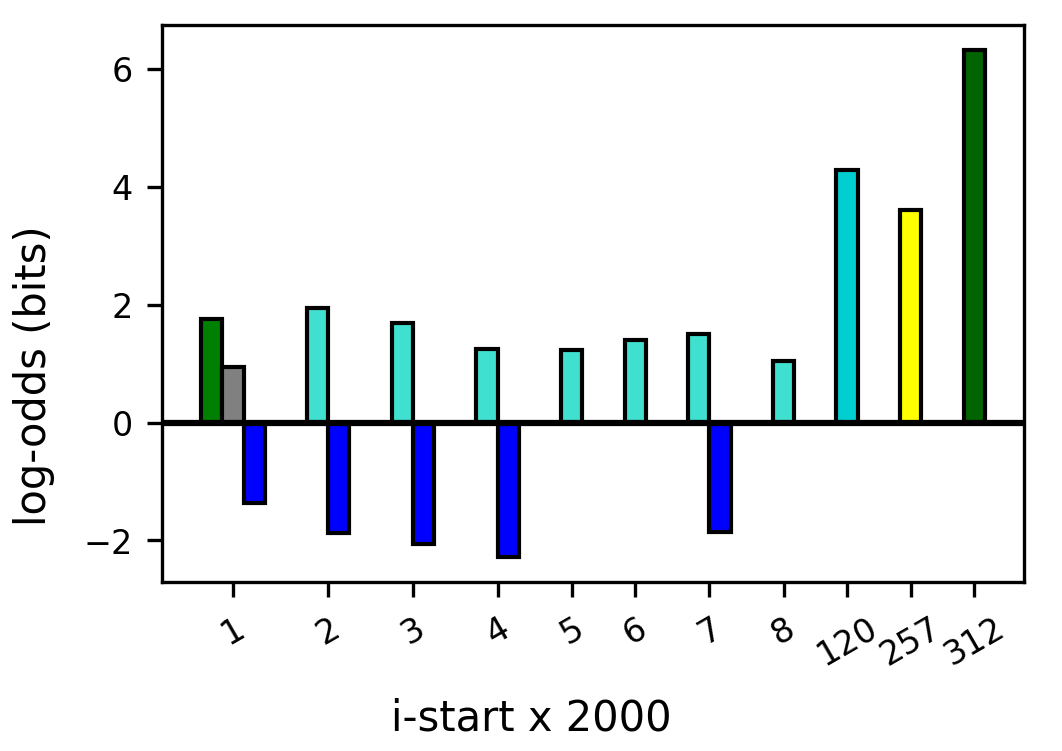

### kegg_path_level1_blue_log_odds.png

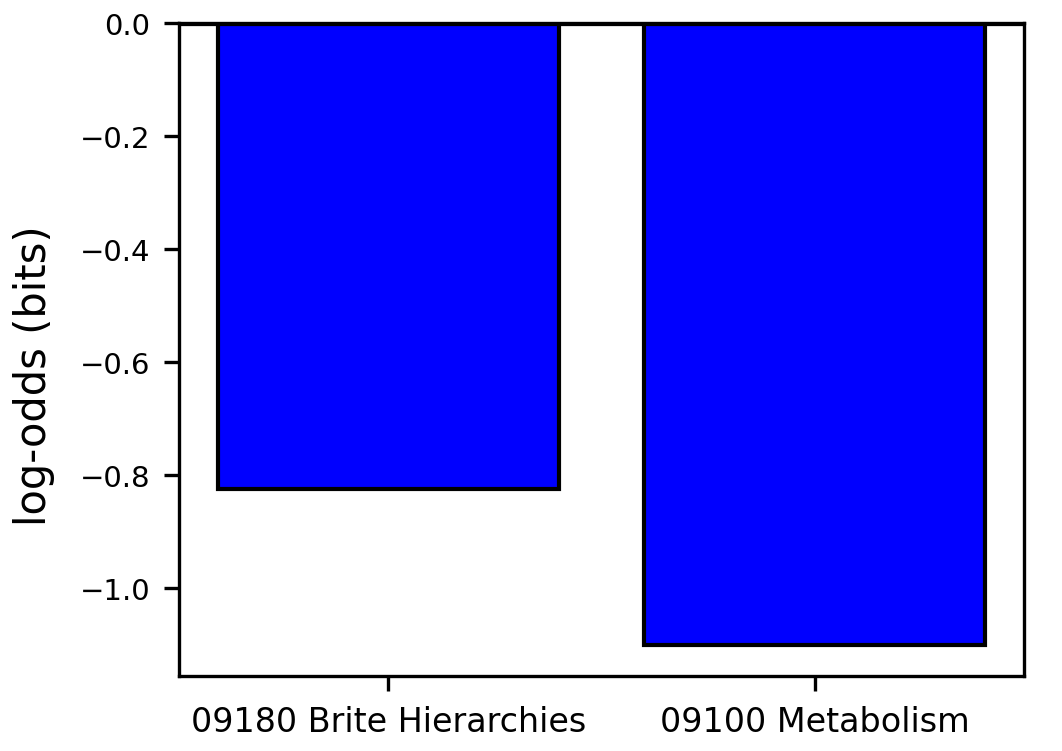

### kegg_path_level1_lightgreen_log_odds.png

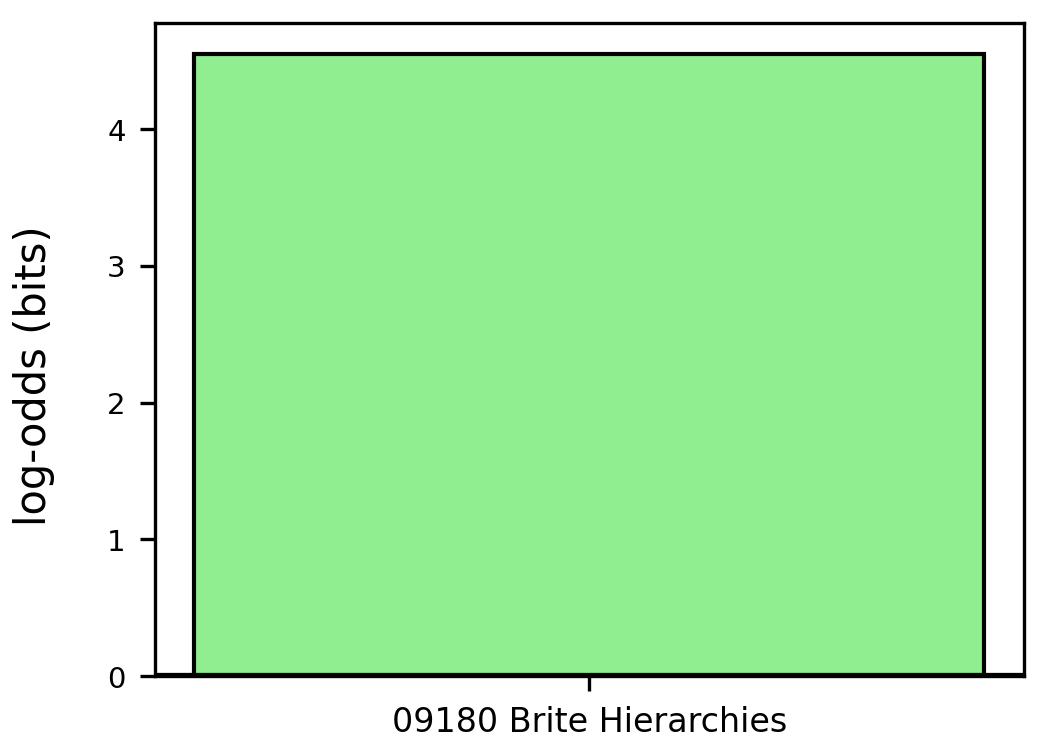

### kegg_path_level1_log_odds.png

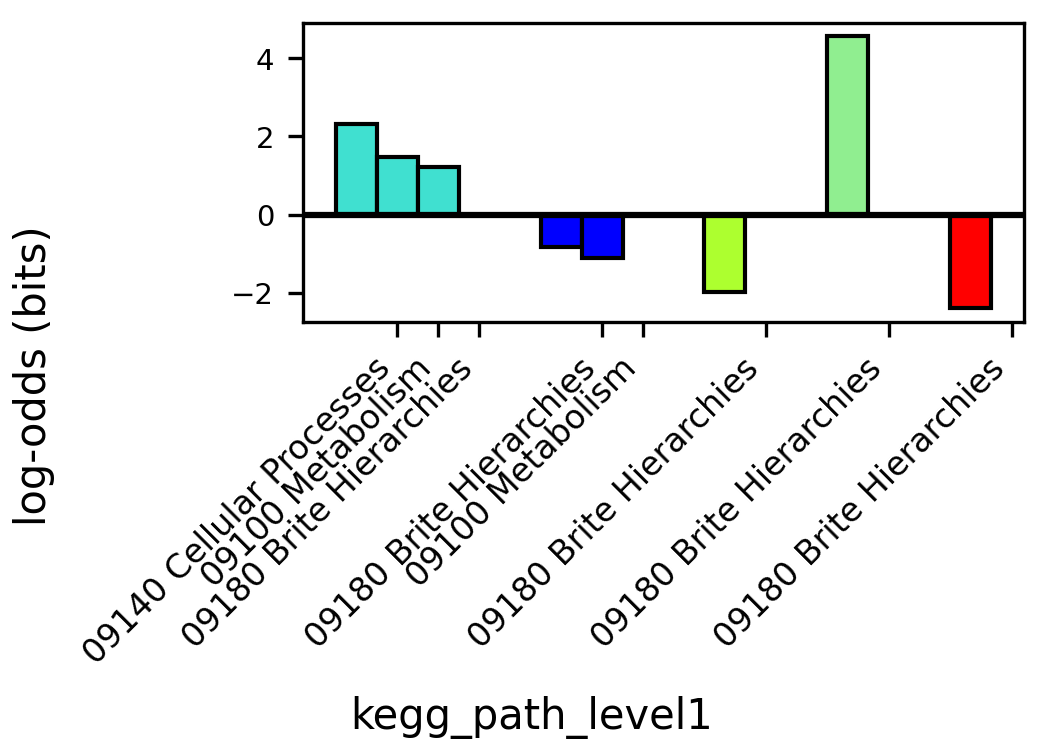

### kegg_path_level1_num.png

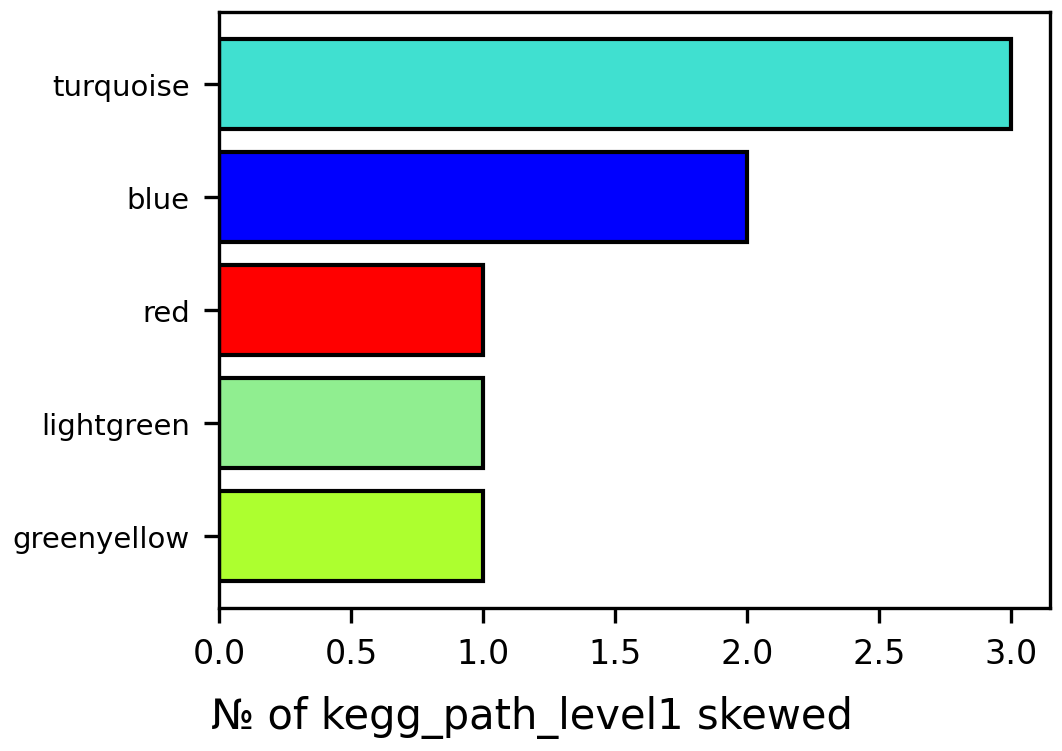

### kegg_path_level1_red_log_odds.png

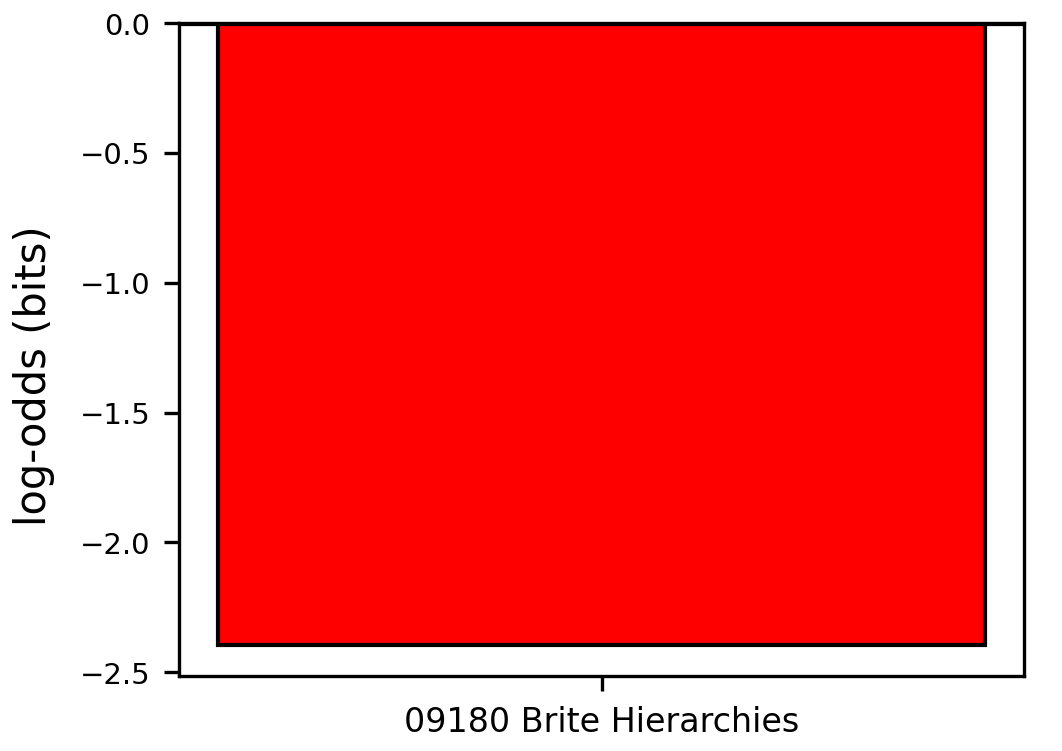

### kegg_path_level2_blue_log_odds.png

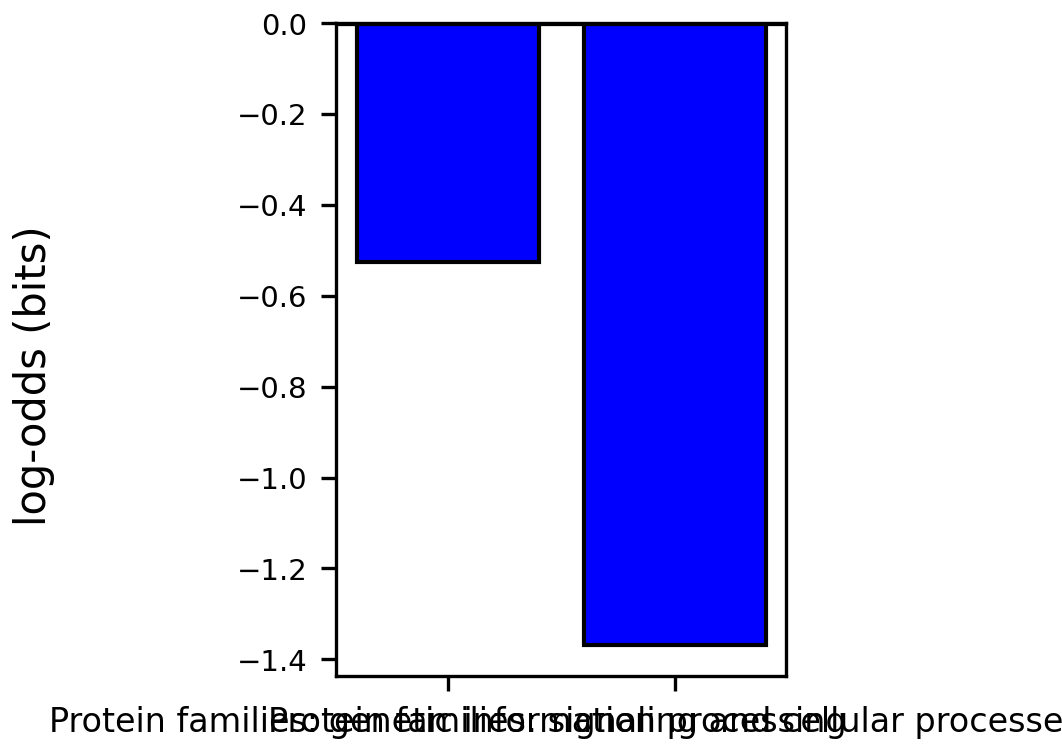

### kegg_path_level2_green_log_odds.png

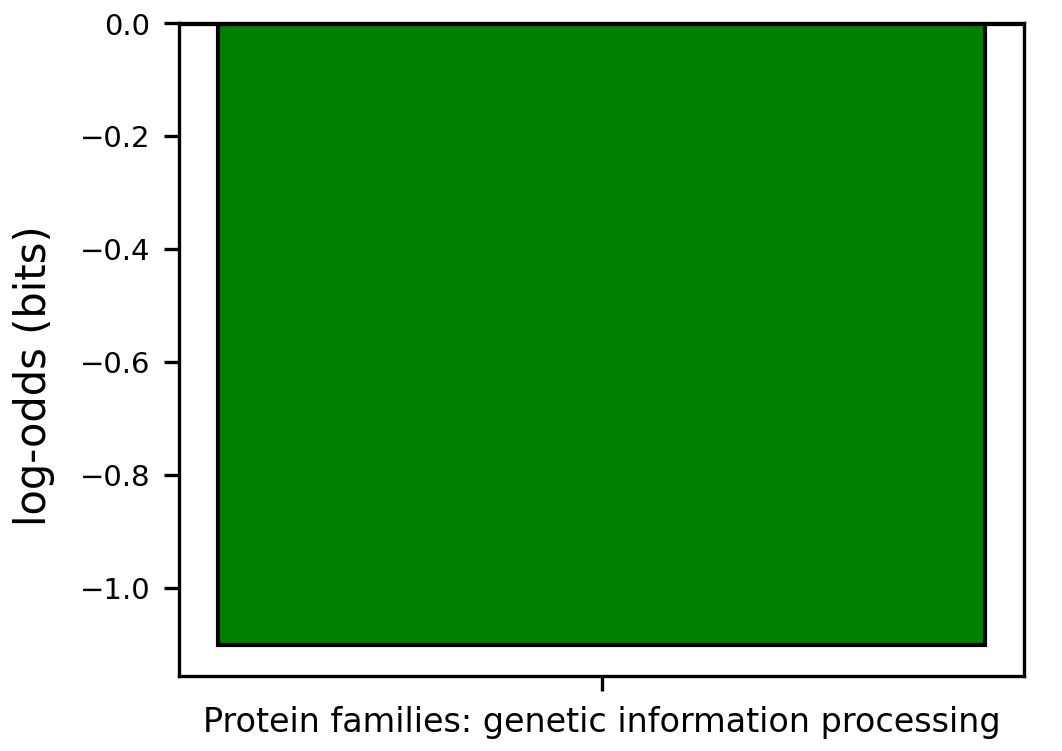

### kegg_path_level2_greenyellow_log_odds.png

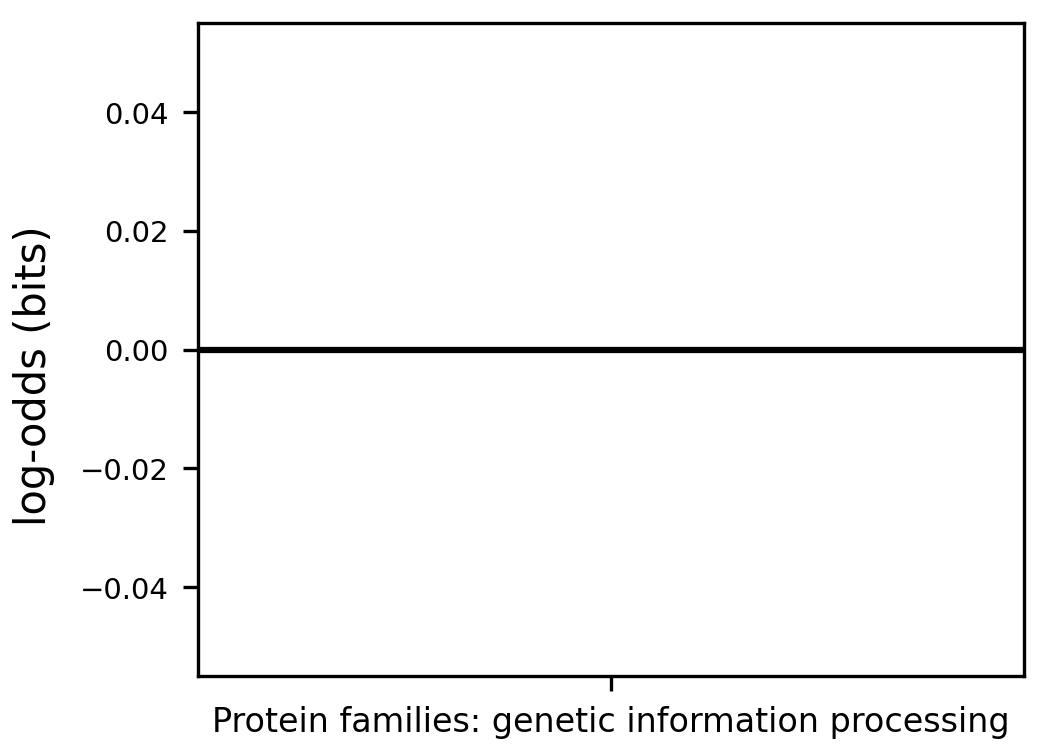

### kegg_path_level2_lightgreen_log_odds.png

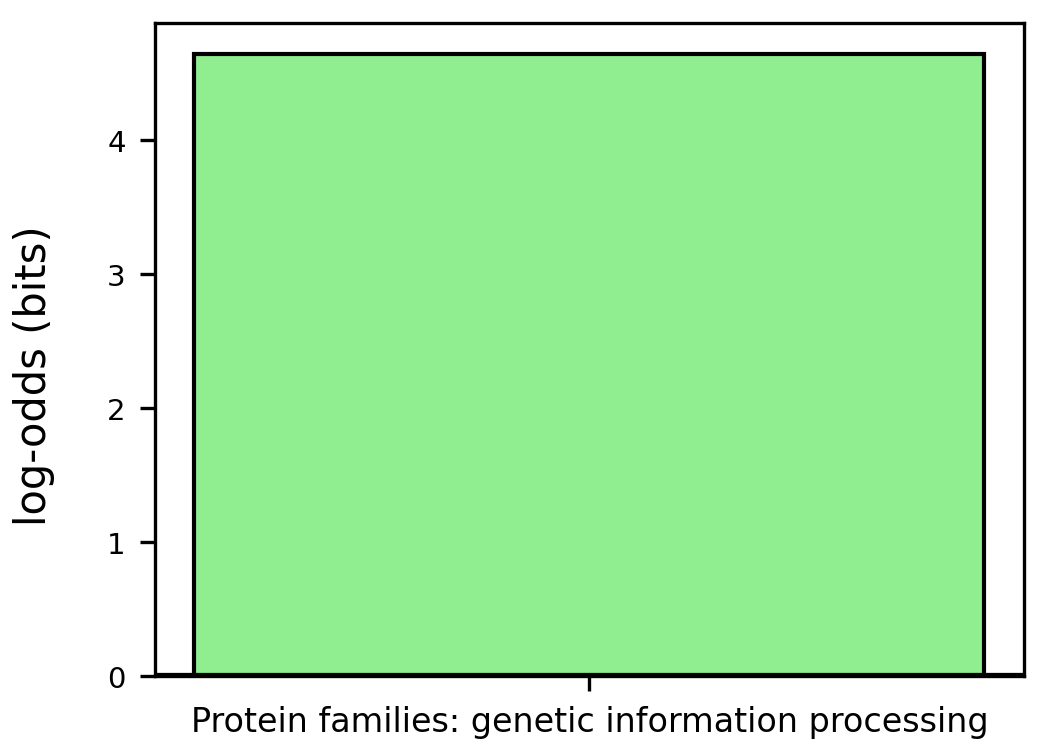

### kegg_path_level2_log_odds.png

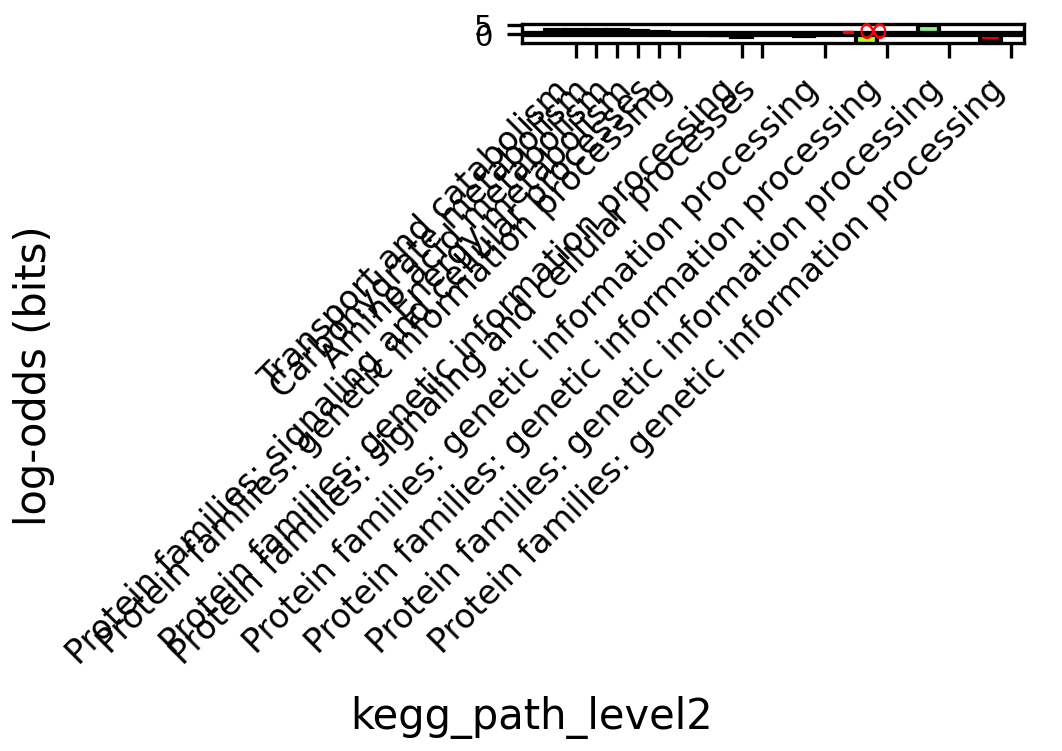

### kegg_path_level2_num.png

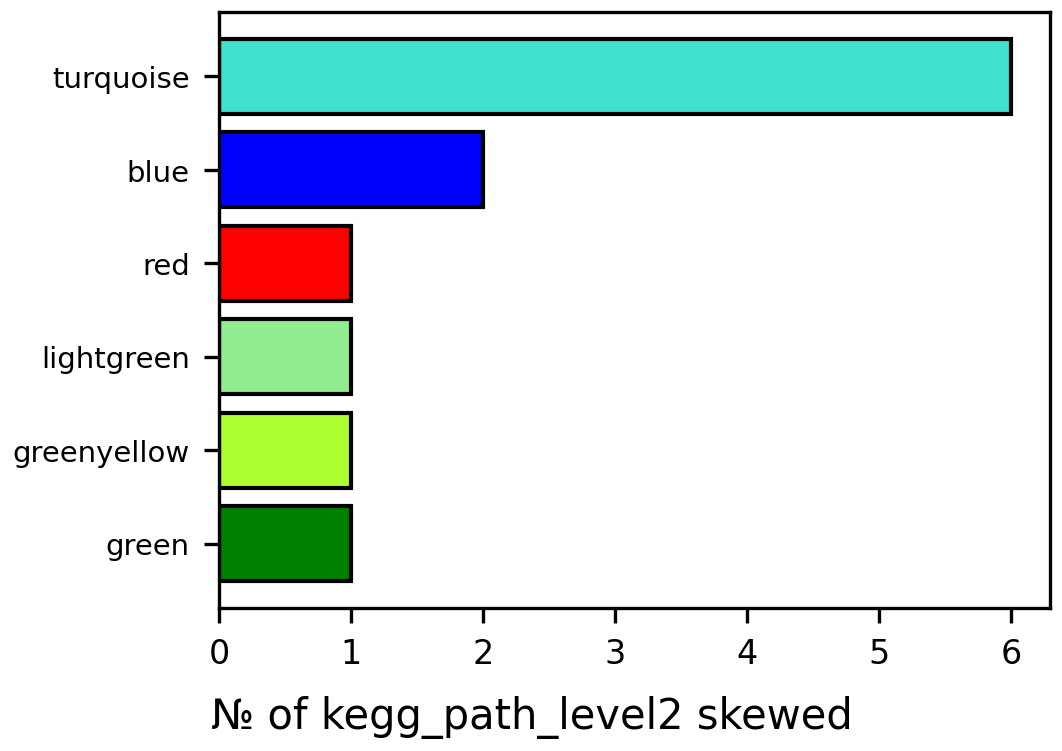

### kegg_path_level2_red_log_odds.png

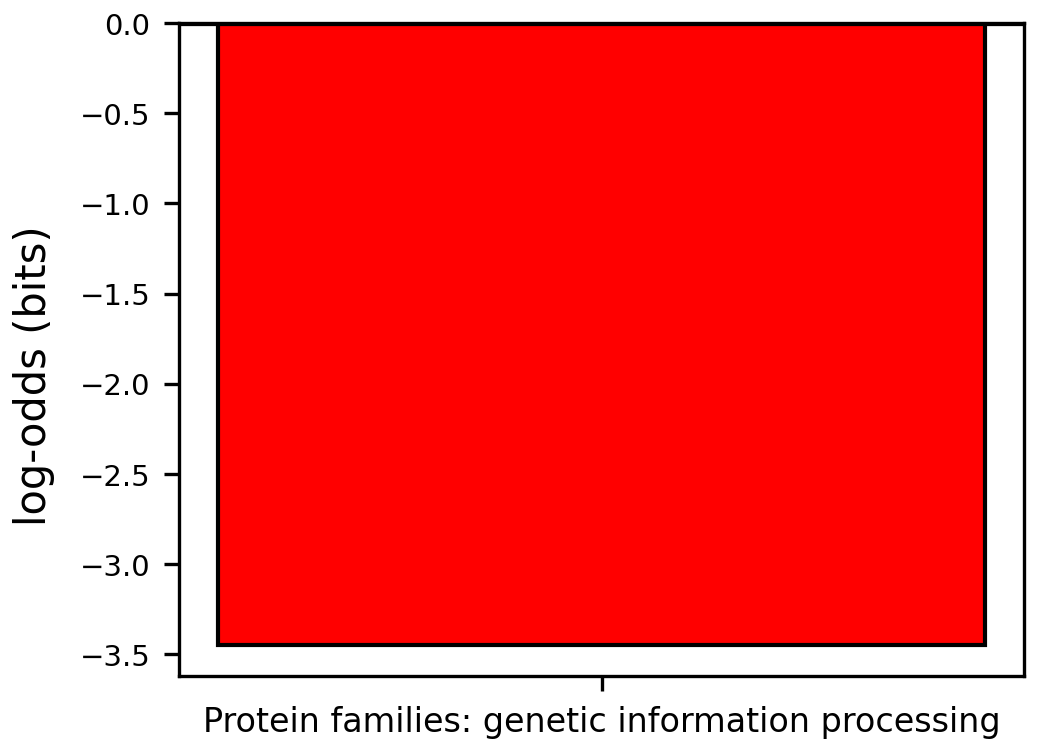

### kegg_path_level2_turquoise_log_odds.png

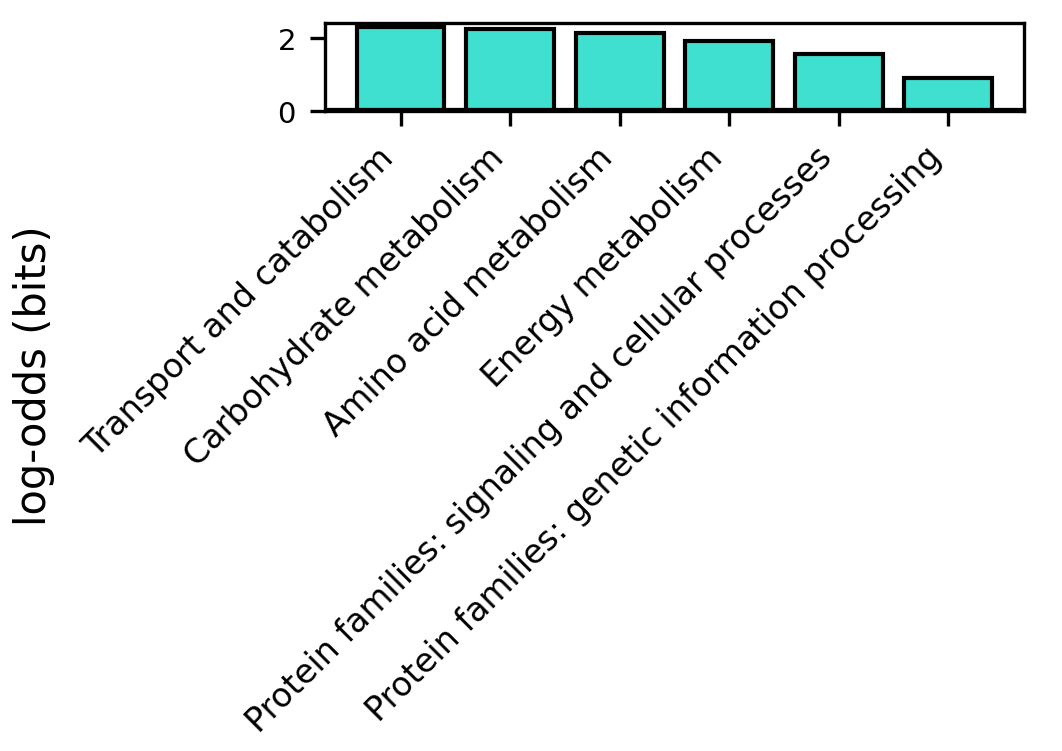

### kegg_path_level3_lightgreen_log_odds.png

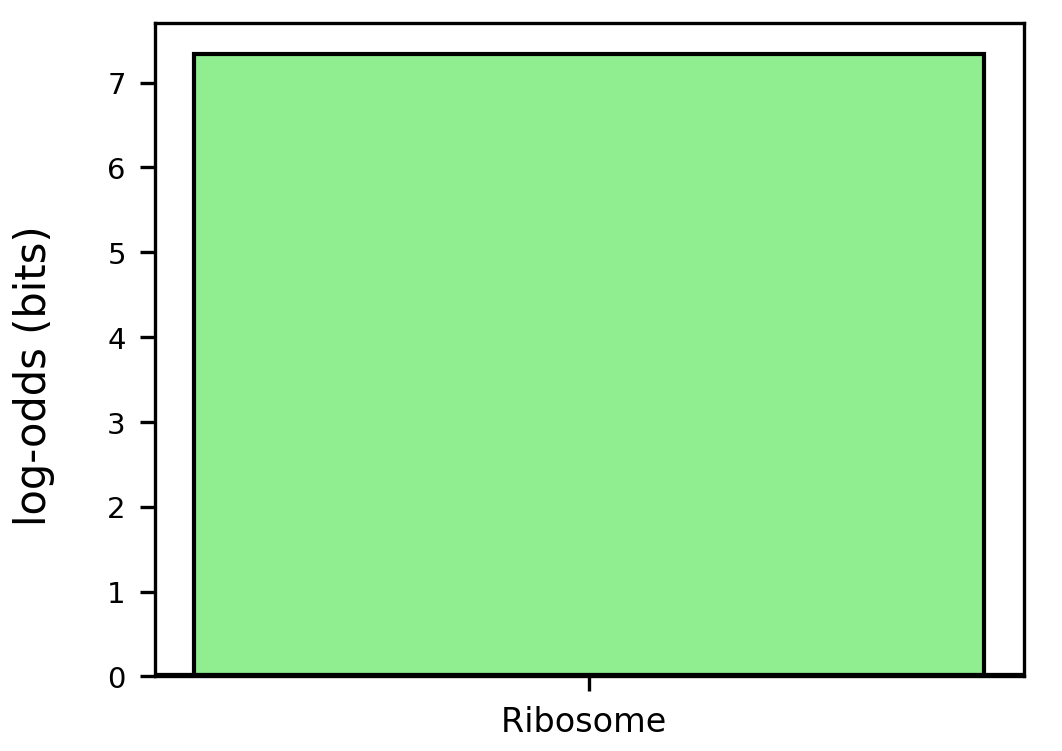

### kegg_path_level3_log_odds.png

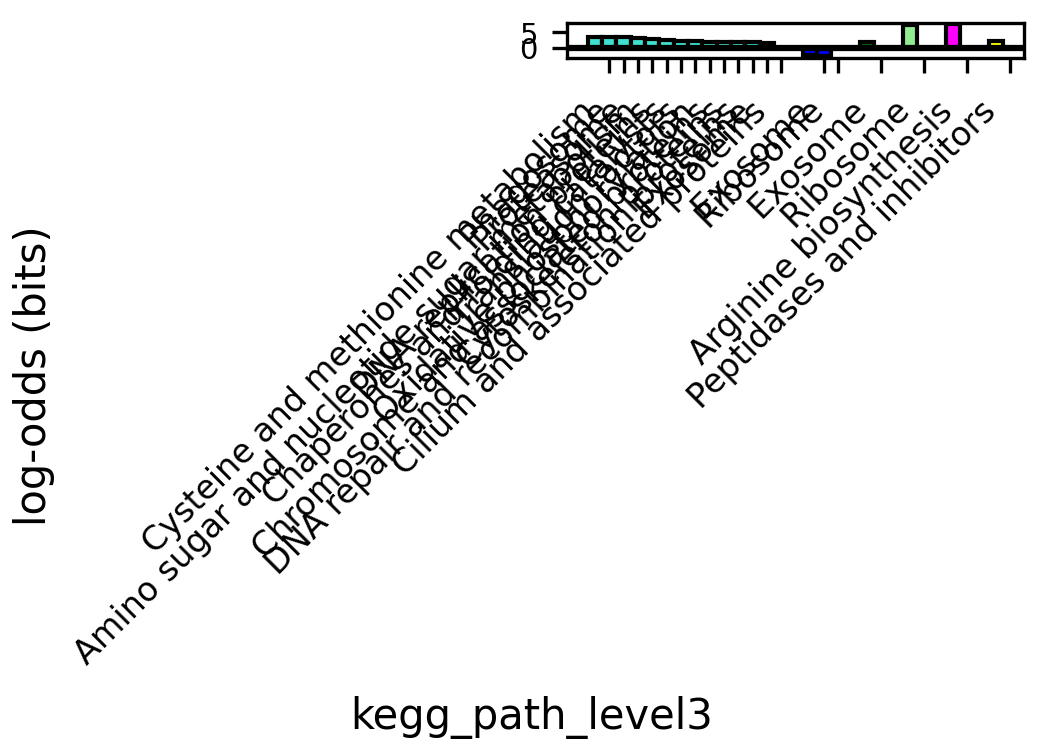

### kegg_path_level3_num.png

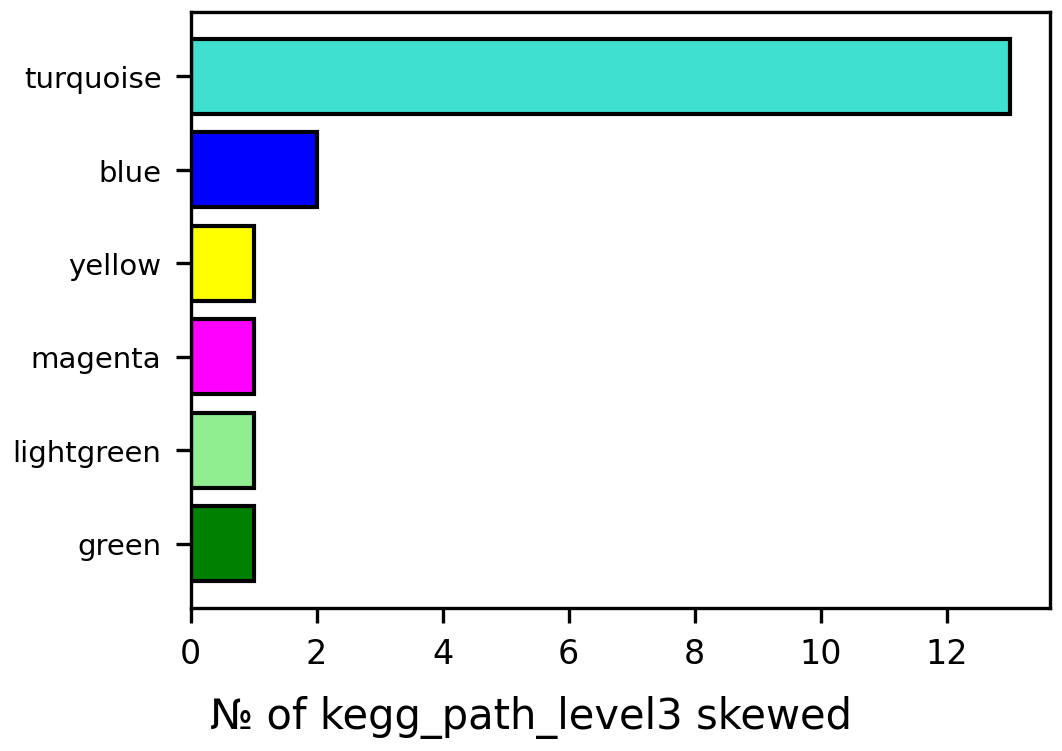

### kegg_path_level3_turquoise_log_odds.png

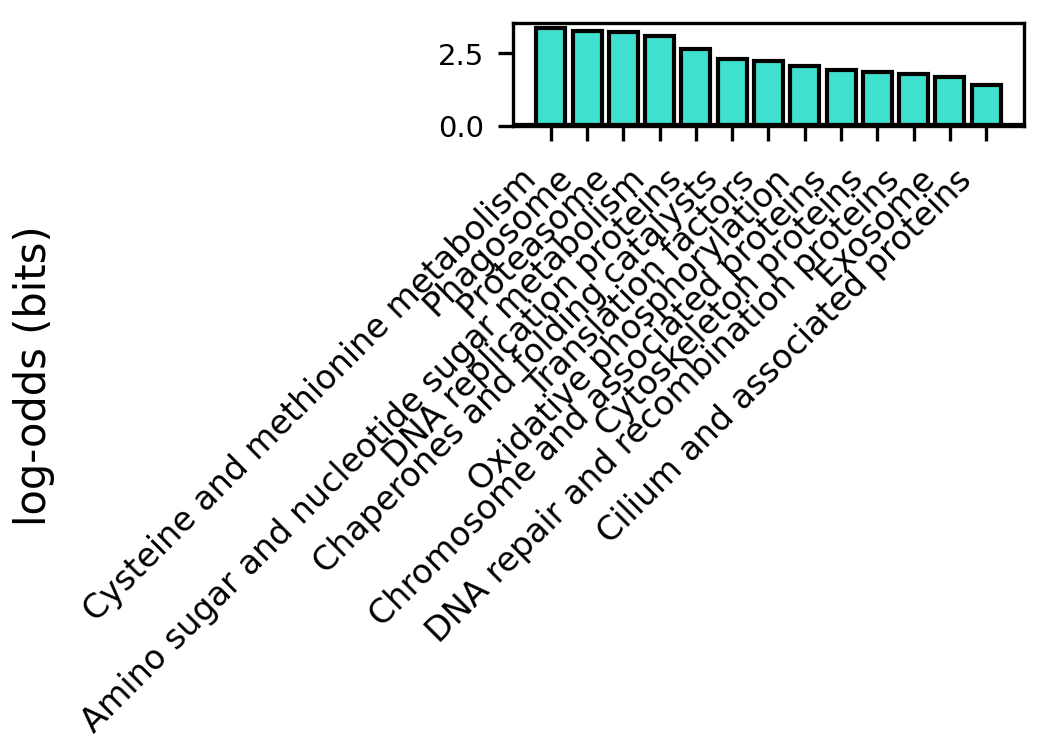

### kegg_path_level3_yellow_log_odds.png

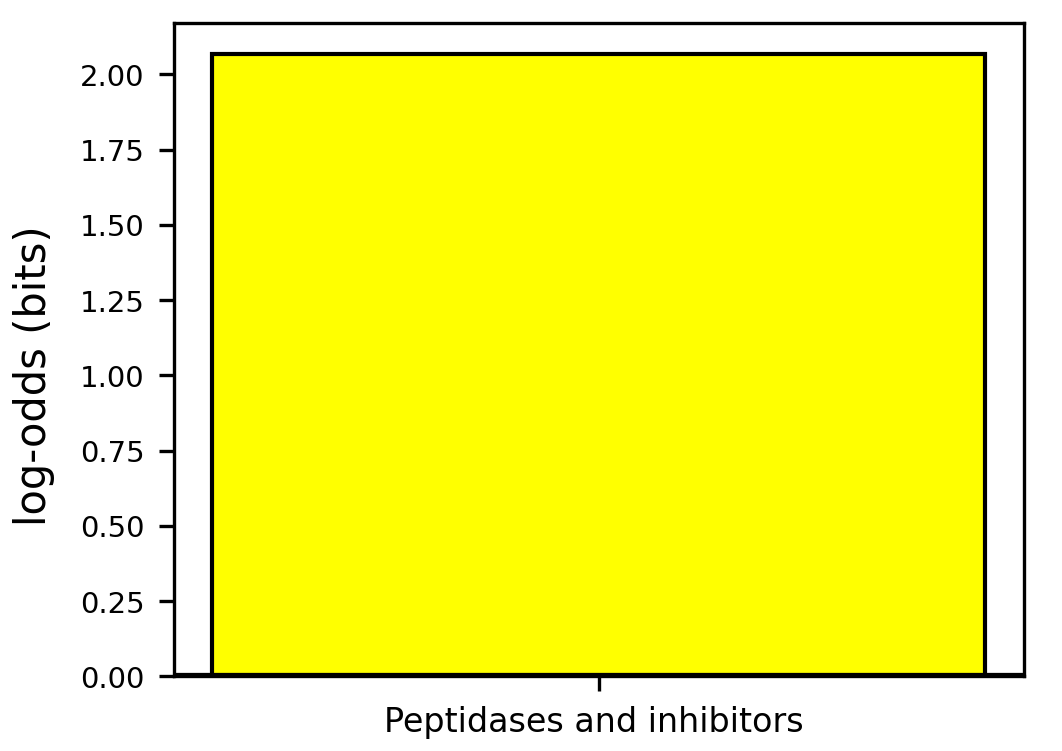

### located_in_log_odds.png

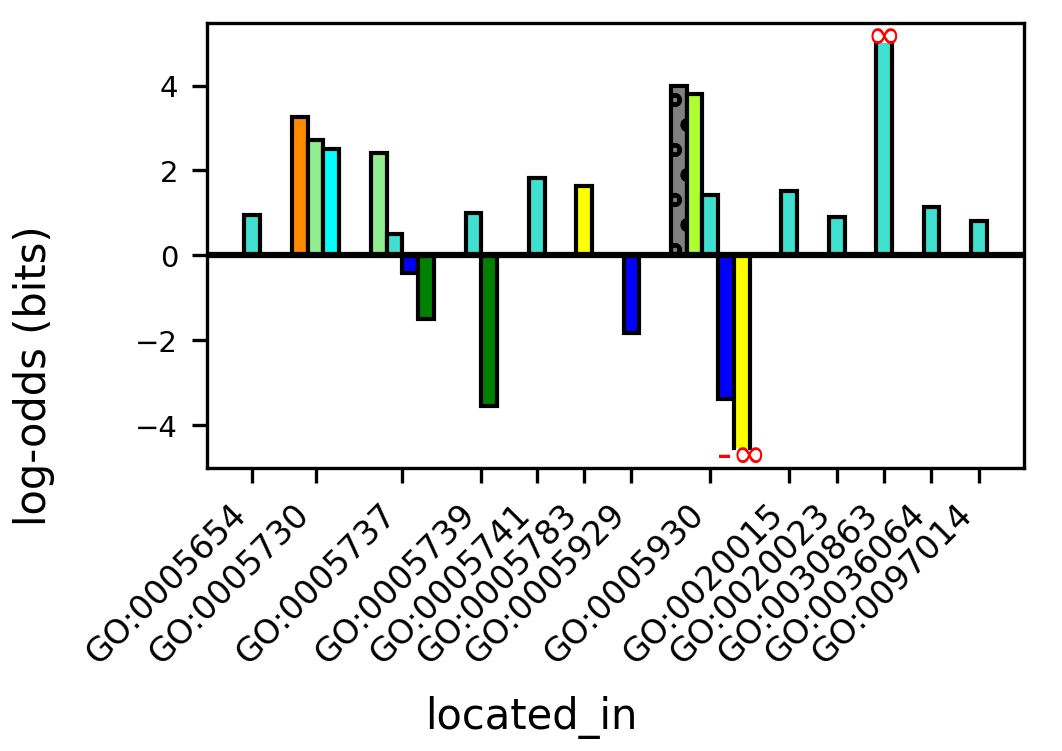

### motifs_black_log_odds.png

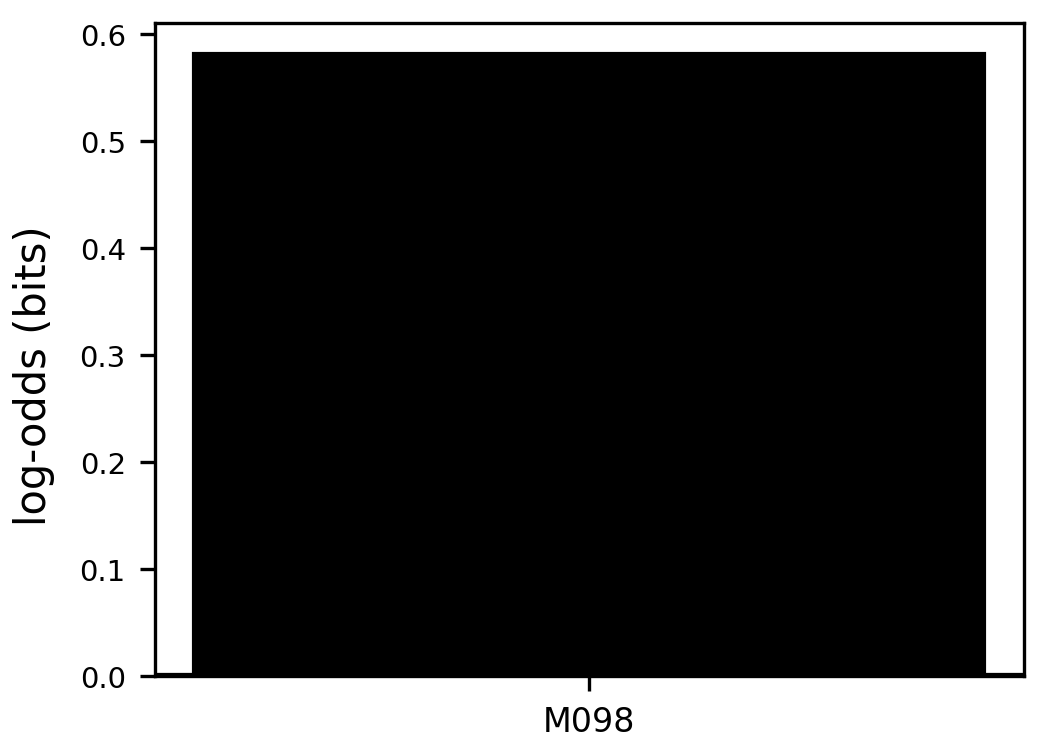

### motifs_blue_log_odds.png

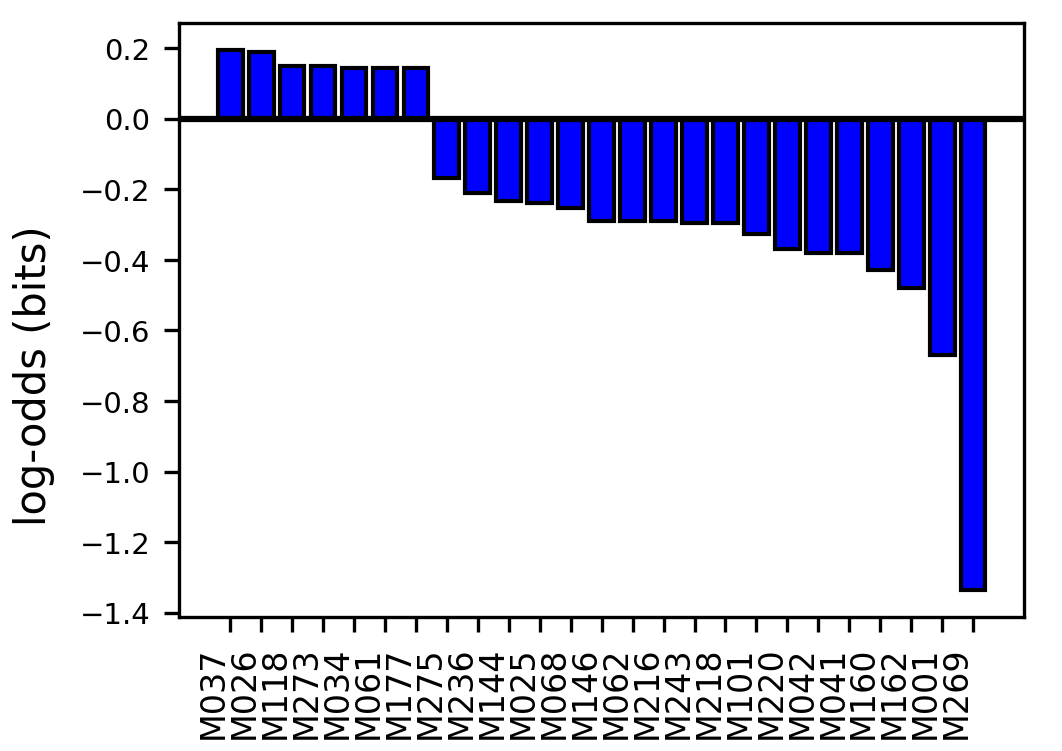

### motifs_brown_log_odds.png

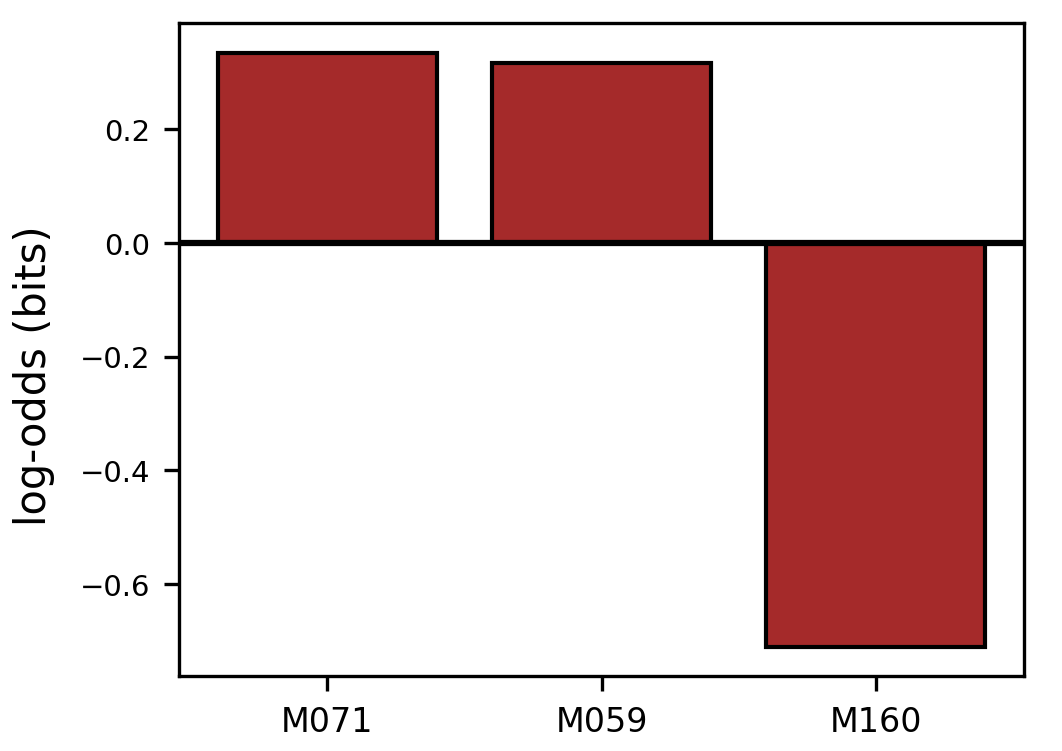

### motifs_cyan_log_odds.png

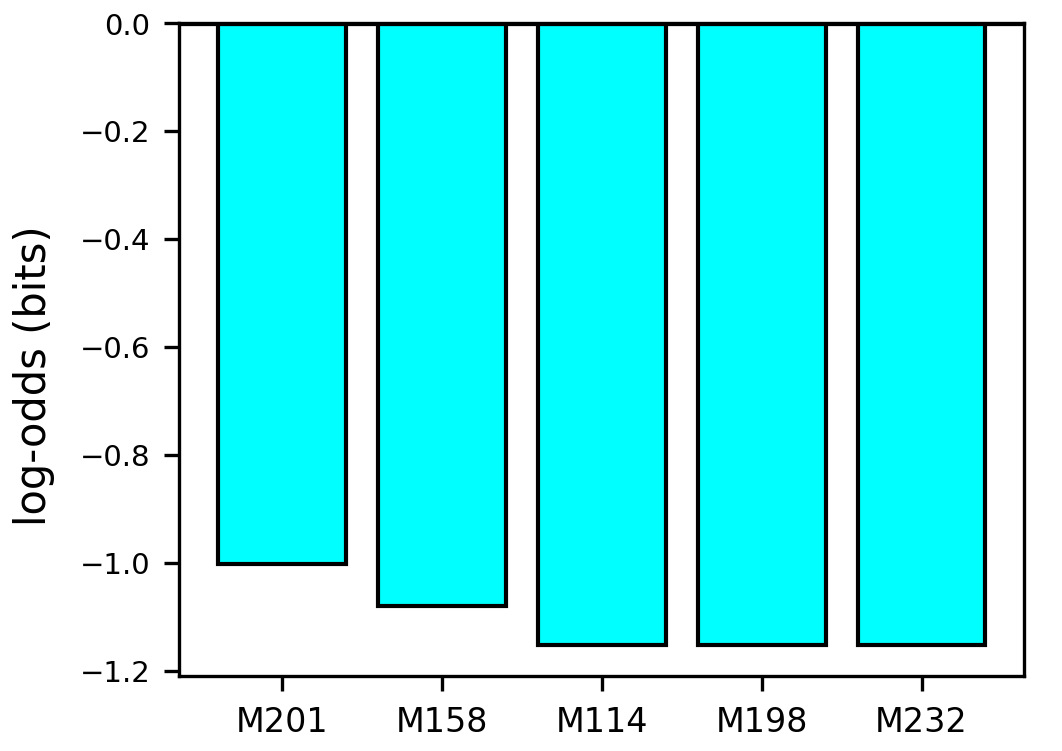

### motifs_darkgreen_log_odds.png

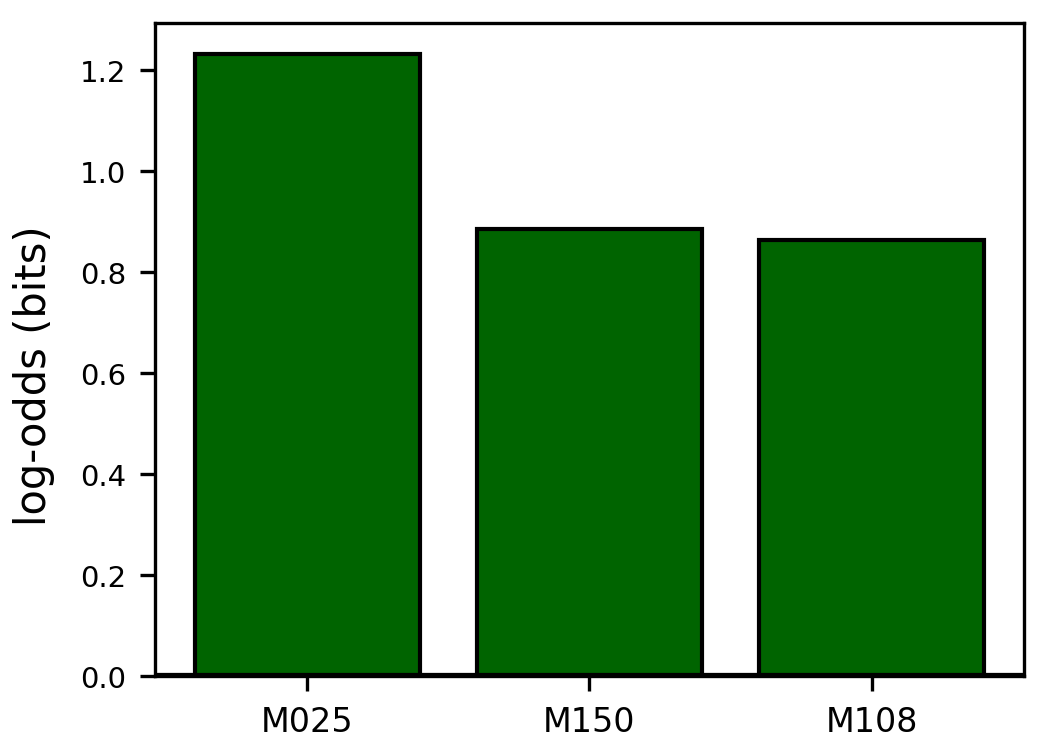

### motifs_darkgrey_log_odds.png

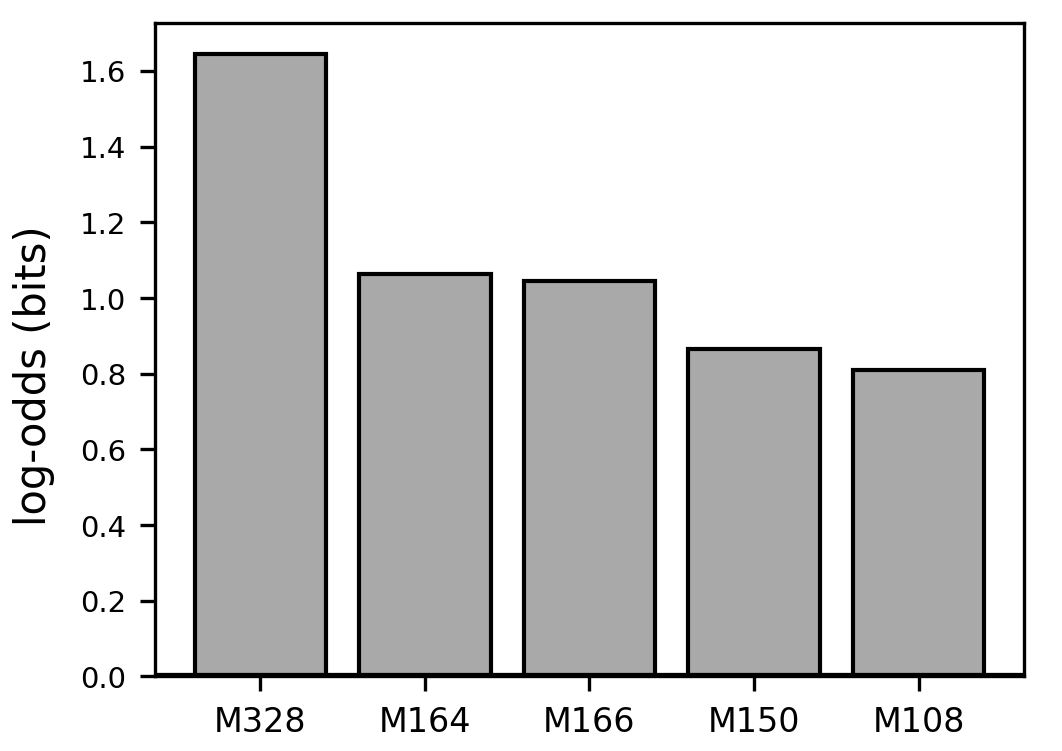

### motifs_darkorange_log_odds.png

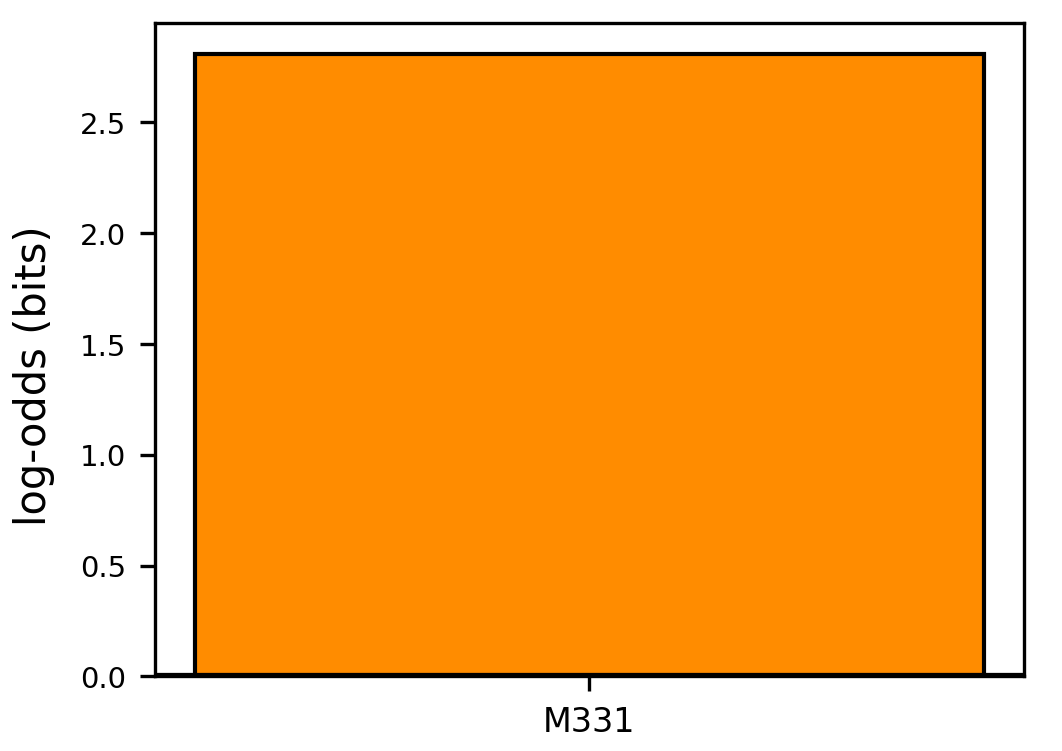

### motifs_green_log_odds.png

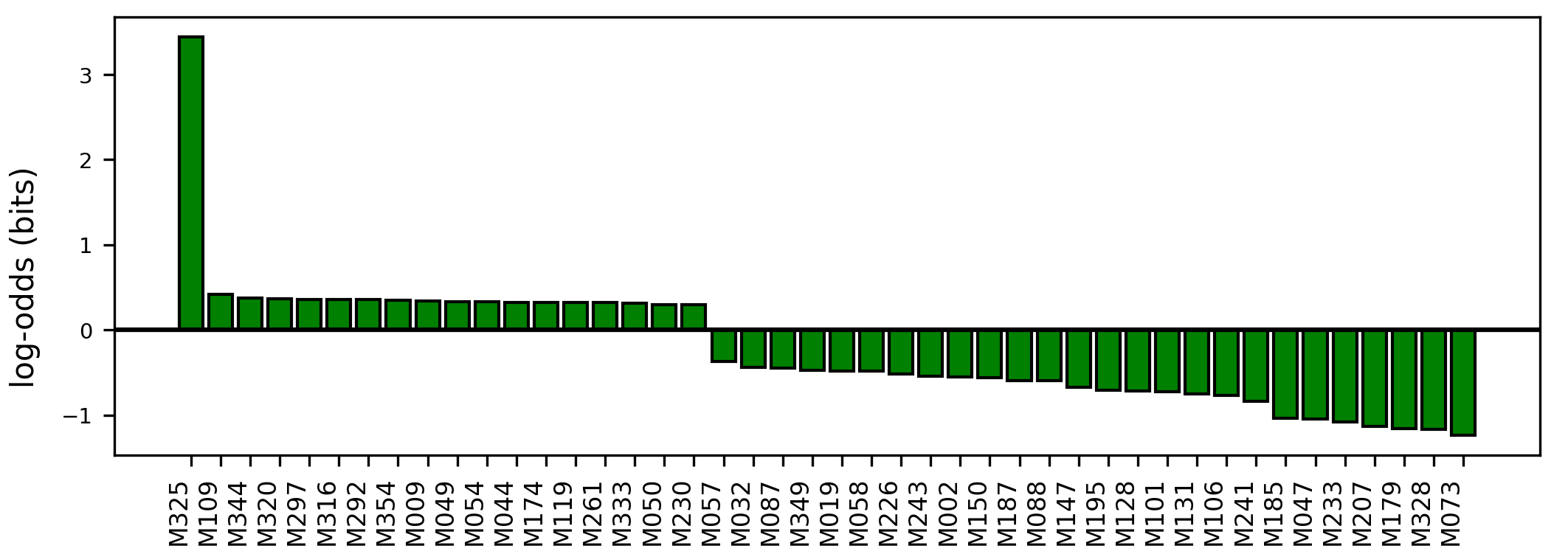

### motifs_greenyellow_log_odds.png

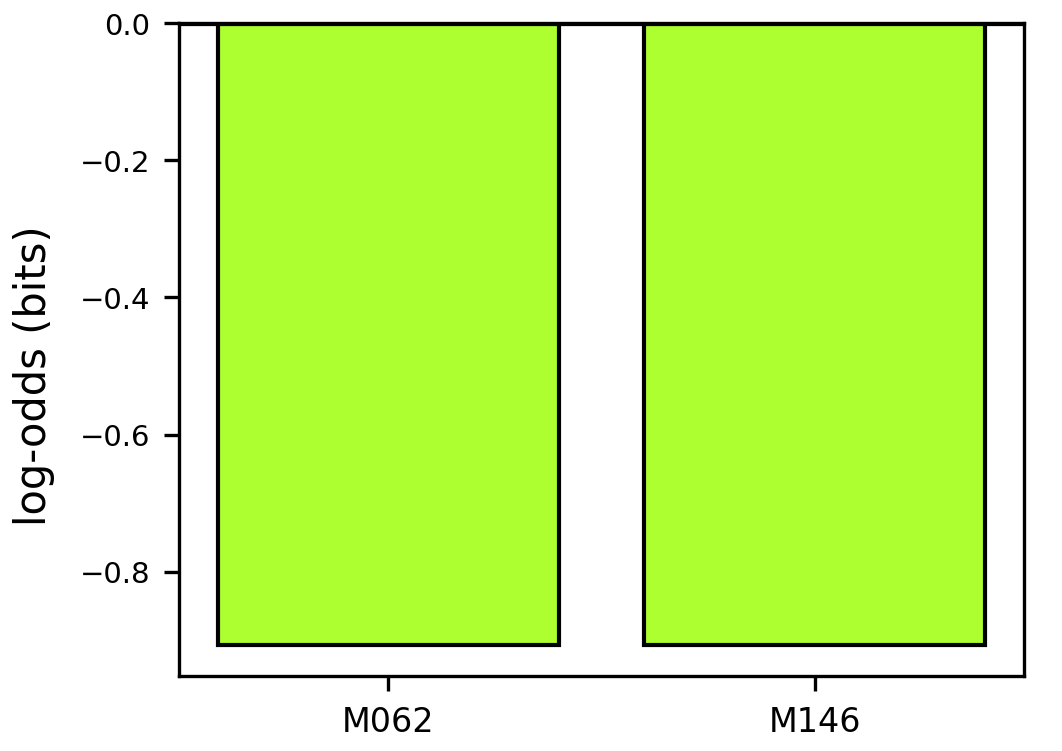

### motifs_grey60_log_odds.png

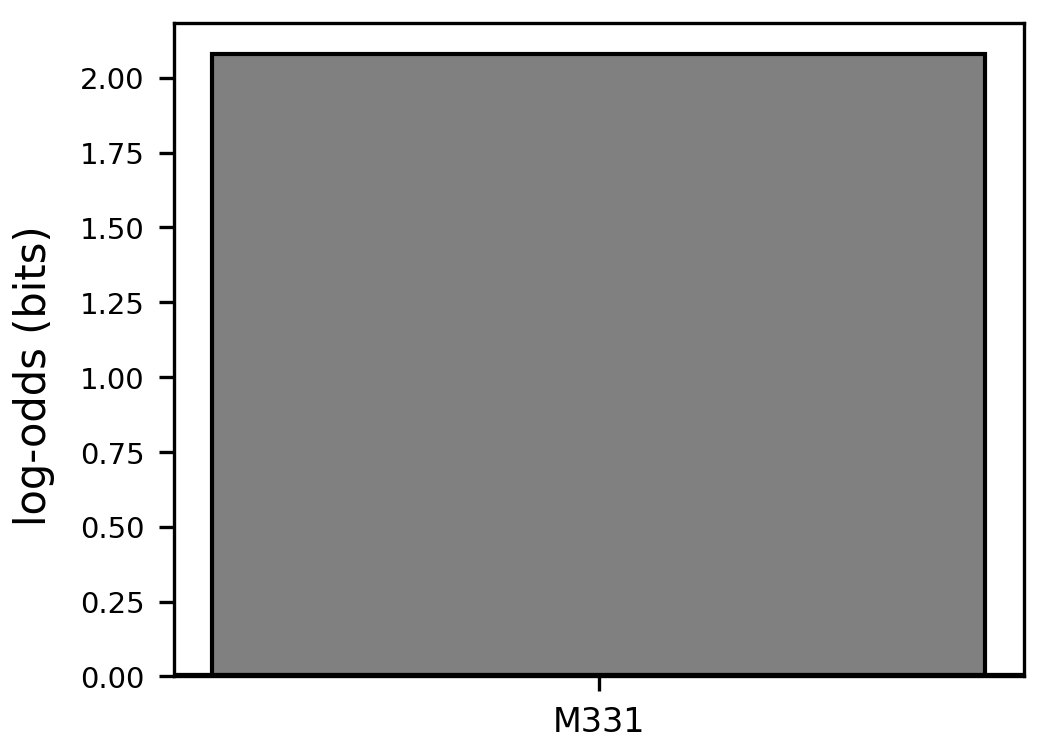

### motifs_grey_log_odds.png

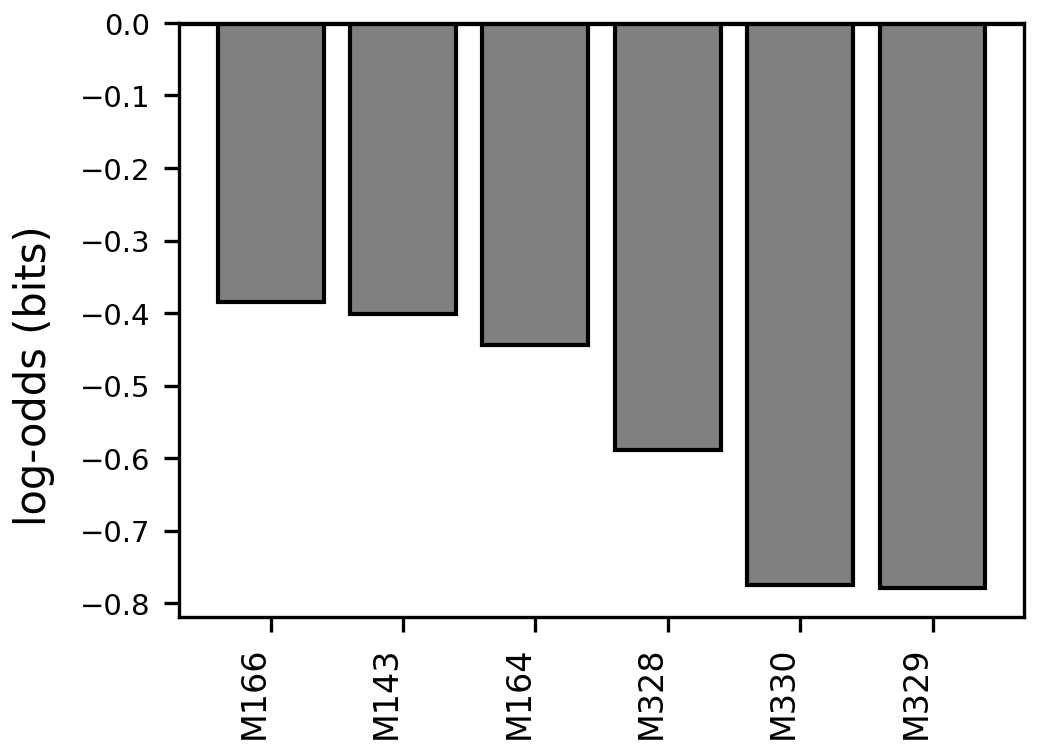

### motifs_lightcyan_log_odds.png

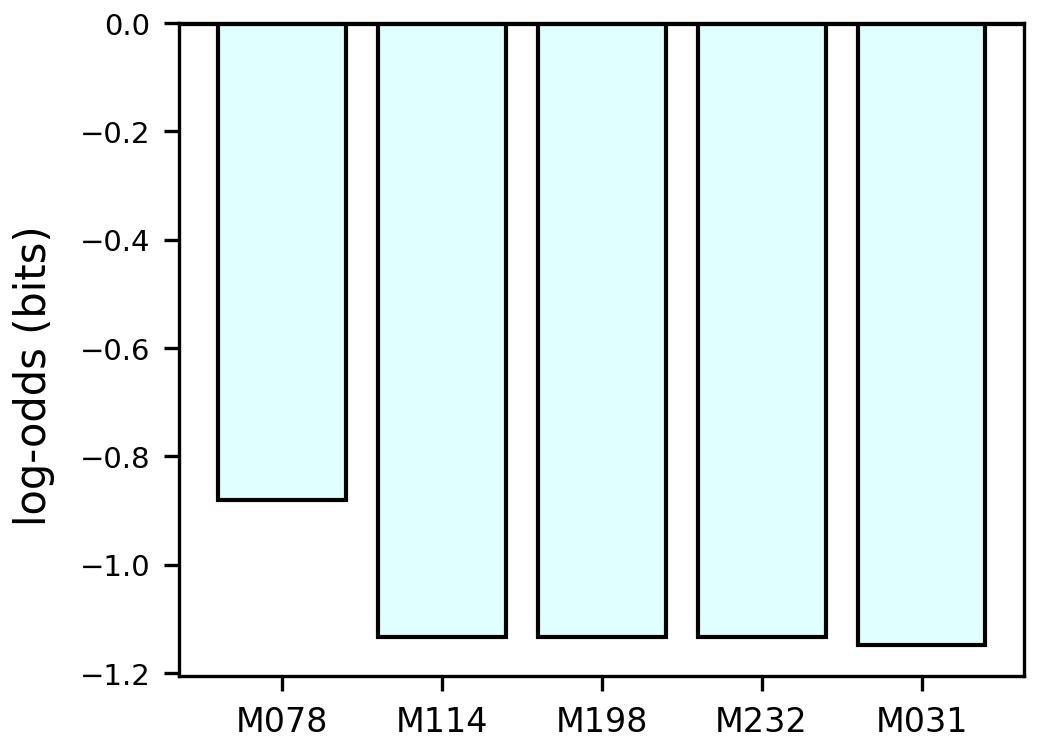

### motifs_lightgreen_log_odds.png

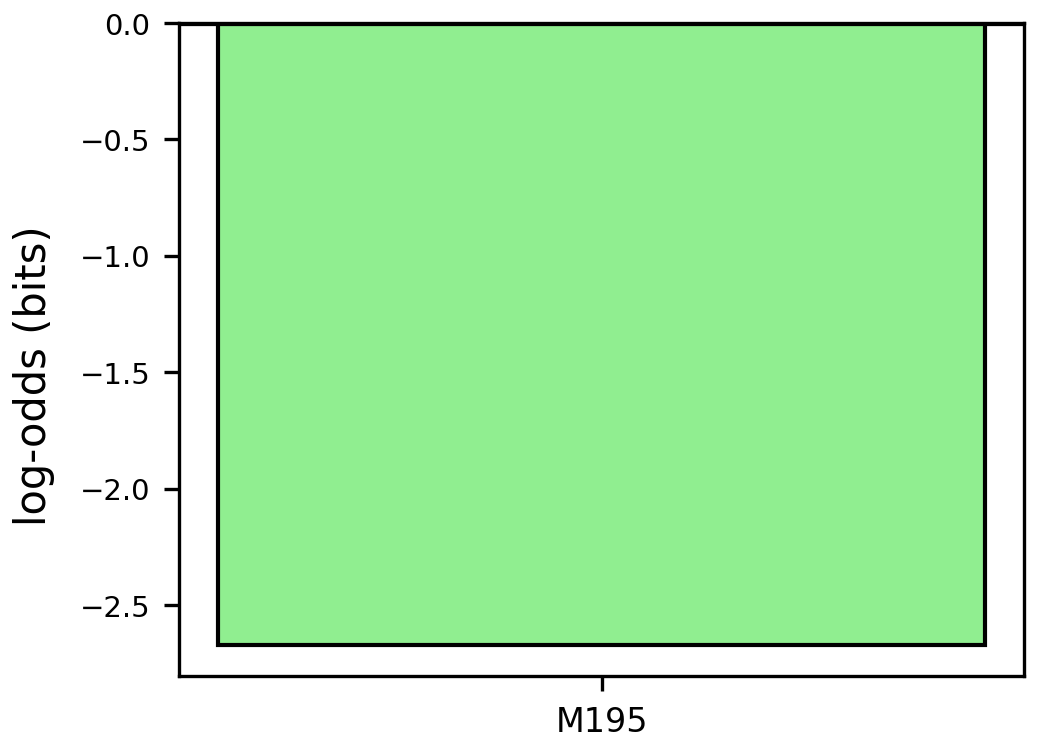

### motifs_log_odds.png

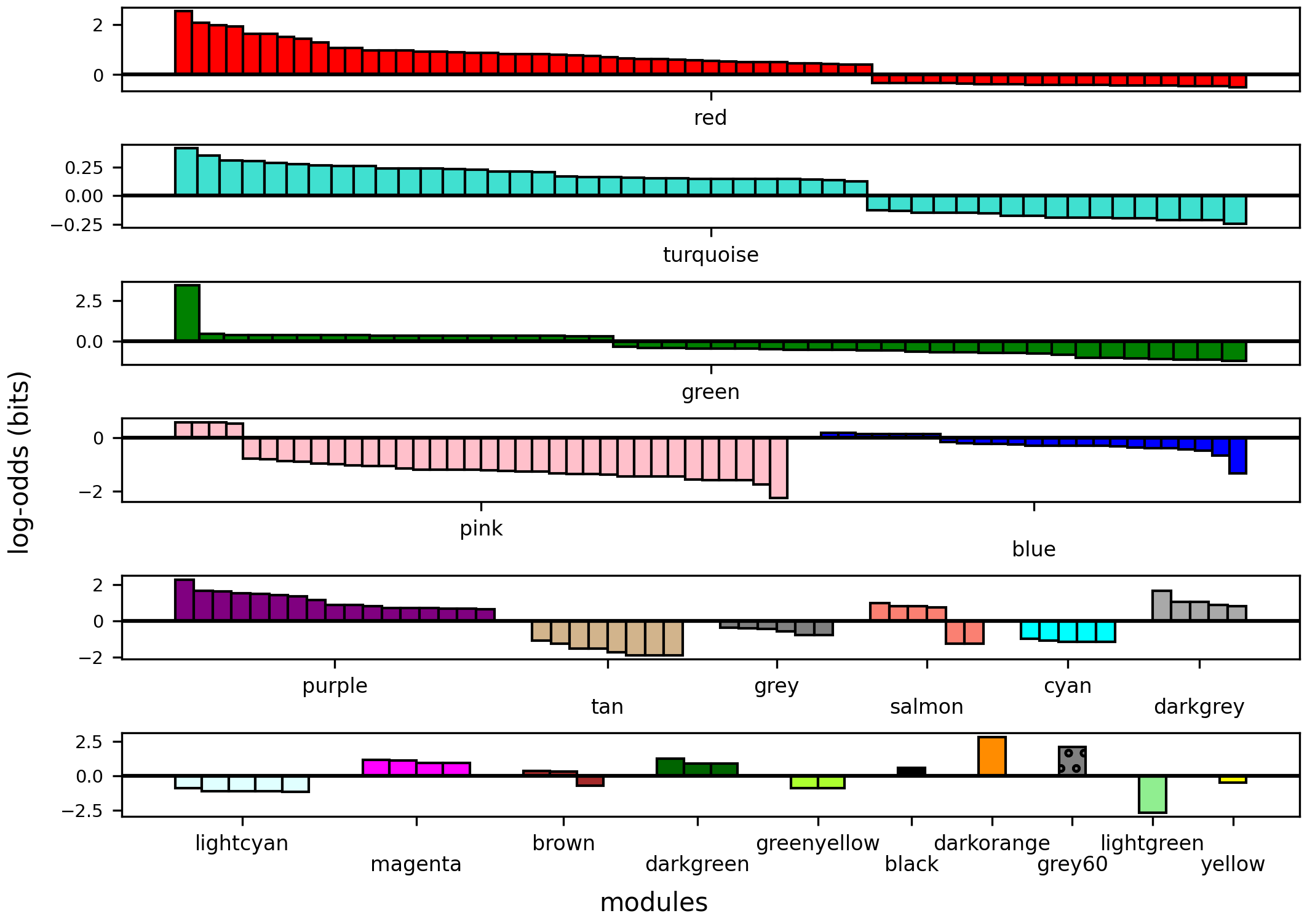

### motifs_magenta_log_odds.png

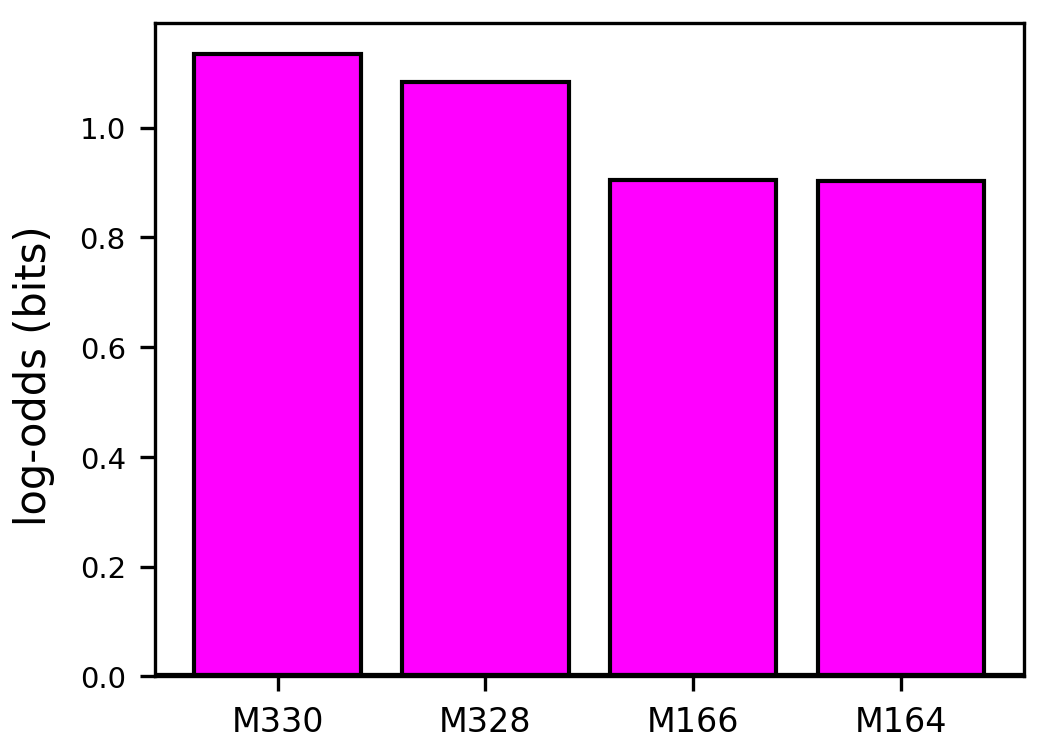

### motifs_num.png

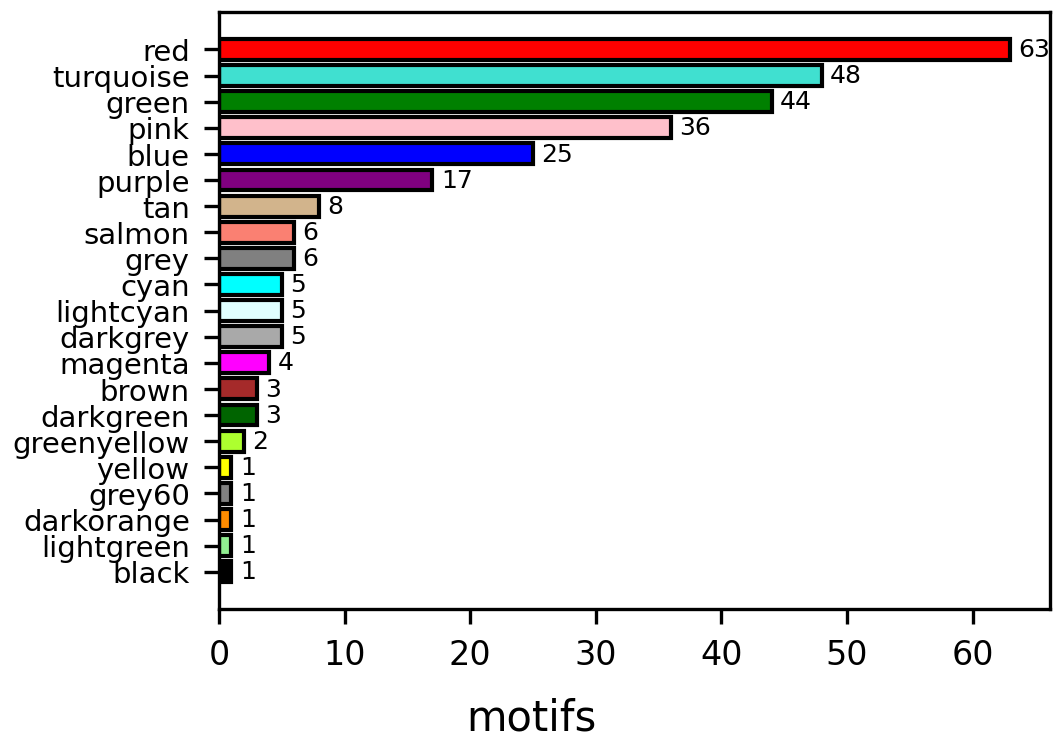

### motifs_pink_log_odds.png

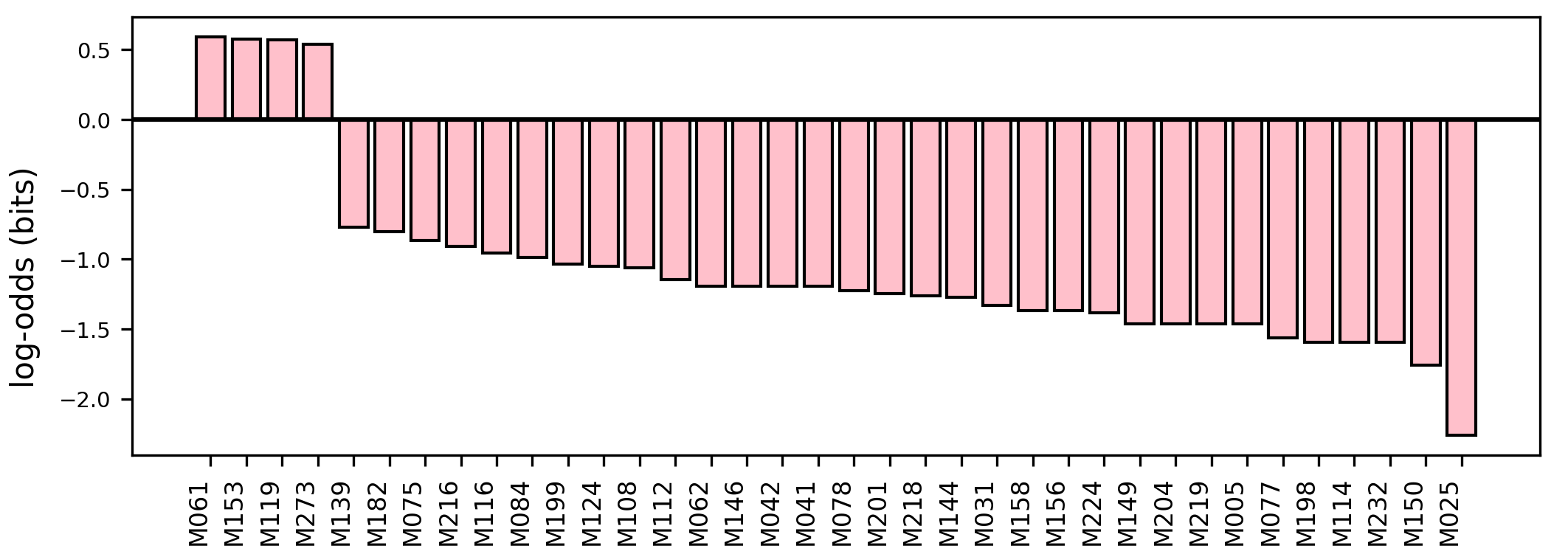

### motifs_purple_log_odds.png

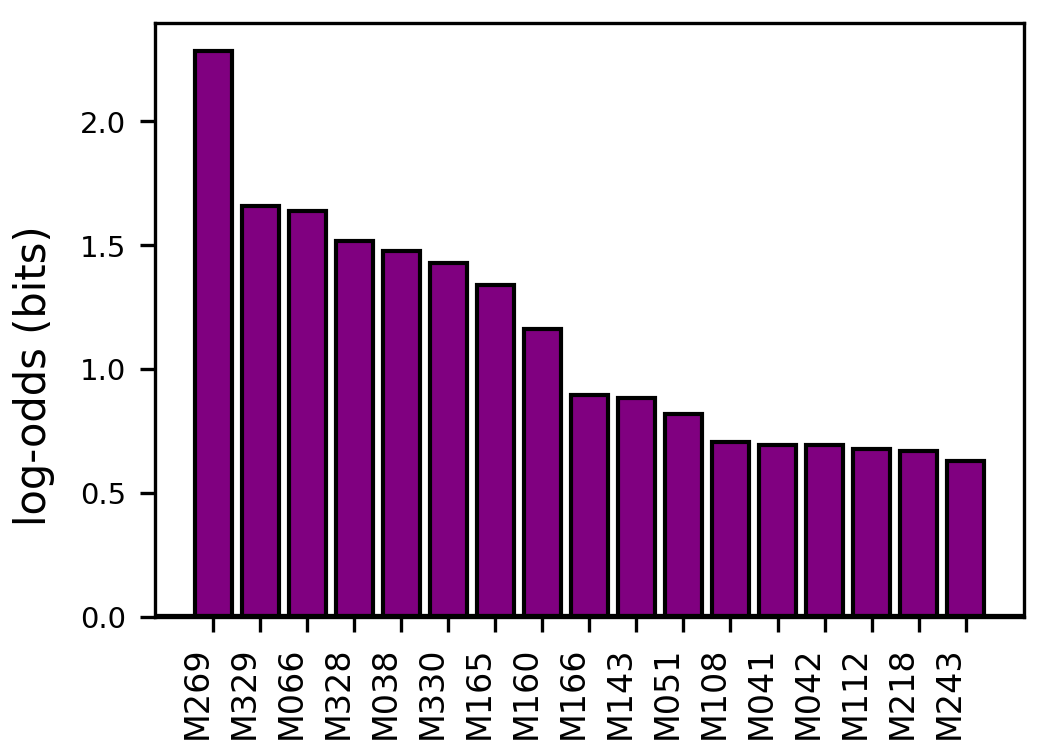

### motifs_red_log_odds.png

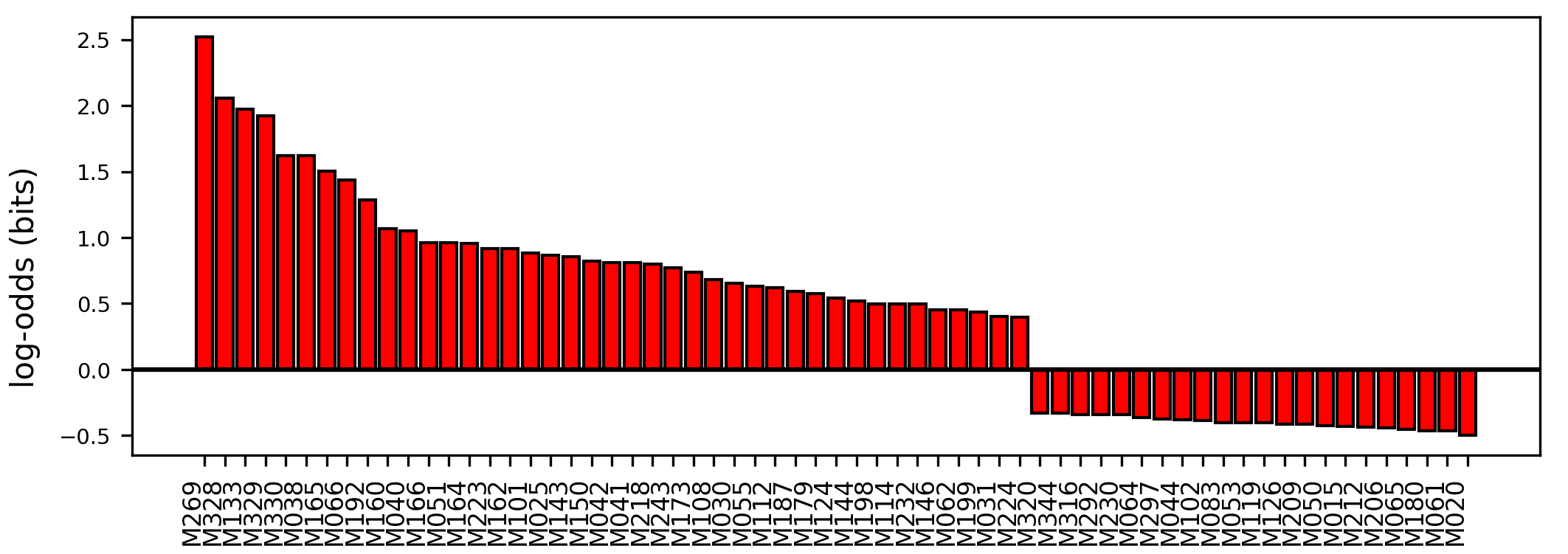

### motifs_salmon_log_odds.png

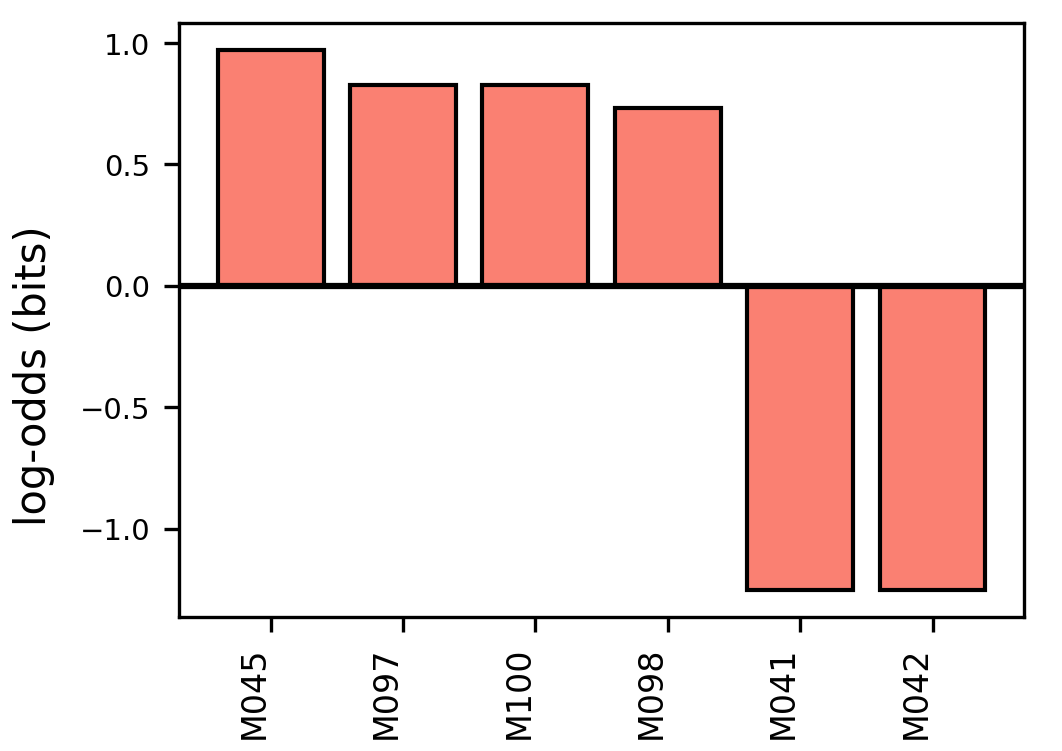

### motifs_tan_log_odds.png

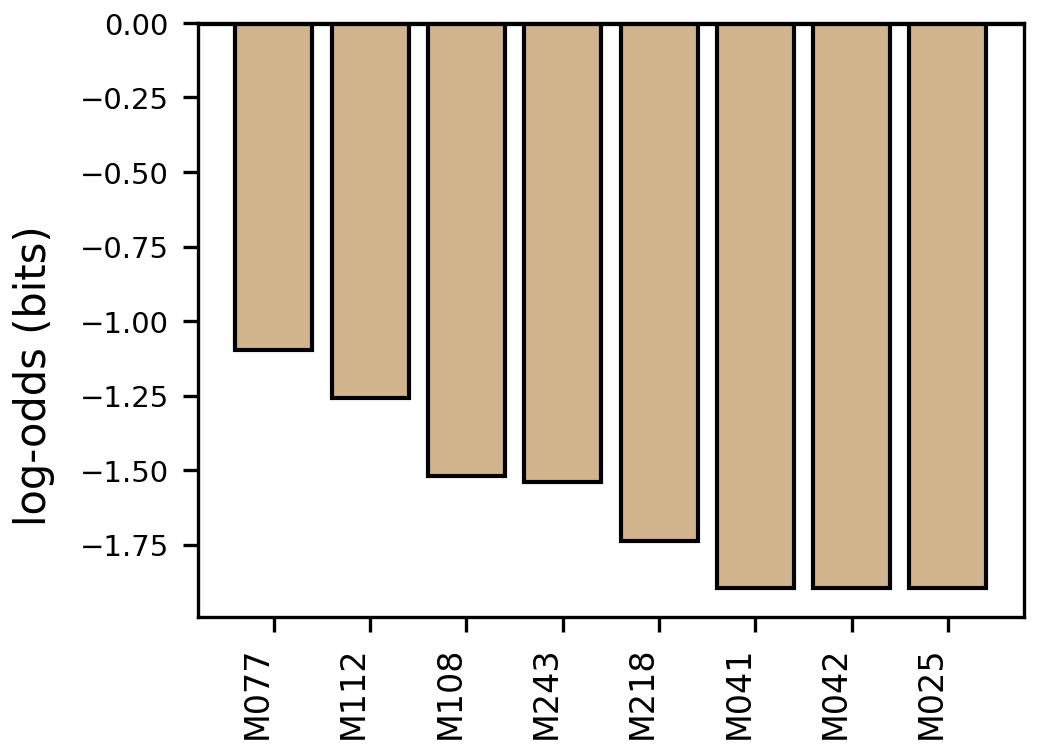

### motifs_turquoise_log_odds.png

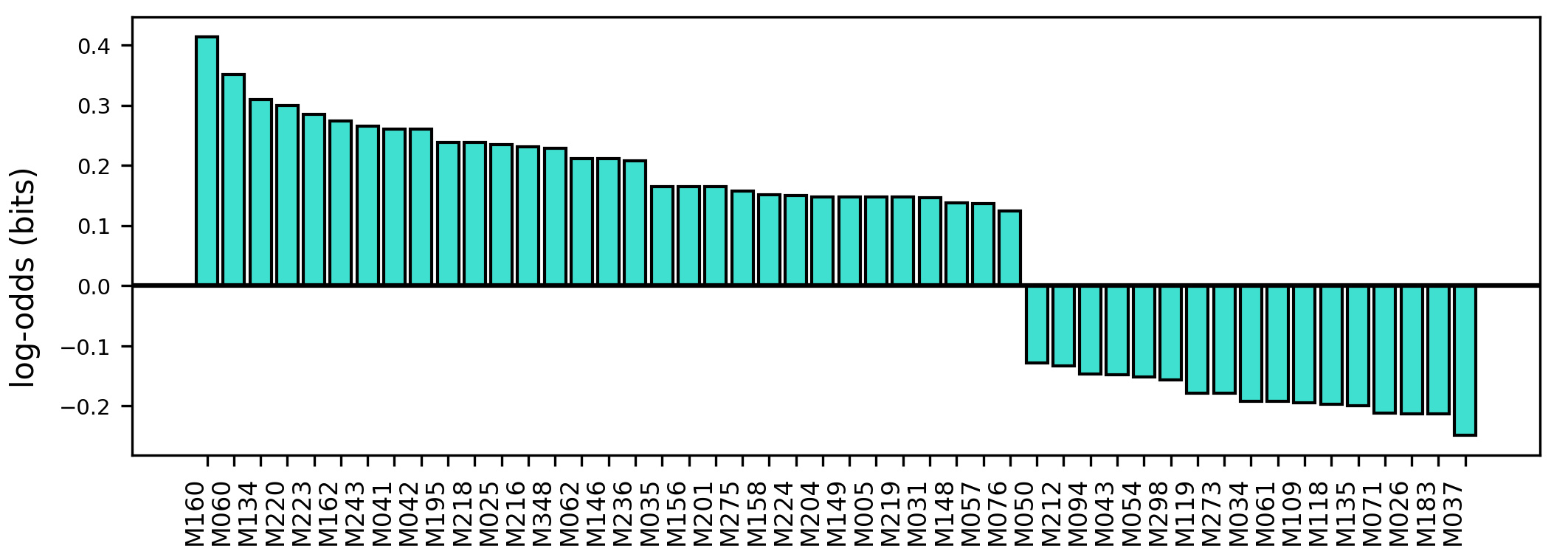

### org-ltximg_c34c51d1b8aa00e6bcafc05c8e3a419948093bd2.png

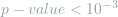

### part_of_log_odds.png

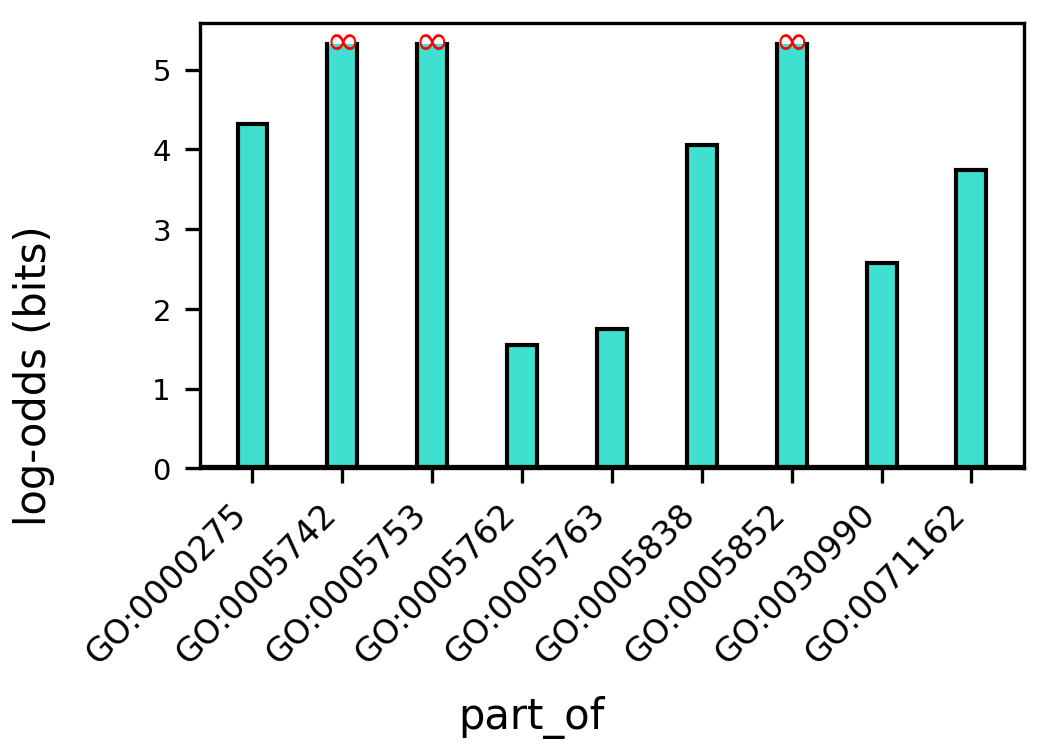

### poly-cistron_log_odds.png

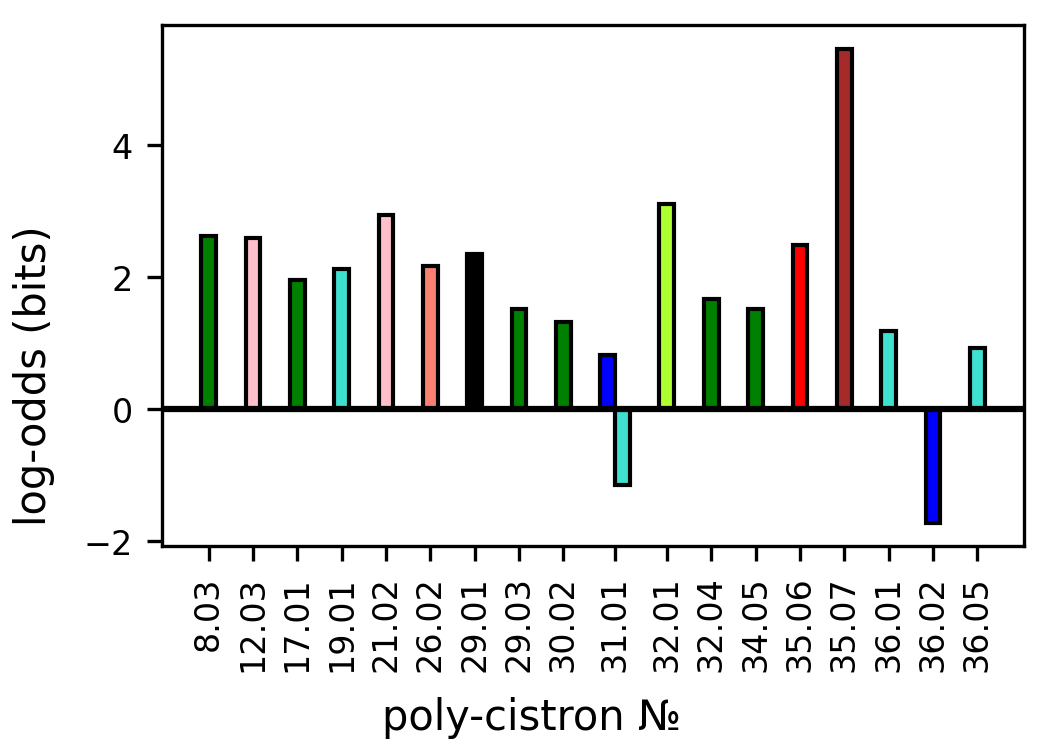

### terms_black_log_odds.png

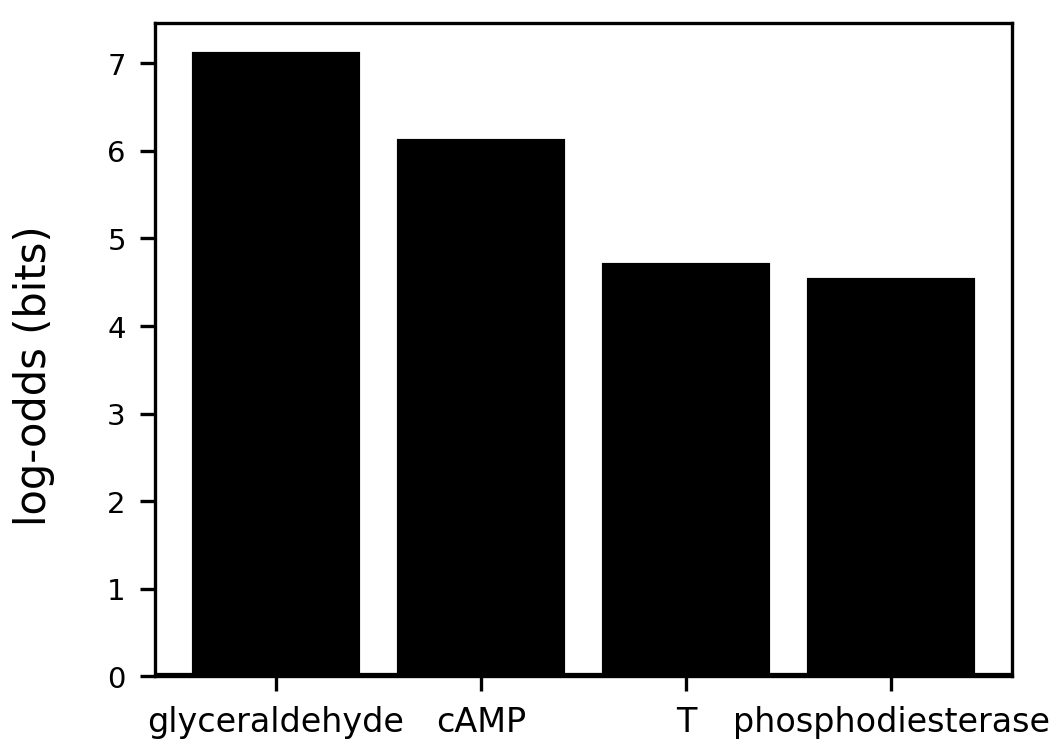

### terms_blue_log_odds.png

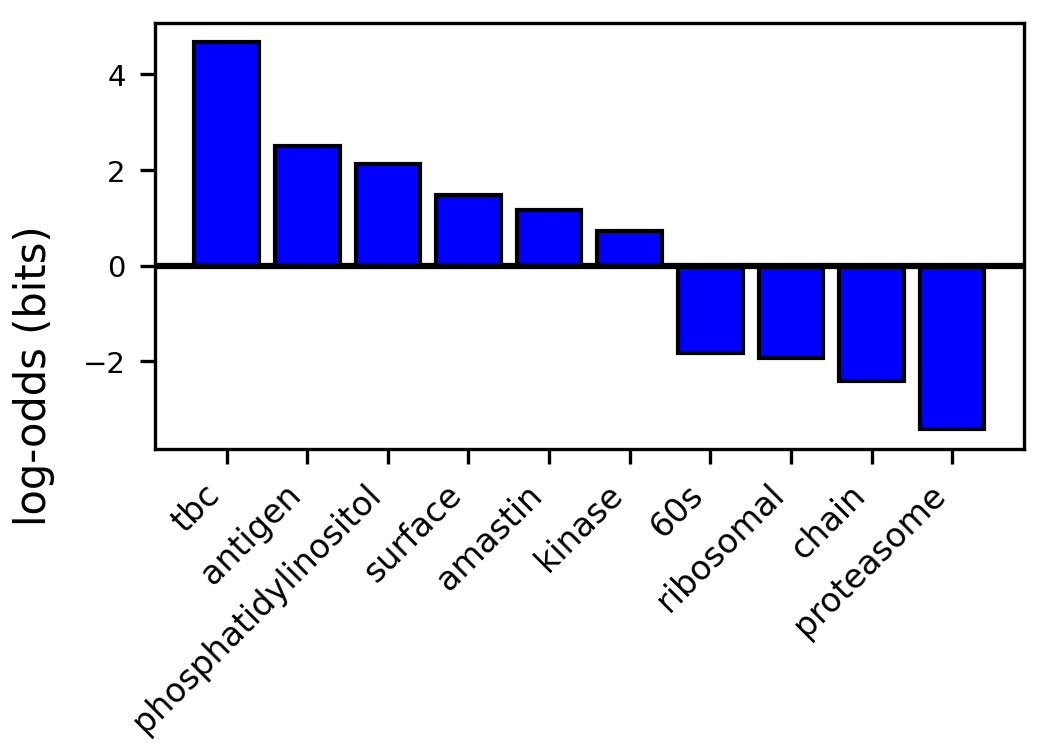

### terms_darkorange_log_odds.png

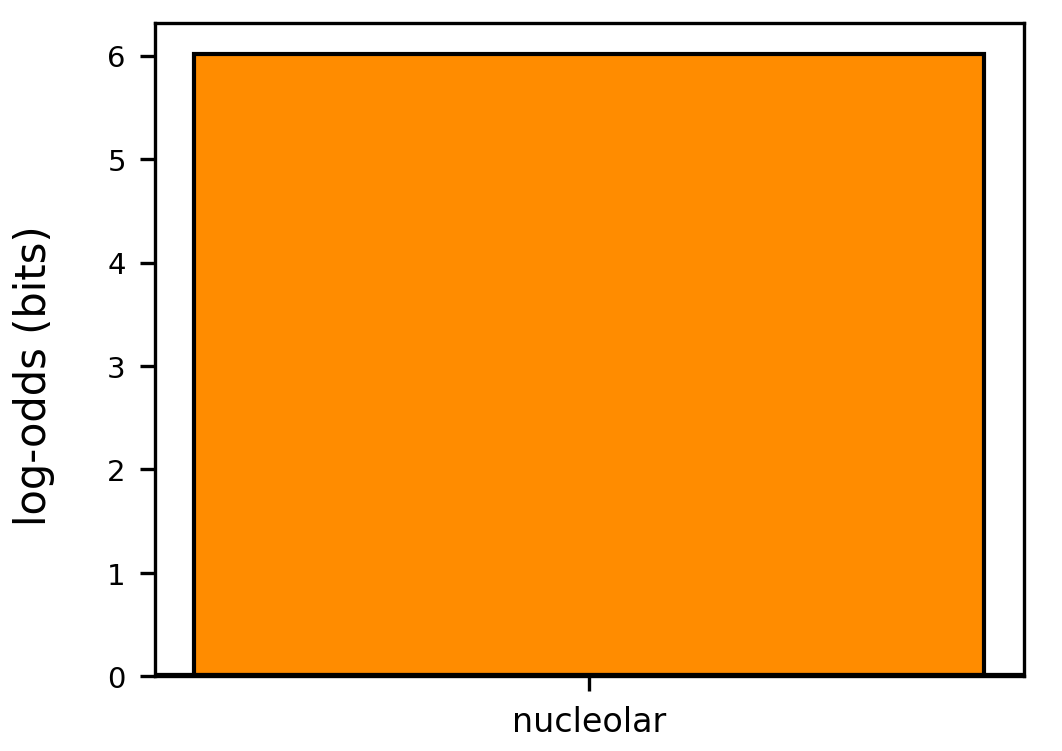

### terms_green_log_odds.png

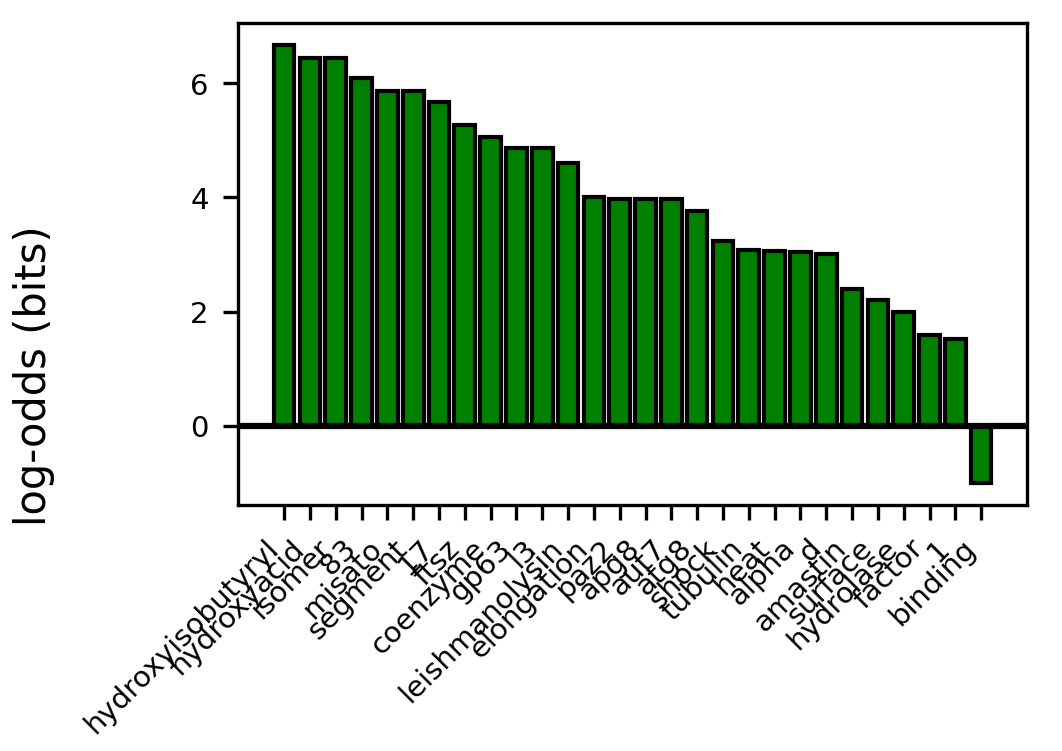

### terms_greenyellow_log_odds.png

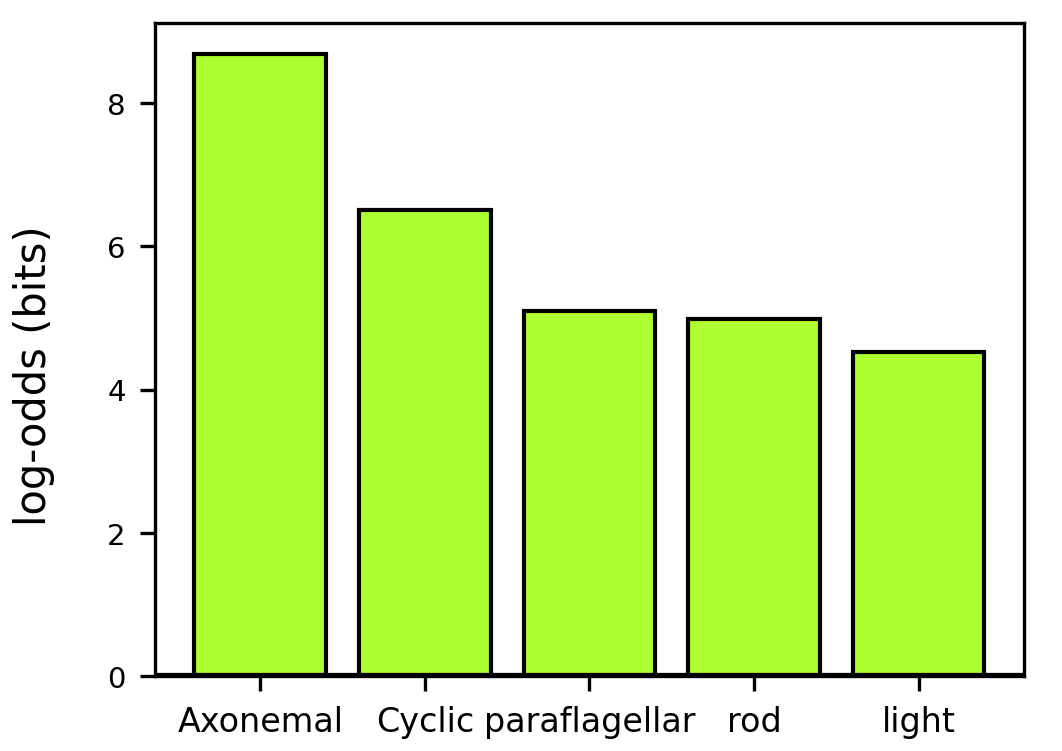

### terms_grey_log_odds.png

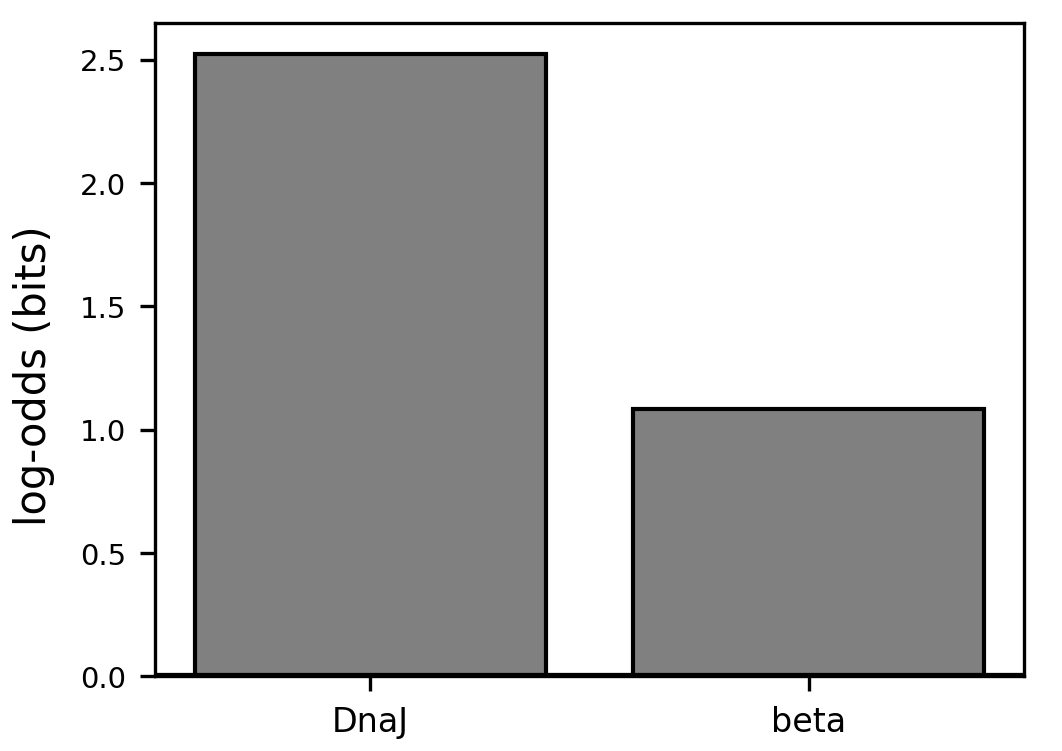

### terms_lightgreen_log_odds.png

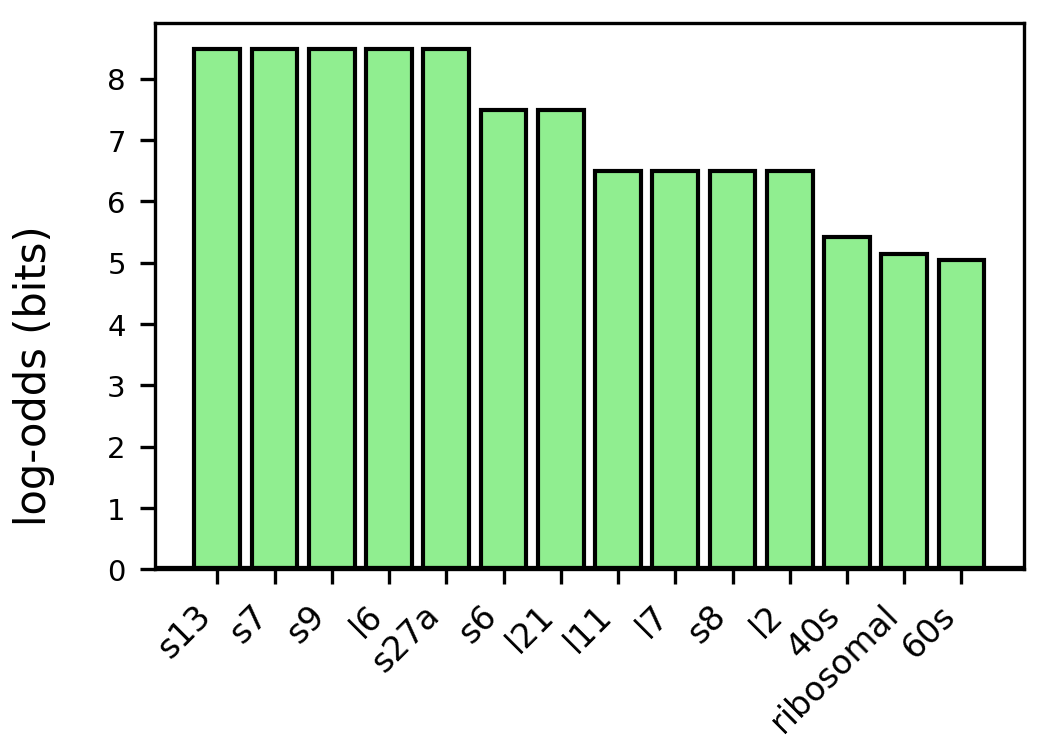

### terms_log_odds.png

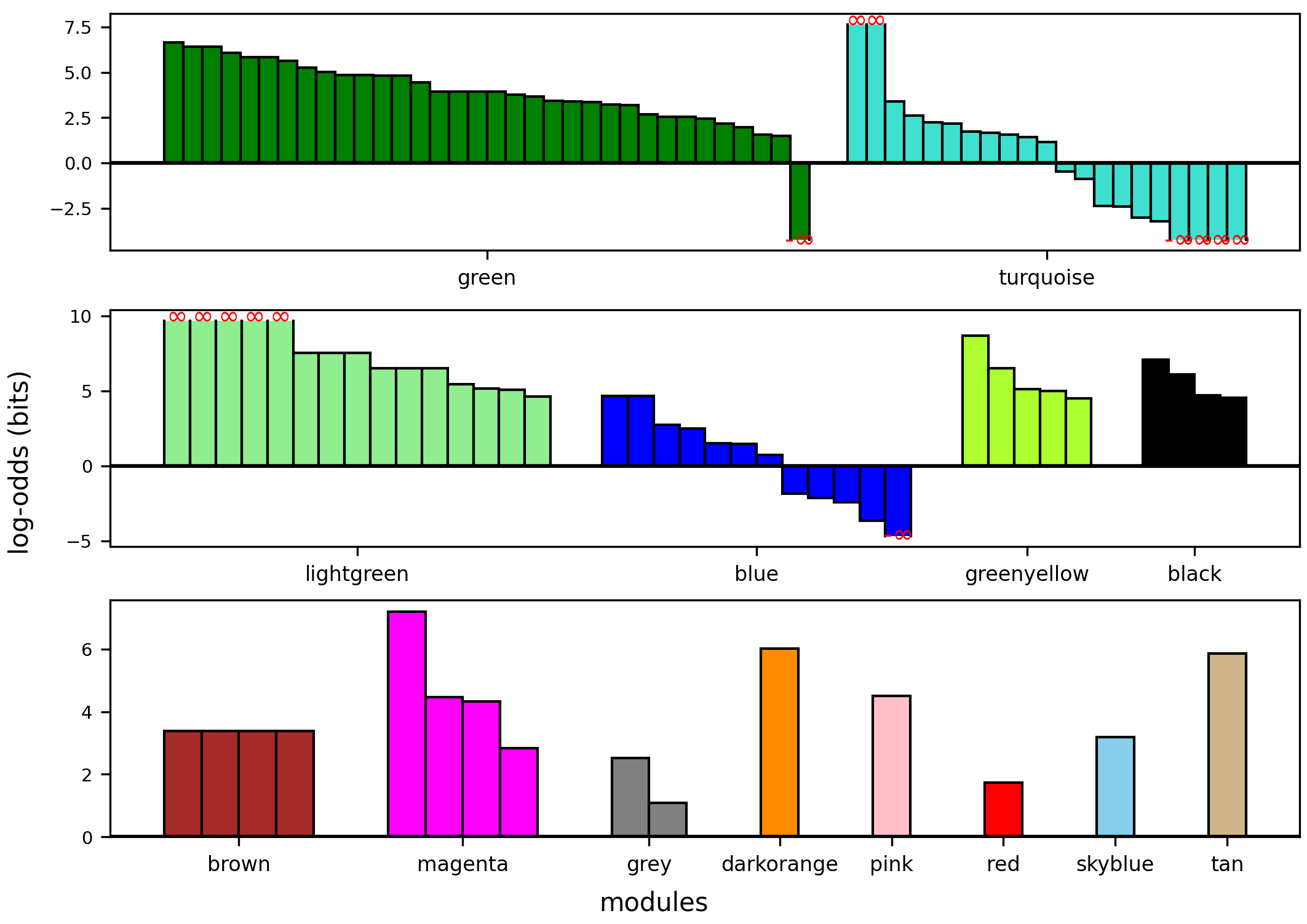

### terms_num.png

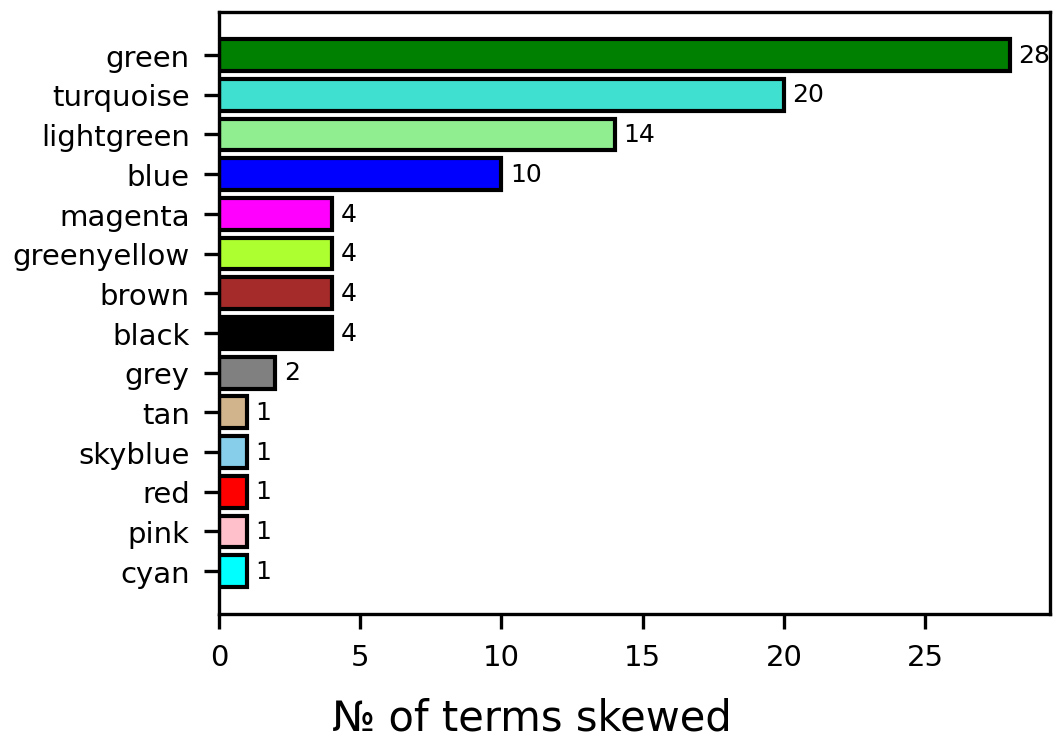

### terms_pink_log_odds.png

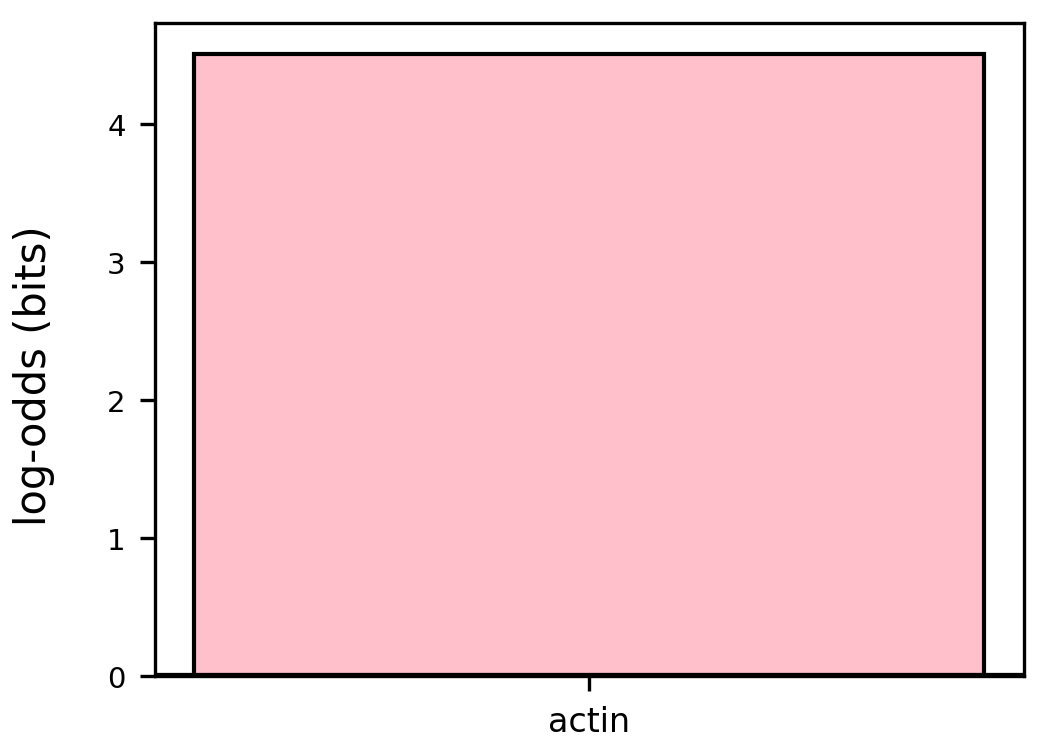

### terms_red_log_odds.png

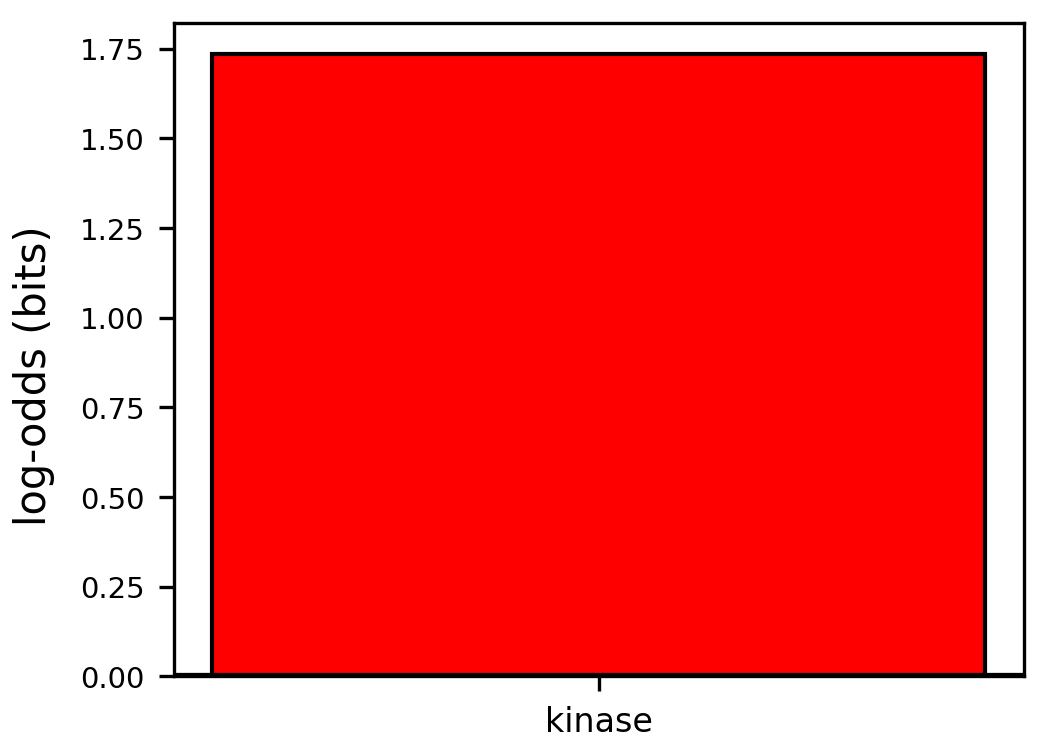

### terms_skyblue_log_odds.png

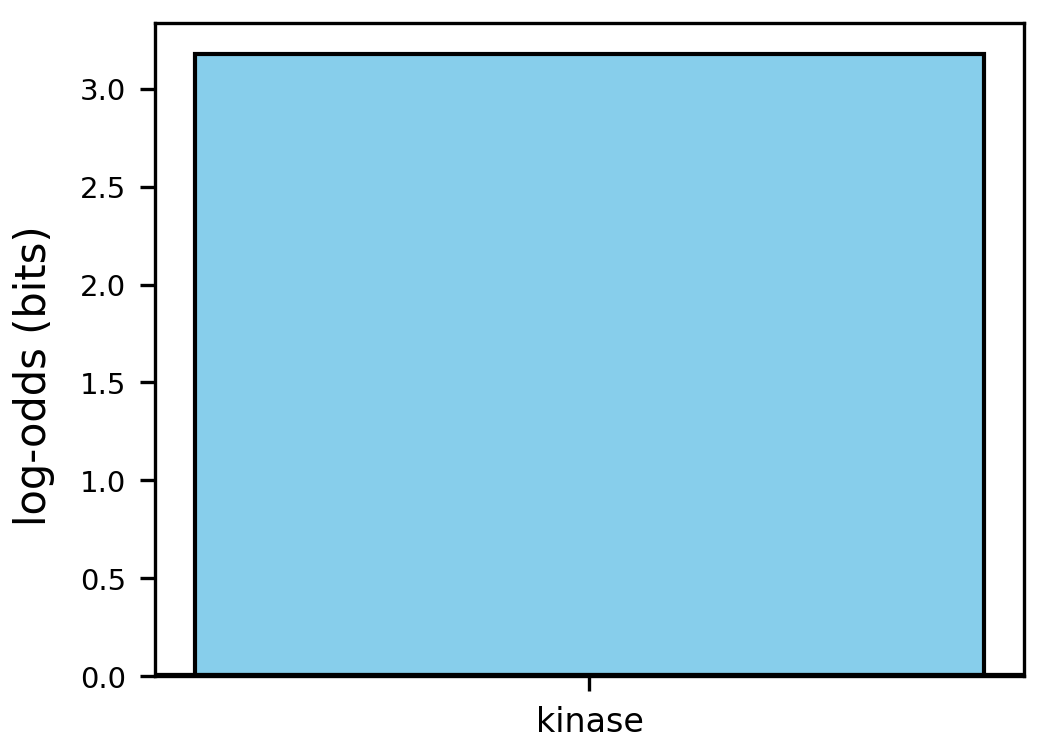

### terms_tan_log_odds.png

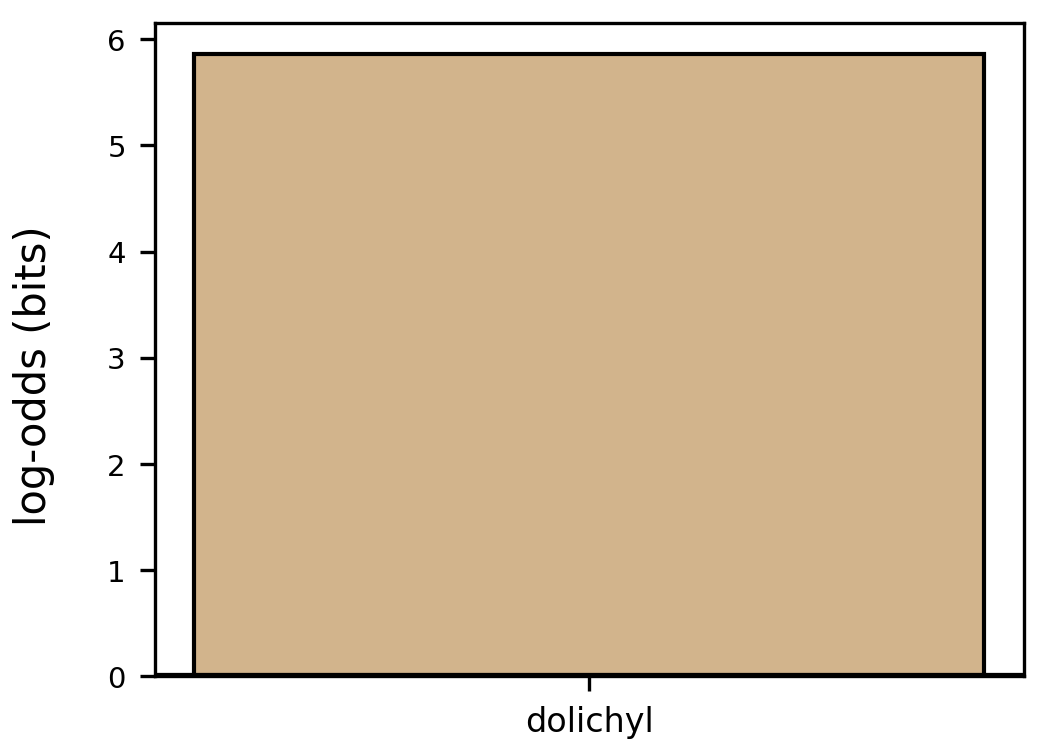

### terms_turquoise_log_odds.png

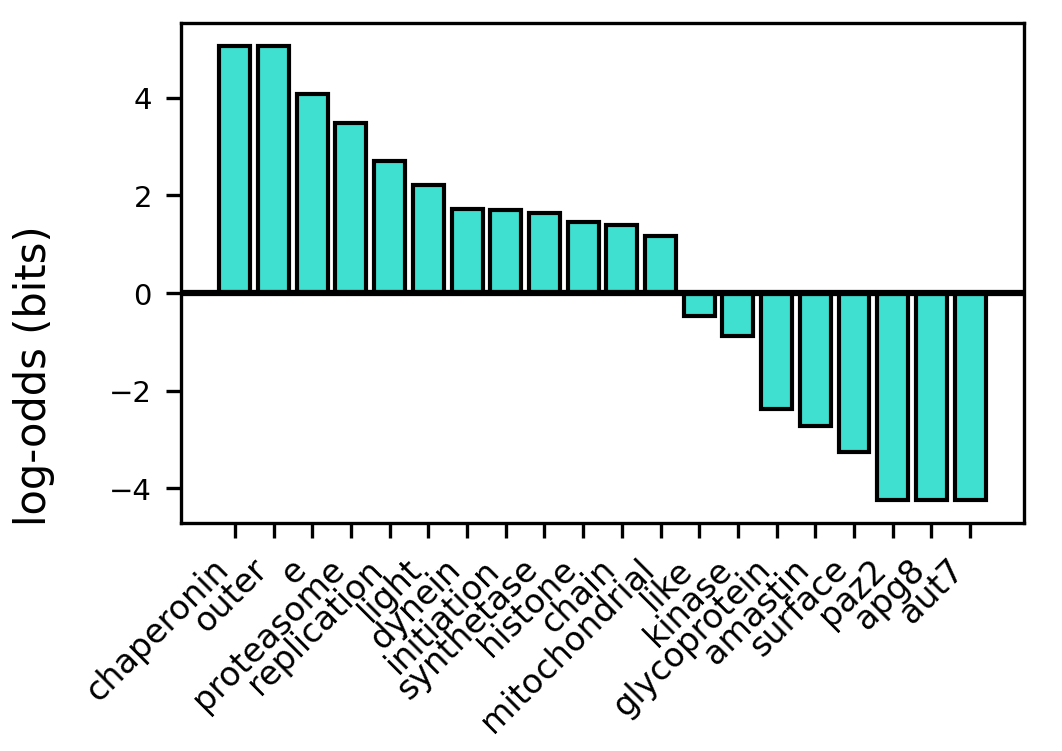
