## Supplementary File 1 for "Meta-analysis of the pathogen *Leishmania donovani*’s transcriptome reveals multiple modes of regulation including two reciprocally regulated gene modules"

### WGCNA clustering of pathogen *Leishmania donovani*'s transcriptome reveals multiple modes of regulation including two reciprocally regulated gene modules: Supplementary data

January 27, 2024

#### 1 Inputs

##### 1.1 Datasets raw reads accession

| Conditions | Accession |
|---|---|
| Amastigotes Mouse Liver | SRR10903037 |
| Amastigotes Mouse Liver | SRR10903038 |
| Amastigotes Mouse Liver | SRR10903039 |
| Amastigotes Mouse Liver | SRR10903040 |
| Amastigotes Mouse Liver | SRR10903041 |
| Amastigotes Mouse Spleen | SRR10903052 |
| Amastigotes Mouse Spleen | SRR10903053 |
| Amastigotes Mouse Spleen | SRR10903054 |
| Amastigotes Mouse Spleen | SRR10903055 |
| Amastigotes Mouse Spleen | SRR10903056 |
| Promastigotes Log Arginine Repleated | SRR2136265 |
| Promastigotes Log Arginine Repleated | SRR2136266 |
| Promastigotes Log Arginine Depleted | SRR2136267 |
| Promastigotes Log Arginine Depleted | SRR2136268 |
| Axenic Amastigotes Arginine Repleated | SRR2136269 |
| Axenic Amastigotes Arginine Repleated | SRR2136270 |
| Axenic Amastigotes Arginine Depleated | SRR2136271 |

Continued on next page

Continued from previous page

| Conditions | Accession |
|---|---|
| Axenic Amastigotes Arginine Depleated | SRR2136272 |
| Amastigotes Macrophage Human THP1 | SRR2136273 |
| Amastigotes Macrophage Human THP1 | SRR2136274 |
| Promastogotes Stat SL | SRR5272520 |
| Promastigotes Stat SL | SRR5272521 |
| Promastigotes Stat SL | SRR5272522 |
| Promastigotes Stat SL | SRR5272523 |
| Promastigotes Stat | SRR5272528 |
| Promastigotes Stat | SRR5272529 |
| Promastigotes Stat | SRR5272530 |
| Promastigotes Stat | SRR5272531 |
| Promastigotes Log SL | SRR5272524 |
| Promastigotes Log SL | SRR5272525 |
| Promastigotes Log SL | SRR5272526 |
| Promastigotes Log SL | SRR5272527 |
| Promastigotes Log | SRR5272532 |
| Promastigotes Log | SRR5272533 |
| Promastigotes Log | SRR5272534 |
| Promastigotes Log | SRR5272535 |
| Mouse Macrophage amastigotes SL | SRR5272514 |
| Mouse Macrophage amastigotes SL | SRR5272515 |
| Mouse Macrophage amastigotes SL | SRR5272516 |
| Mouse Macrophage amastigotes SL | SRR5272517 |
| **Conditions (dropped)** | **Accession** |
| Axenic Amastigotes | SRR924395 |
| Axenic Amastigotes Cutaneous | SRR924396 |
| Promastigotes Log | SRR924397 |
| Promastigotes Log Cutaneous | SRR924398 |
| Axenic Amastigotes Switch | SRR924399 |
| Axenic Switch Amastigotes Cutaneous | SRR924400 |
| Amastigotes Mouse Macrophage Passage 2 | SRR8479211 |
| Amastigotes Mouse Macrophage Passage 25 | SRR8479212 |
| Promastigotes SL | SRR5272501 |
| Promastigotes SL | SRR5272504 |
| Promastigotes SL | SRR5272507 |
| Amastigotes Macrophage Human | SRR5272508 |
| Amastigotes Macrophage Human | SRR5272509 |
| Amastigotes Macrophage Human | SRR5272510 |
| Amastigotes Macrophage Human | SRR5272511 |

Continued on next page

Continued from previous page

| Conditions | Accession |
|---|---|
| Promastigotes Purine Repleted | SRR923820 |
| Promastigotes Purine Depleted | SRR923821 |

Table 1: **Input data sets and accessions**. Experimental runs in the lower part of the table were dropped for lack of replicates or due to quality control.

#### 2 Batch-adjusted dendrogram

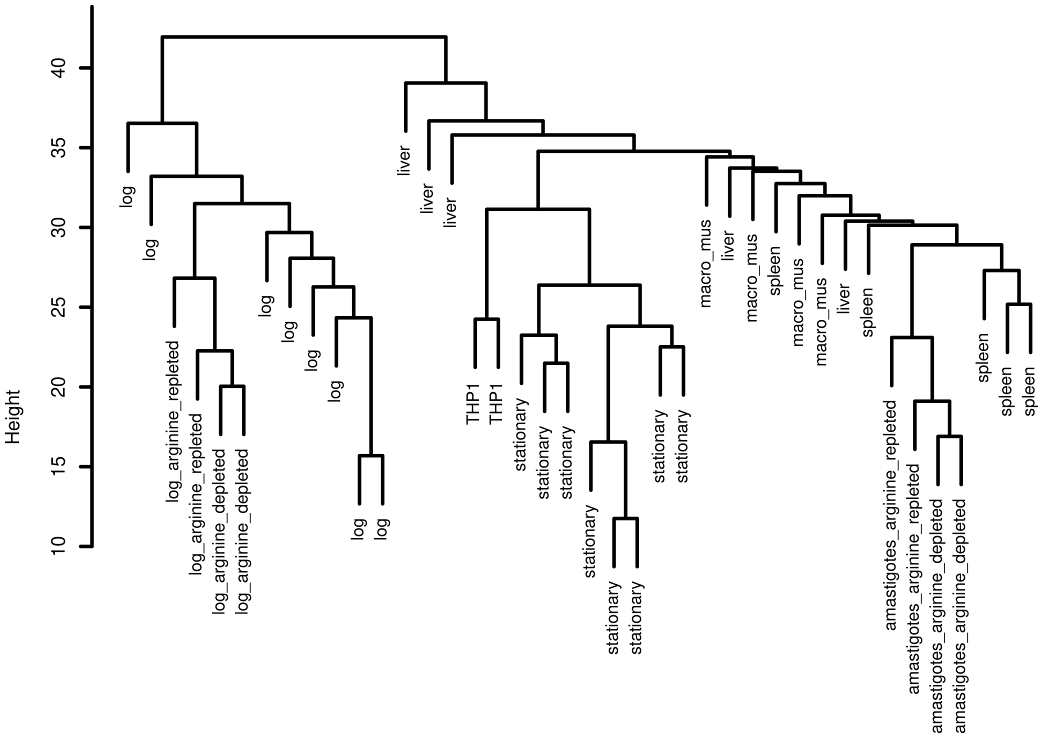


Figure 1: Clustering of conditions with read-counts adjusted for batch-effects.

#### 3 Validation of WGCNA Modules

##### 3.1 Annotation terms

A test of validity of WGCNA of *Leishmania* transcriptome would be that the resultant modules of co-regulated genes would be enriched for genes of related regulation, function, pathways, etc.

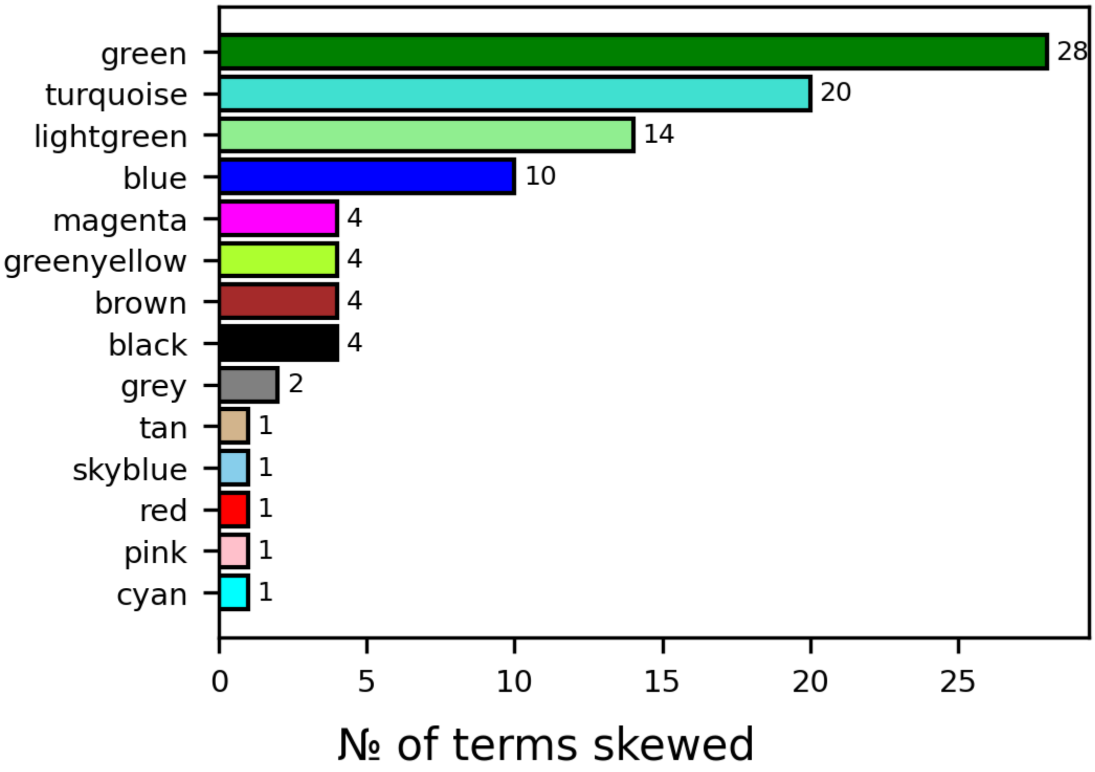


Figure 2: Non-random assortment of annotation terms. Color of bar indicates module. $[p - value < 10^{-3}]$

Typically, researchers describe genes of their interest submitted by them to databases using terms describing structural, regulatory or functional characteristics of the concerned genes. Based upon such manual annotations and sequence similarity, the process is facilitated by automated gene-annotation.

We pooled all informative terms used for annotation of LdHU3 genes, *i.e.* case-insensitive words in annotated names, after filtering for uninformative terms such as `hypothetical, conserved, unknown`, which either do not pertain to regulation and function, and/or are likely to be added during automated annotation. Supplementary table (2) is an exhaustive list of terms deemed *uninformative*. For each term, occurrence of genes qualified using that term in each WGCNA module was counted to generate a contingency table of terms and modules. Under a null hypothesis of independence of module identity and term-usage, $\chi^2$ test was performed for the contingency table. Usage of terms was found to be inter-dependent with identity of modules ($\chi^2$ test $p - value = 2.343 \times 10^{-06}$), indicating non-random clustering.

We further checked association between a given WGCNA module and occurrence of genes annotated with each term using *Fisher's exact* test, ( $p - value < 10^{-3}$). *Fisher's exact* test identifies uneven partitioning of elements from the available collective pool, hence, accounting for excess of scarce occurrence of any specific term. Figure (2) represents number of terms that are associated with each module. We describe here in detail, individual modules in which, genes with at least 5 terms are enriched. Supplementary files contain a complete list of modules and terms that describe genes enriched in each module.

The *green* module (Figure 3) is enriched with genes described by 27 terms including `hydroxyisobutyryl, hydroxyacid, leishmanolycin, gp63`, while being depleted with genes whose annotation includes the term `binding.`

Ribosome-associated terms such as `ribosomal`, `60s`, `40s`, `s13`, `s7` are enriched in the *light-green* module (Figure 4).

*Blue* module (Figure 5) is enriched with genes described with the terms `tbc, antigen, phosphatidylinositol, surface` and `kinase` while being depleted for genes described by the terms `60s, ribosomal, proteasome`.

*Turquoise* (Figure 6) module is enriched with genes described by 12 terms including `chaperonin, outrt, proteasome, replication` while being depleted from terms described by 8 terms including `aut7, apg8, paz2, suface`.

In addition to supporting apt classification by WGCNA, identity of terms [NON RANDOMLY: PRADY OPINION=TAUTOLOGY] skewed in each module further extends information about other genes in that module.

##### 3.1.1 Terms deemed uninformative

| Ignored terms |
|---|
| protein |

Continued on next page

| Continued from previous page |
|---|
| Ignored terms |
| conserved |
| hypothetical |
| putative |
| subunit |
| domain |
| containing |
| unknown |
| uncharacterized |
| uncharacterised |

Table 2: Terms that occur in automated annotation, without adding information.

#### 3.2 GO-Qualifiers

| GO Qualifier | p-value |
|---|---|
| located_in | 1.93e-43 |
| aspect | 3.74e-125 |
| acts_upstream_of_or_within | 5.76e-18 |
| part_of | 8.00e-205 |
| enables | 0.426 |

Table 3: GO: qualifier

Gene Ontology project functionally qualifies genes from organisms based on current bio-informatics knowledge. With an expectation that genes involved in similar functional paradigms should be co-regulated, we checked if any of the GO-Qualifiers is unevenly distributed amongst WGCNA modules. GO-identifiers for *Leishmania major* orthologs that formed basis for automated annotation of LdHU3 transcripts were identified from GO-annotations file (gaf).

For each available qualifier category (3), occurrence of genes bearing GO-terms was counted against each WGCNA module. Table (3) indicates $\chi^2$ p-values for inter-dependence of WGCNA-classification and GO-Qualifier. Barring the qualifier "enables", all other qualifiers show strong dependence with WGCNA modules confirming their {validity,correctness}.

Table (4), enlists association of indicated WGCNA modules with GO-terms used to qualify the GO-Qualifier `located in` (*Fisher's exact test*, p-value <0.001). Positive and negative log-odds respectively indicate enrichment and depletion of the corresponding GO-terms, as log-odds (bits) of 1 indicate 2 fold occurrence over random expectation. Log odds indicated by `inf` denote exhaustive inclusion of *all* genes with that GO qualifier term in the corresponding module where as those indicated by `-inf` denote complete exclusion of *all* genes with that GO qualifier term from the corrsponding module.

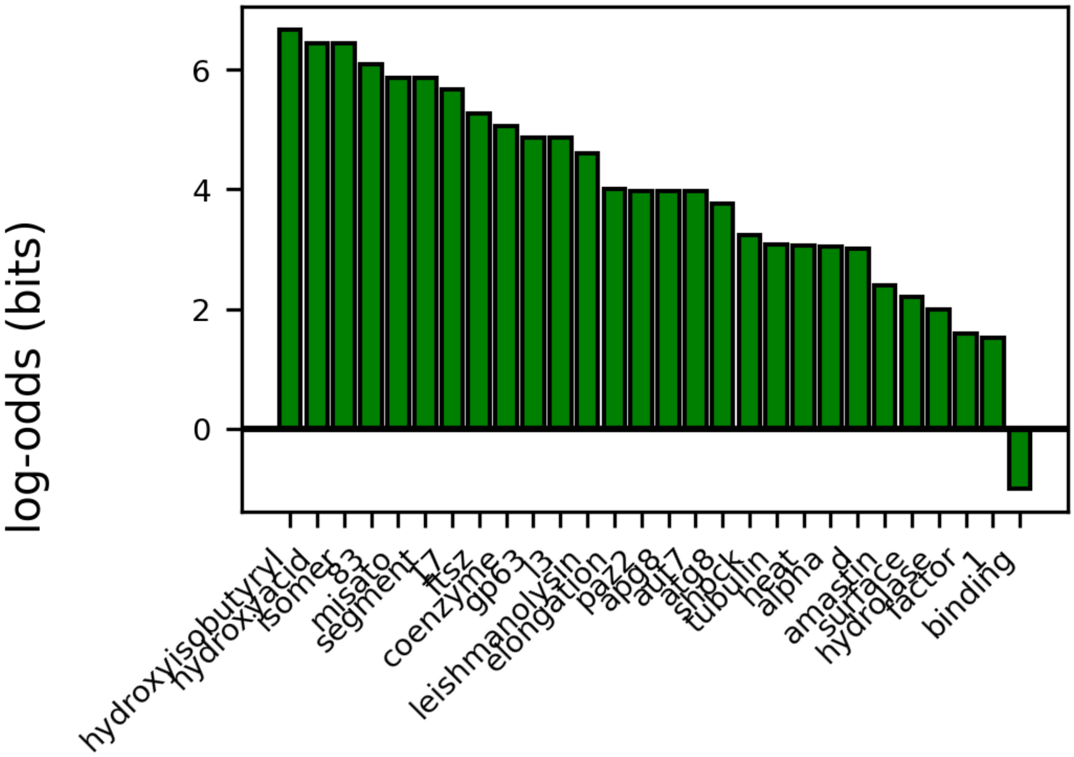


Figure 3: Skewed occurrence of annotation terms in *green* module

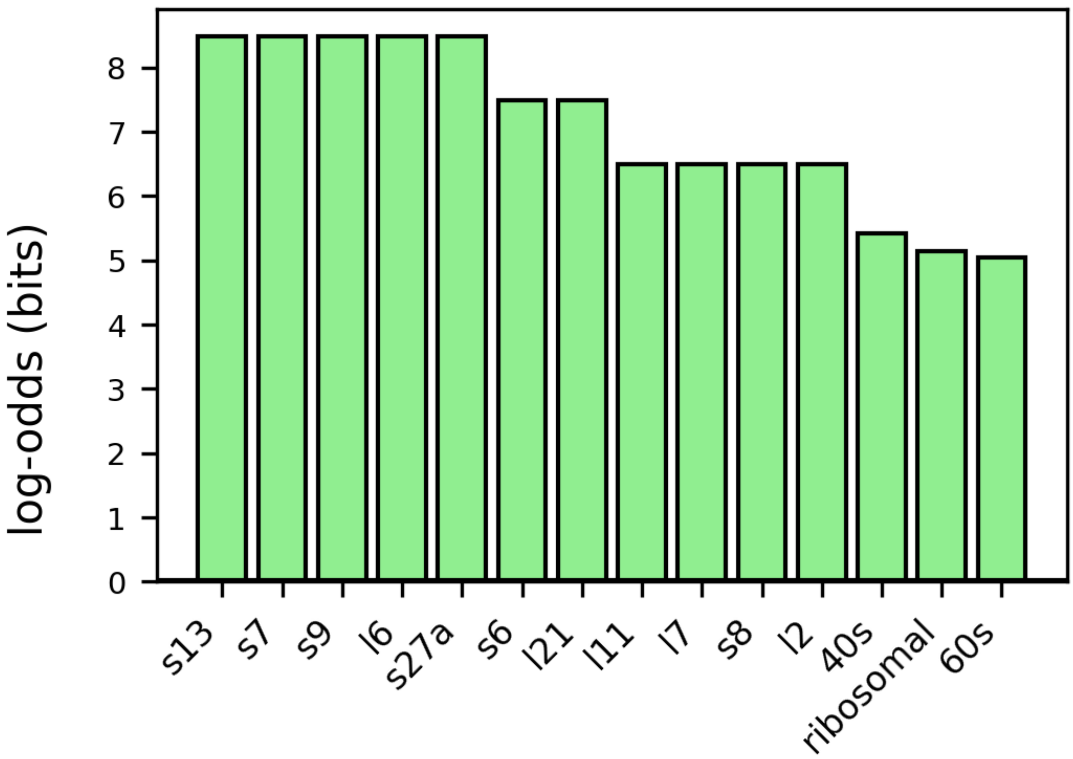


Figure 4: Skewed occurrence of annotation terms in *light-green* module

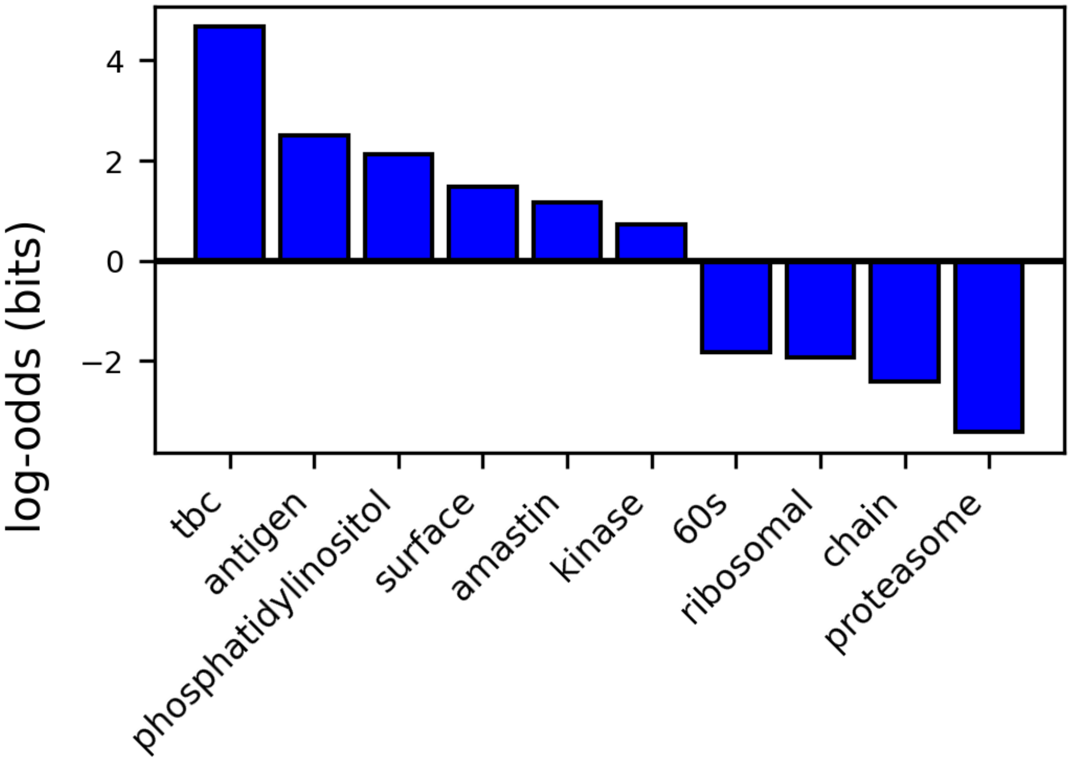


Figure 5: Skewed occurrence of annotation terms in *blue* module

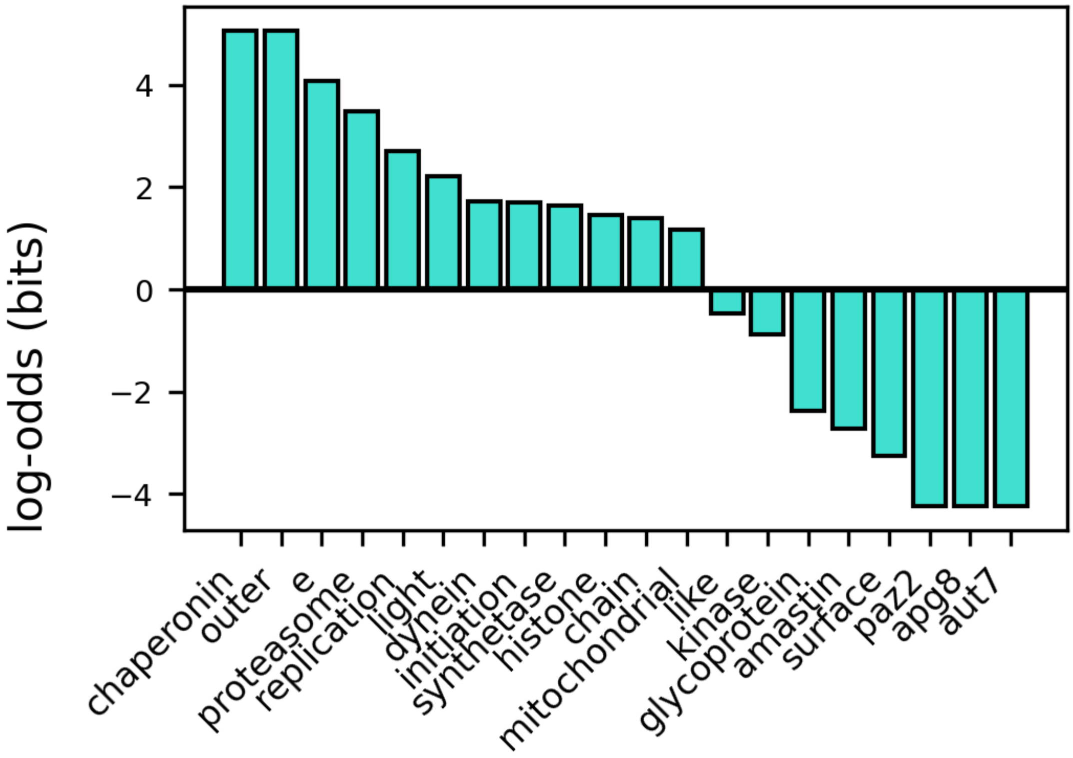


Figure 6: Skewed occurrence of annotation terms in *turquoise* module

| Modules | Located in | log-odds (bits) |
|---|---|---|
| turquoise | cortical cytoskeleton | inf |
| | mitochondrial outer membrane | 1.8210 |
| | glycosome | 1.5060 |
| | axoneme | 1.4189 |
| | ciliary basal body | 1.1359 |
| | mitochondrion | 0.9978 |
| | nucleoplasm | 0.9462 |
| | kinetoplast | 0.9115 |
| | ciliary plasm | 0.8104 |
| | cytoplasm | 0.5091 |
| blue | cytoplasm | -0.4268 |
| | cilium | -1.8424 |
| | axoneme | -3.3892 |
| green | cytoplasm | -1.4922 |
| | mitochondrion | -3.5458 |
| lightgreen | nucleolus | 2.7189 |
| | cytoplasm | 2.417 |
| yellow | endoplasmic reticulum | 1.6351 |
| | axoneme | -inf |
| cyan | nucleolus | 2.5148 |
| darkorange | nucleolus | 3.2695 |
| greenyellow | axoneme | 3.8119 |
| grey60 | axoneme | 4.0042 |

Table 4: GO: Located in

#### 3.3 Motifs

##### 3.3.1 Uncertain Motifs

| Motif | p-value |
|---|---|
| M033 | 0.2206 |
| M036 | 0.1153 |
| M080 | 0.0291 |
| M107 | 0.0291 |
| M137 | 0.0015 |
| M176 | 0.0220 |
| M205 | 0.7624 |
| M245 | 1.0 |
| M260 | 0.0124 |
| M274 | 0.0161 |
| M287 | 0.6400 |
| M291 | 0.8236 |
| M299 | 0.0183 |
| M300 | 0.0183 |
| M317 | 0.7819 |
| M323 | 0.0010 |
| M334 | 1.0 |
| M346 | 0.1621 |
| M350 | 0.0263 |

Table 5: Motifs without certainty about skewed occurrence in *Leishmania*. $p - value > 0.001$
